# From Wikipedia to AI: Measuring 25 years of synthesis of human genetics research in the public-facing information ecosystem

**DOI:** 10.64898/2026.07.27.741055

**Authors:** Alex Diaz-Papkovich, Abigail Kuntzleman, Sophia C. Davis, Sohini Ramachandran

**Author notes:** Corresponding authors. Email: alex (A.D.P.); (S.R.).

## Abstract

Genetics research frequently intersects with ethnicity, nationality, and race, making it uniquely vulnerable to misrepresentation. Yet, 25 years after the initial sequencing of the human genome, there is little understanding of how human genetics research exists in the public-facing information ecosystem. We analyze 3,050,422 historical revisions from 6,738 Wikipedia pages about ethnicity, nationality, and race spanning 25 years. We find genetics terminology is present in 14.8% of these pages (55.5% in the top 1,000 pages) and in 67.8% of pages about nationalities, suggesting research is synthesized to present a biological element to ethnicity and nationality. We also find that 10.1% of 56,908 discussions from these pages contain genetics terminology. We further analyze responses from three popular chatbots queried about nationalities and find that they commonly reference both genetics and Wikipedia. Lastly, we analyze 133 pages from Grokipedia, an AI-generated encyclopedia, and find it mentions genetics more frequently than Wikipedia and hallucinates or misrepresents human genetics research.

---

Spurred by the technology that eventually enabled the publication of the human genome draft sequence in 2003, millions of human genomes have been analyzed since the late 20th century. This body of research—which we refer to as “human genetics research”—has yielded new insights into human evolution (e.g. (*1*)), genetic determinants of health and disease (e.g. (*2*)), and human population histories (e.g. (*3*)), all while becoming increasingly accessible through a vast and expanding landscape of peer-reviewed publications.

Because human genetics research frequently intersects with race and ethnicity, it is uniquely vulnerable to misunderstanding by the public and misappropriation by extremists (*4–7*). In response to these concerns, professional scientific organizations such as the American Society of Human Genetics (ASHG) and the National Academies of Sciences, Engineering, and Medicine (NASEM) have issued recommendations and technical guidance on how to avoid the conflation of genetics with social labels such as ethnicity or race (*8, 9*). Beyond these conversations within the scientific community, researchers are increasingly studying how human genetics research is interpreted, remixed, and sometimes co-opted by various communities to serve ideological ends (*5, 10–12*). However, the vast majority of Americans do not interface with human genetics research through peer-reviewed papers or conference presentations, but rather through media such as the news, social media, or websites (*13, 14*). To date, we do not have an understanding of how human genetics research lives in the public-facing information ecosystem and, in particular, we do not know how a typical person will encounter human genetics research online in the context of ethnicity, nationality, or race.

One of the world’s most popular websites is Wikipedia (*15*), an online encyclopedia. Its English language edition received over 264 billion page views from 940 million unique devices between January 2025 and July 2026 (*16*), and it regularly appears at the top of search engine results and in “knowledge panel” summaries (*17*). It is one of the most-trusted sources of information on the internet (*18*), and its contents are influential contributors to training data for modern AI systems like large language models (LLMs) (*19*). These factors make Wikipedia a cornerstone of today’s online information ecosystem. Further, readers are encouraged to write and edit comprehensive pages with text and media that can be verified through published reliable sources, such as news reports and scientific manuscripts; Wikipedia contributions effectively combine to synthesize entire fields of research and make these fields accessible to the public.

Here, we use 25 years of Wikipedia data for a metascientific analysis of how human genetics research is synthesized in the context of ethnicity, nationality, and race. We analyze 6,738 Wikipedia pages about ethnicity, nationality, and race, as well as their 3,050,422 historical revisions spanning from January 2001 to December 31, 2025. We identify keywords related to genetics in these pages, as well as the contexts in which the keywords appear, their supporting citations and figures, whether a page has a dedicated section to genetics, topics covered when genetics terms are present, and trends in the representations of human genetics research. We also mine 56,098 user discussions associated with pages about ethnicity, nationality, and race and measure how often genetics is mentioned. Using pageview data, we estimate the public-facing impact of human genetics research when presented in the context of ethnicity, nationality, and race. Across 970,644 paragraphs in Wikipedia pages and 730,560 paragraphs in user discussions, we identify where genetics keywords appear and analyze 14 common topics such as “ancestry & origins” and “identity & ethnicity”, and provide examples of how human genetics research is synthesized and presented. We also highlight three pages as case studies (“Ashkenazi Jews”, “Black people”, and “White people”) to trace the influence of hereditarian writers, and use link click data to estimate reader interest in genetics.

To further highlight the public-facing impact of the intertwining of human genetics research with ethnicity, nationality, and race, we also query three popular LLMs (ChatGPT, Claude, Gemini) about nationalities, and measure both whether they mention genetics and whether they cite Wikipedia. Lastly, we analyze Grokipedia, an AI-generated encyclopedia created by Elon Musk’s company xAI. Here, we compare the use of genetics in writing between Grokipedia pages and their Wikipedia counterparts, using 130 pages about nationalities plus the three case studies. Taken together, we find the synthesis of human genetics research on Wikipedia in the context of ethnicity, nationality and race is inadvertently framed to support “genetic essentialism”—the misconception that complex social and political identities are fixed biological realities simplistically determined by DNA.

## RESULTS

We examine how human genetics research is packaged for public consumption. We analyzed 6,738 Wikipedia pages and their 3,050,422 historical revisions. Of the 6,738 pages, 6,541 consist of pages about extant ethnic groups, nationalities, and racial groups (hereafter referred to as the “corpus”) and their 2,864,806 revisions; the remaining 197 pages are national-level “Demographics of” pages (e.g. “Demographics of Algeria”) and their 185,616 revisions, referred to as “demographics pages”. Demographics pages, while not strictly about ethnic groups, sometimes contained text related to human genetics research that was eventually spun off into pages about, e.g., nationalities. Revisions on the English language Wikipedia can be accessed via URL using unique integer IDs (see Methods—Data collection). When discussing specific revisions, we cite the ID in parentheses e.g. “(revision 12345)”. The cut-off date for analysis of Wikipedia pages was December 31, 2025. The earliest revision in our dataset was March 28, 2001.

To identify pages for the corpus of 6,541 pages, we recursively explored Wikipedia’s page categorization system (see Methods—Data collection). These pages include those about ethnoreligious groups (e.g. “Amish”), nation-level demonyms (e.g. “Japanese people”), indigenous groups (e.g. “Māori”), ethnolinguistic groups (e.g. “Punjabis”), and trans-national ethnic groups (e.g. “Romani”). The corpus additionally includes diaspora groups (e.g. “Chinese Canadians”), census classifications (e.g. “White British”), and pages otherwise categorized by Wikipedia editors as ethnic groups (e.g. “Lithuanian minority in Poland”). Unless otherwise stated, all statistics we report about Wikipedia pages refer to pages within the corpus.

### Genetics content in corpus pages becoming more prevalent over time

The earliest appearance of genetics keywords in the corpus dates to October 30, 2001, when the page “Ainu people” referenced—without citation—that “recent genetic and morphological studies” had reported similarities between the Ainu, American indigenous peoples, and Japanese samurai (revision 453193519). At the end of 2001, 47 pages from the corpus had been created, and “Ainu” remained the only one with genetics keywords. Wikipedia then experienced rapid growth, with its millionth page being created March 1, 2006. The number of corpus pages increased accordingly, and the proportion of corpus pages containing genetics keywords also increased each year almost monotonically (figure 1A). By our analysis cut-off date, 970 of 6,541 (14.8%) pages contained genetics keywords.

**Figure 1:**
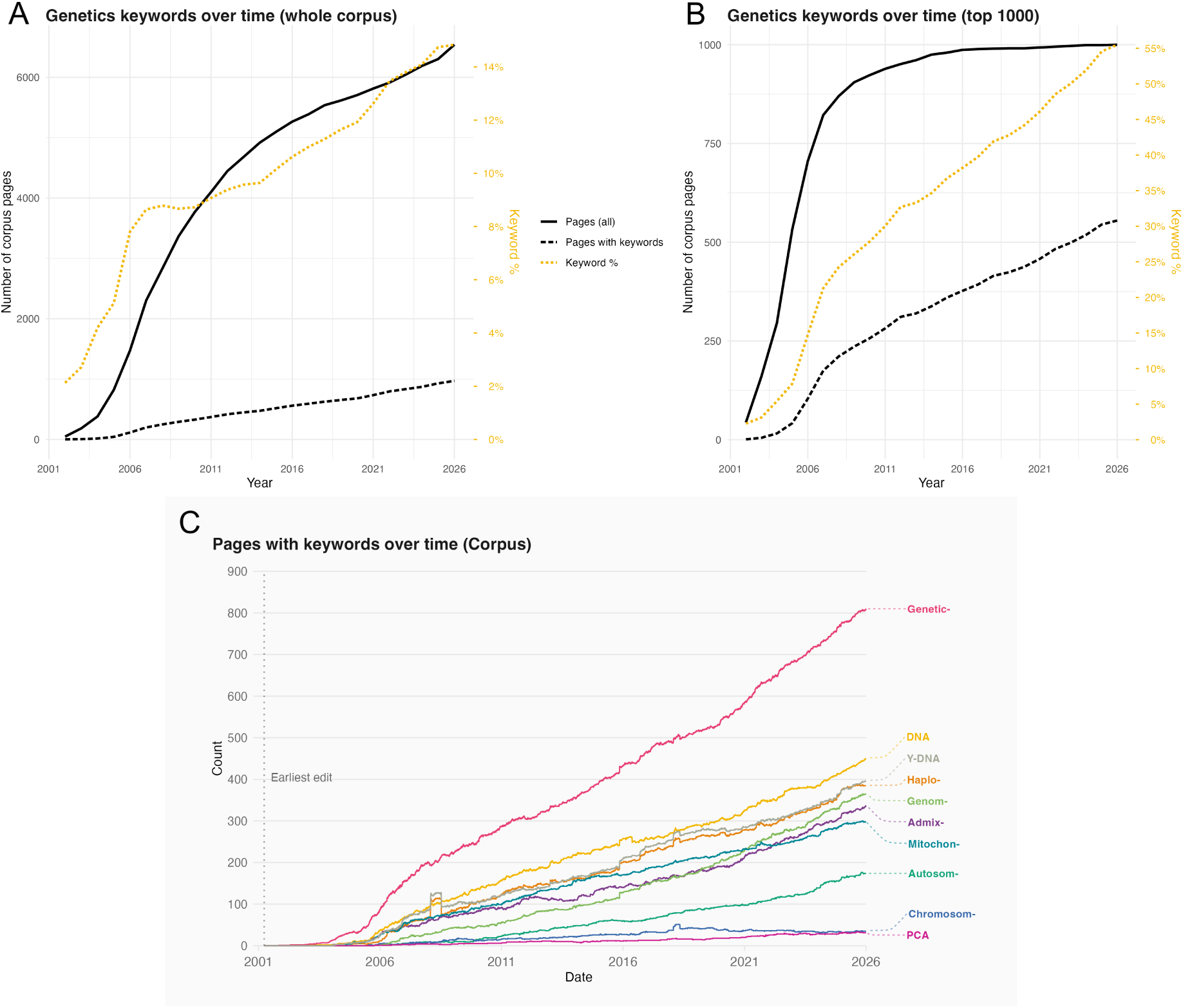
There has been a steady increase over 25 years in the use of genetics keywords in our Wikipedia corpus. **(A,B)**. The number of pages in our Wikipedia corpus that contained genetics keywords, from December 31, 2001 to December 31, 2025. The solid black line measures the number of pages in our corpus created at that point, and the dashed black line represents indicates the number of those pages that contained genetics keywords. The dotted marigold line indicates the percent of existing corpus pages at that time that contained genetics keywords (y-axis on right, in marigold). **(A)** All 6,541 corpus pages. **(B)** A subset of the top 1,000 most-viewed pages in the corpus for 2025. **(C)** The number of pages containing specific keywords across our Wikipedia corpus at the end of every day from March 28, 2001 until December 31, 2025.

Attention to Wikipedia pages, from both contributors and readers, is not uniform. Many pages are short and receive few views and edits, e.g. the page “Kango people” has been edited 17 times since its creation in 2011, was visited 1,170 times in 2025, and contains 795 characters of text, including spaces. Though there are 2,864,806 revisions in the corpus, the median number of revisions to a single page is 107 (figure S1). Similarly, corpus pages were viewed a collective 239,019,588 times in 2025, though the median page received 5,608 views, and pageviews followed an approximately log-normal distribution (fitted parameters *μ* = 8.800, *σ* = 1.829, figure S2). Genetics keywords are present in 555 (55.5%) of the top 1,000 most-viewed pages in 2025 (figure 1B). A logistic regression of keyword presence in all pages against log of the number of daily page views showed pageviews are a significant predictor of the presence of genetics keywords (*P* > |*z*| ≈ 0, OR = 2.58, pseudo *R*-squared = 0.295); that is, the most popular pages in the corpus are far more likely to have genetics keywords than less-viewed pages.

The most common keywords to appear across all corpus pages were “genetic” (appearing in 810 pages; 12.4%), “DNA” (521; 8.0%), “haplo” (385; 5.9%), “gene(s)” (385; 5.9%), and “genom” (365; 5.6%). By number of appearances overall, the most common keywords were “genetic” (appearing 6,709 times), “haplo” (3,979), “DNA” (3,181), “genom” (1,430), and “admix” (1,261) (figure 1C).

### Genetics content in corpus pages frequently organized into separate sections

Wikipedia pages are organized into sections, and section headings are prominently displayed, appearing as navigable links in the table of contents on the website and mobile app. We searched for genetics keywords in section titles; if keywords were present, we identified the section as a “genetics section”. Example genetics section titles include “Autosomal DNA” and “Genetic studies”. Pages can have multiple genetics sections (e.g. the page “Bulgarians” had a subsection titled “Bulgarian ethnogenetic conception” under the section “Ethnogenesis”, and a different section titled “Genetic origins”).

The first genetics section in our corpus appeared on November 21, 2003 in the page “Basques” (revision 1791706). It was titled “Genetics” and contained one paragraph, which discussed the frequencies of blood types among Basques and an unreferenced sentence reading, “[m]odern genetic techniques are also being applied to the Basques and it has been found that there is a great deal of difference between the Basques and their Spanish neighbours.” The number of pages with genetics sections increased steadily over time (figure S3). By our analysis cut-off date, 419 pages (6.4%) in the corpus had at least one genetics section.

Genetics sections had a mean length of 2,704 characters (11.3% of a page’s plain text characters) and a median of 1,464 characters (7.6%), though this distribution is skewed (figure S4). We compared genetics sections to two other common section topics for pages in the corpus: history and language. History sections tended to be much larger, with a mean length of 5,100 characters (38.5%) and median of 2,358 (36.0%), while language sections were comparable to genetics sections, having a mean length of 1,260 (10.5%) and median of 681 (7.0%). The mean and median proportions for each section type have remained stable over time (figure S5).

As with genetics keywords, genetics sections were far more common among the most-viewed pages in the corpus. Among the top 1,000 most-viewed pages, 304 (30.4%) had a genetics section. In a logistic regression of a binary indicator versus the log of pageviews, pageviews were a significant predictor of the presence of a genetics section (*P* > |*z*| ≈ 0, OR = 2.69, pseudo *R*-squared = 0.295). Wikipedia pages sometimes contain figures derived from genetic data, such as principal components analysis (PCA) plots, genome-wide ancestry plots, maps of historical migrations, or haplogroup distributions. We analyzed markup within the corpus (see Methods—Text analysis) and identified 266 genetics figures across 91 (1.4%) of pages. Again, the figures were more common among the most-viewed Wikipedia pages, with genetics figures being present in 81 (8.1%) of the top 1,000 most-viewed pages. A logistic regression of a binary indicator showed pageviews were a significant predictor of the presence of a genetics figure (*P* > |*z*| ≈ 0, OR = 3.14, pseudo *R*-squared = 0.311).

### Genetics sections are common in pages about nationalities

A majority of corpus pages about nationalities had genetics keywords and genetics sections. As a proxy for nationality, we use national-level demonyms (e.g. “Canadians”, “Spaniards”, etc.). We used a list of states based on United Nations definitions (see Methods—Data collection) as well as the four constituent countries of the United Kingdom, for a total of 202 potential pages.

Of these, 137 (67.8%) had a Wikipedia page about their demonym. These pages are among the most-visited pages in the corpus (165,005 mean pageviews for demonyms in 2025 versus 36,333 for others; 131,564 median pageviews for demonyms versus 5,702 for others). There were 93 (67.8%) pages that had genetics keywords (see figure 2A for a world map). The most common keywords across these pages were “genetic” (appearing in 65.0% of demonym pages), “DNA” (53.3%), “genom” (43.7%), “chromosom” (42.3%), and “gene(s)” (41.6%) (figure S6).

**Figure 2:**
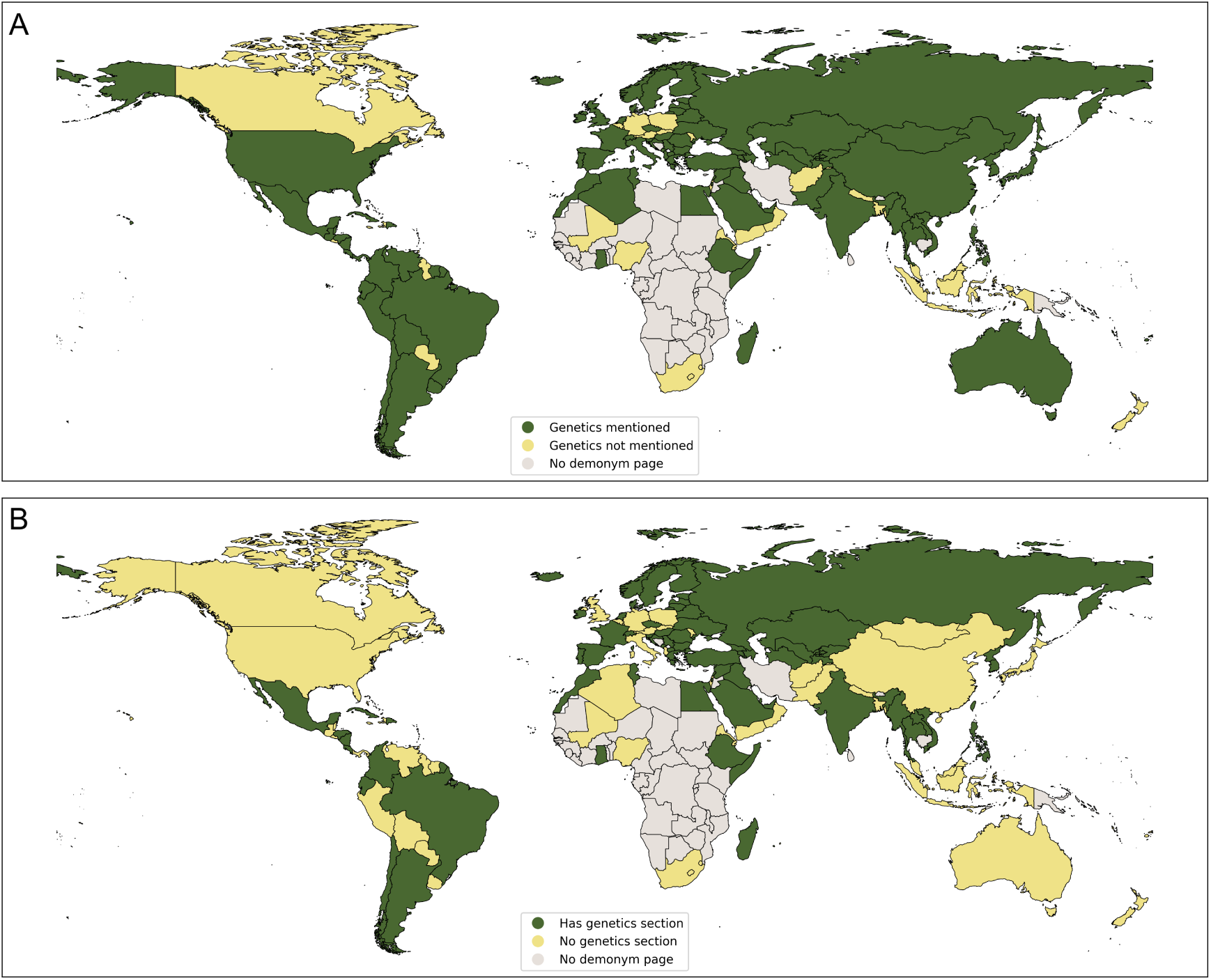
The majority of countries with Wikipedia pages about their demonym discuss genetics. Countries in green have Wikipedia pages dedicated to their demonyms, and genetics are mentioned. Countries in yellow have Wikipedia pages dedicated to their demonyms, and genetics are not mentioned. Countries in gray do not have Wikipedia pages about their demonyms. In contrast to the rest of the world, many African countries do not have pages for their demonyms. Of 137 demonyms that have Wikipedia pages, **(A)** 67.8% had genetics keywords and **(B)** 52.5% had a genetics section. While not all countries have pages for their demonyms, they all have “Demographics of…” pages (e.g. “Demographics of Namibia”). Of 197 demographics pages, 55 (27.9%) had genetics keywords and 13 (6.6%) had genetics sections. For maps of demographics pages maps, see figure S7.

Of the 137 pages, 72 (52.5%) had genetics sections (see figure 2B for a world map). Demonym pages tended to be longer than non-demonym pages in the corpus, with a median length of 31,280 characters (95% CI: 28488, 34080) compared to 21,725 (95% CI: 20106, 23344). The median number of characters in demonym genetics sections was 2,853 (95% CI: 2315, 3387), and the median proportion of text taken up by the sections was 9.6% (95% CI: 8.5%, 10.7%). In contrast, genetics sections in non-demonym pages had a median length of 1314 characters (95% CI: 1145, 1483) and a proportion of 7.0% (95% CI: 6.3%, 7.7%).

Using section headers, we further analyzed the contexts in which genetics keywords appeared, out of five possibilities: Biology; Origins; Culture; Race, ethnicity, or nationality; and Reference. Keywords that appeared before any sections were defined were considered part of the Introduction, and keywords could also occur in multiple contexts (see Methods—Text analysis and table S1 for definitions). The most common context as of the analysis cut-off date was Biology (72 pages, 52.6%) followed by Origins (37, 27.0%) and Culture (33, 24.1%) (figure 3 and table S2). The most common context was initially “Origins”, but beginning in 2010, sections titled “Genetics” (considered a “Biology” context here) became more common in pages about nationalities, and by the cut-off date, human genetics research in these pages most frequently appeared in such sections. Twenty-five pages (18.3%) contained figures that either cited genetics research (e.g. a pie chart of genetic ancestry) or were figures derived from genetic data, such as principal components analysis scatterplots (table S3).

**Figure 3:**
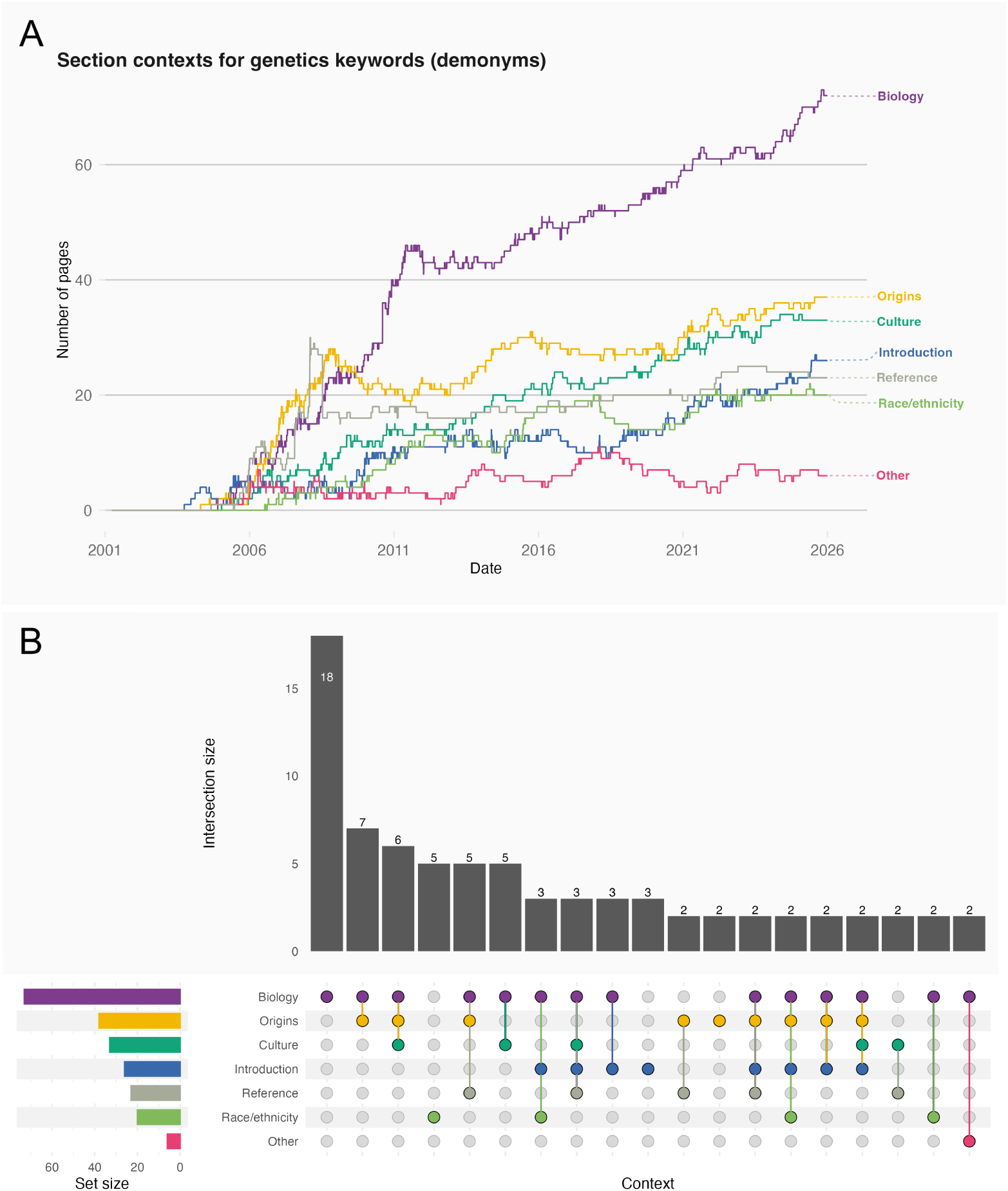
A growing number of Wikipedia pages about nationalities are incorporating genetics keywords, most often presenting research in a biological context. Genetics terms appear in a variety of contexts, such as ancient DNA research used to support archaeological findings (e.g. the page “Italians” cited genetic research about the origins of ancient Etruscans (*51*)) or modern genomic studies tracing migrations for historical research (e.g. the page “Portuguese people” cited a genomic study identifying migrations between 860 and 1120 CE (*52*)). **(A)** Beginning in 2010, biology became the dominant context for genetics terms in demonym pages, most often as a standalone section titled “Genetics”, separate from contexts such as history. By December 31, 2025, genetics keywords had appeared in a biological context in 72 pages (52.6%), while appearing in Origins (the second-most frequent context) in only 37 (27.0%). **(B)** Though genetics keywords often appeared in multiple contexts, in 18 cases they appeared exclusively in a Biology context; the second most frequently exclusive context was Race, Ethnicity, or Nationality (5 pages).

While some states did not have a page for their demonym, all of them had a “Demographics of” page (e.g. “Demographics of Kenya”). We consider these pages alongside pages about nationalities because they may contain text about the ethnic makeup of a country and, in some cases, users are automatically redirect from a demonym to a demographics page (e.g. “Kenyans” forwards users to “Demographics of Kenya”). Separately from the corpus, we analyzed 197 demographics pages and their 185,616 revisions; of these 197 pages, we found that 55 (27.9%) contained genetics keywords and 13 (6.6%) had genetics sections as of the cut-off date (figure S7). Four (2.0%) of the demographics pages had genetics figures (Bahrain, Italy, Mexico, Tunisia).

### Genetics sections are broad synthesis of published research

To study whether there are any dominant or influential publications in the corpus, we identified all citations and references in genetics sections and extracted their data and metadata (full details in Methods—Citation analysis). Across the 419 corpus pages with genetics sections, we identified 4,776 total citations within their genetics sections (here citations are unique within pages, but not across them). Analyzing citation markup metadata, we identified that the three most common citation types were academic journals (3,461 citations; 72.5%), websites (648; 13.6%), and books (275; 5.8%). For the 4,378 citations with years available, 3,976 (90.8%) were published after 2003, the final year of the Human Genome Project (figure S8).

Among academic journal publications, there were no dominant citations. Among 1,195 citations with unique digital object identifiers (DOI), the median number of citations was 1, with 835 (69.9%) cited in exactly one page (table S4 and figure S9). Twenty-three DOIs appeared in more than five pages, and the most frequently cited publication appeared on 18 pages: “Genetic Heritage of the Balto-Slavic Speaking Populations: A Synthesis of Autosomal, Mitochondrial and Y-Chromosomal Data”, a 2015 publication by Kushniarevich and Utevska et al. (*20*). Four publications were cited on ten or more pages (table S5). The three most-cited journals were the *American Journal of Human Genetics* (76 citations), the *European Journal of Human Genetics* (62 citations), and the *American Journal of Physical Anthropology* (56 citations) (table S6).

### Pages with genetics content have wide reach

We estimated corpus pages were viewed 239,019,588 times in 2025 (see Methods—Pageview data). Pageviews were skewed towards a small proportion of corpus pages; the top 1000 pages received 192,975,827 views (80.7%). While there were 970 corpus pages (14.8%) that contained genetics keywords, they received 158,285,287 views (66.2%). Similarly, 419 corpus pages contained genetics sections (6.4%), but received 90,743,136 views (38.0%). Pages with genetics figures such as PCA plots, genome-wide ancestry plots, haplogroup maps, or figures that cited genetics literature received 27,651,241 views (11.6%).

The 137 demonym pages received 22,605,730 views (9.5%) despite being only 2.1% of the corpus. Demonym pages with genetics keywords received 18,305,830 views (7.7%), and demonym pages with genetics sections received 14,683,729 (6.1%); these accounted for, respectively, 80.1% and 65.0% of views for demonym pages specifically.

### Users frequently discuss genetics in pages about ethnicities

Each Wikipedia page has an associated “talk page”, in which users can discuss the contents of a page. In cases where pages have editing privileges restricted (e.g. because of disputes over text or malicious editing), the talk pages are still open to contributors to provide feedback. Talk pages use the same markup language as regular pages, though sections are used to delineate discussion topics rather than to organize pages, e.g. a section may be titled “History section missing 1850s”. We collected all active and archived discussion topics from corpus talk pages and measured the presence of genetics keywords (see Methods—Talk page text).

Within the corpus, many talk pages are empty: 2,672 (40.8%) have no discussions at all; among the remaining 3,869 pages, there were 56,098 discussions Overall, 633 (9.7%) pages had at least one discussion which contained a genetics keyword, and 6,654 (10.1%) discussions had genetics keywords. Genetics keywords are much more likely to appear in the talk pages of Wikipedia pages that are more popular. A logistic regression on the presence of keywords in a talk page versus the log of daily page views was statistically significant (*P* > |*z*| ≈ 0, OR = 4.06, pseudo *R*-squared = 0.457).

The five pages with the highest number of discussions containing genetics terms are: “Black people” (272 discussions; 29.6% of all discussions for this page), “White people” (265; 30.1%), “Palestinians” (172; 28.6%), “Ashkenazi Jews” (138; 33.6%), and “English people” (130, 30.4%) (table S7). The talk pages for “Black people”, “White people”, and “Ashkenazi Jews” contained a high number of discussions about hereditarianism, which is the view that genetic inheritance plays a central role in traits such as behaviour and intelligence; we detail these in subsequent sections.

### Identifying topics in Wikipedia text on human genetics research

Wikipedia pages that incorporate human genetics research are written through varying lenses. They may focus on, e.g., ancestry and origins, or race and admixture, or comparing ethnic groups through genetic clustering. To identify and measure these topics, we extracted every paragraph of text that had a genetics keyword in its underlying markup across all pages in the corpus. This excludes infoboxes, tables, figures, titles, and navigation templates (e.g. sub-heading text such as “*See also: Genetic history of the Middle East*”). We then converted the markup to plain text and analyzed it using a set of terms related to a pre-determined set of topics; a single paragraph may have multiple topics. The topics and terms for this analysis were selected through a combination of manual curation and via analysis by a large language model (see Methods—Text topic analysis for full details and examples of text). We carried out this process both for Wikipedia pages in the corpus, and separately for their talk pages.

### In Wikipedia pages

We analyzed snapshots of corpus pages on December 31 of every year from 2001 to 2025. Of 970,644 paragraphs, we identified 34,532 that had genetics keywords in their underlying markup (see table S8 for annual counts and figure S10 for a time series). Of these, “ancestry & origins” was the most common throughout the corpus history, with 71.3% of corpus pages with genetics keywords having this topic present as of December 31, 2025. Example terms for this topic include “ancestry”, “origin”, “iron age”, “migration”, and “source population”. The five next-most common topics were:

1. “population genetics methods” (56.6%) (example terms: “allele”, “haplogroup”, “*F*_*st*_”)
2. “identity & ethnicity” (55.7%) (e.g. “nation”, “indigenous”, “ethnolinguistic”)
3. “language & linguistics” (52.2%) (e.g. “bantu”, “proto-language”, “dialect”)
4. “admixture & mixing” (46.0%) (e.g. “interbreeding”, “endogamy”, “absorbed”)
5. “group comparisons” (45.8%) (e.g. “difference”, “homogeneous”, “unique cluster”)

Some terms appear in multiple topics (e.g. the term “cluster” appears in the topic “statistics & clustering”, as well as in “unique cluster” as part of the topic “group comparison”). A full list of terms and topics is provided as a supplementary file.

Comparing topic frequencies from December 31, 2025 to December 31, 2005, the largest absolute increases were in “population genetics methods” (from 42.5% of corpus pages with keywords to 56.5%), “language & linguistics” (40.6% to 52.5%), and “subsistence mode” (5.7% to 10.0%), while the largest decreases came in “medical genetics” (14.2% to 9.9%), “religion” (25.5% to 17.6%), and “race” (29.3% to 17.3%). Hereditarianism was largely absent in Wikipedia text, however, we identified outliers in talk page data (detailed in the next section).

Using a subset of corpus pages that had at least five paragraphs of text with genetics keywords (187 pages total), we carried out hierarchical clustering on the topics within paragraphs, normalizing for the number of paragraphs with genetics keywords (see Methods—Text topic analysis). Figure 4 depicts the clustering structure of the topics observed within paragraphs. In the clustering tree (figure S11), the lowest mergers occur between the topics *{*“race”, “identity & ethnicity”, “ancestry & origins”, “admixture”*}*, which we call an “identity framing”.

**Figure 4:**
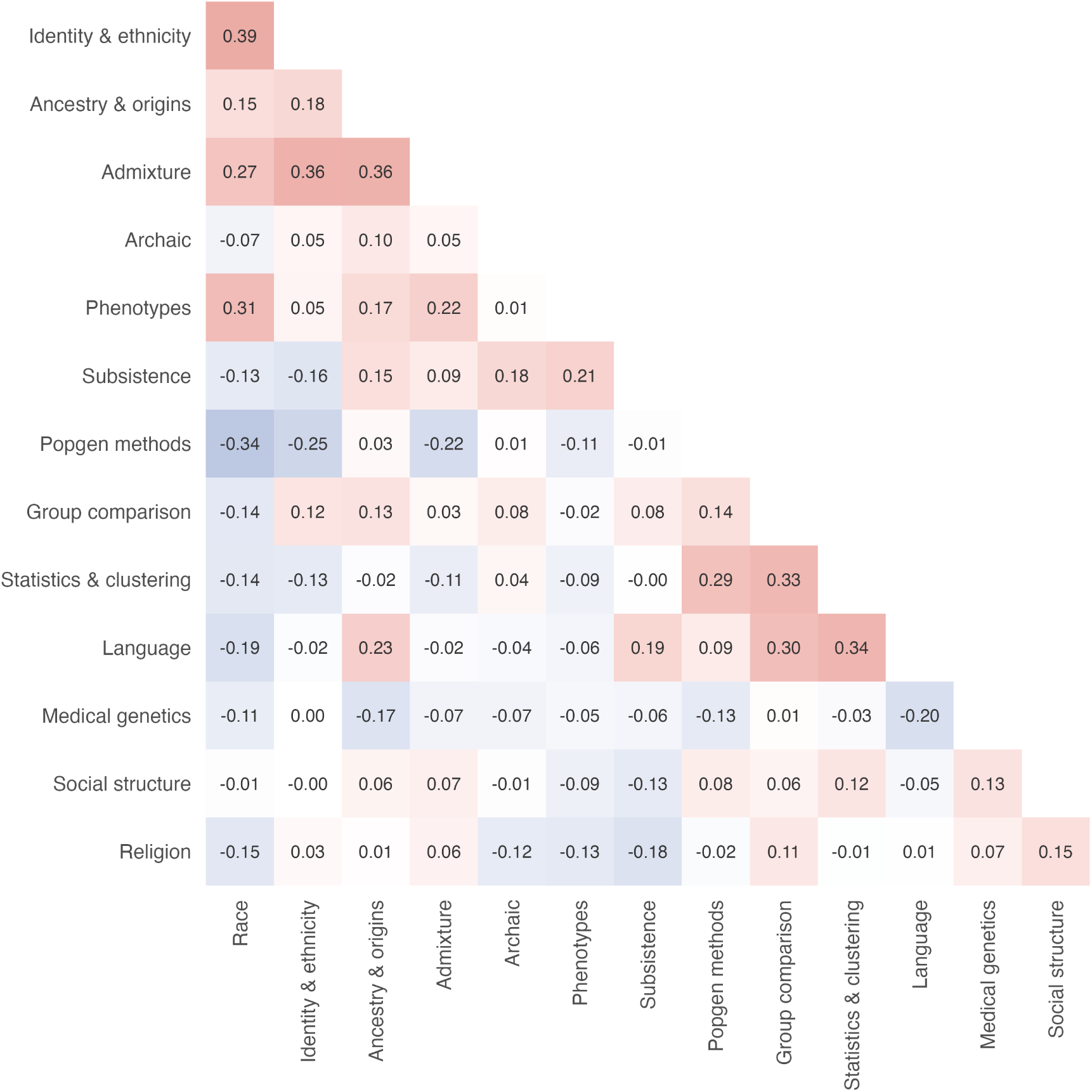
Hierarchical clustering of topics in corpus text. We identified every paragraph of plain text in the corpus that contained a genetics keyword. We then scanned for a set of terms related to pre-defined topics (the x and y axes). We reduced our data to pages with at least 5 paragraphs of text (187 pages) and calculated the proportion of paragraphs that contained each topic, calculated the correlation matrix, and performed hierarchical clustering. The clustering structure (figure S11) is negatively correlated and suggests pages are more likely to be written from either an “identity framing” (writing about e.g. race, identity, ancestry, origins) or a “methods framing” (writing about e.g. clustering, allele frequencies, linguistics, and group comparisons). However, these correlations are low-to-moderate.

Examples of pages with an identity framing include “Mulatto”, “Puerto Ricans”, “Mestizo”, “Hispanic and Latino Americans”, and “White Mexicans”. An example of text from “Puerto Ricans” (revision 1329310619):

> Studies have shown that the racial ancestry mixture of the average Puerto Rican (regardless of racial self-identity) is about 64% European, 21% African, and 15% Native Taino, with European ancestry strongest on the west side of the island and West African ancestry strongest on the east side, and the levels of Taino ancestry (which, according to some research, ranges from about 5%-35%) generally highest in the southwest of the island.

The next lowest mergers in the clustering tree are in *{*“population genetics methods”, “group comparison”, “statistics & clustering”, “language & linguistics”*}*, which we call a “methods framing”. Examples of pages with methods framing include “Bosniaks”, “Hmong people”, “Serbs”, and “Beta Israel”. An example of text from “Bosniaks” (revision 1330184327):

An autosomal analysis study of 90 samples showed that Western Balkan populations had a genetic uniformity, intermediate between South Europe and Eastern Europe, in line with their geographic location. According to the same study, Bosnians (together with Croatians) are by autosomal DNA closest to East European populations and overlap mostly with Hungarians. In the 2015 analysis, Bosnians formed a western South Slavic cluster with the Croatians and Slovenians in comparison to an eastern cluster formed by Macedonians and Bulgarians with Serbians in the middle.

These two framings are negatively correlated, suggesting pages tend to be written either from a more technical perspective, making heavy use of haplogroups, clustering, PCA, and language families, or more from a perspective of identity, race, ancestry, and admixture. However, this sample is small relative to the corpus and the correlations are modest, with a mean off-diagonal value of 0.03 and a range of (−.034, 0.39). Our approach is also sensitive to the terms used to define topics.

### In Wikipedia talk pages

For each page in the corpus, we next analyzed talk page discussions that contained genetics keywords in their markup. Of 730,560 paragraphs across all discussions, we identified 137,603 paragraphs that were part of discussions containing genetics keywords. We used the same topics as in corpus pages, though some of the terms related to topics differ (e.g. the verb “identify” in corpus pages is common in defining groups, while in talk page discussions it is simply a common verb). We do not analyze changes in talk pages over time as there is no equivalent of an annual snapshot; instead, we present the summary of all discussions over 25 years. figure S12 shows how many discussions containing a genetics keyword also contained a topic term across the history of the corpus.

The most common topic was “ancestry & ethnicity”, which reflects discussions about, e.g., origins, indigeneity, migration, peopling, and descent, and appeared in 76.0% of discussions with genetics keywords. (see tables S9 and S10). The five pages with the most discussions containing “ancestry & ethnicity” terms and genetics keywords were “Black people” (213), “White people” (209), “Palestinians” (157), “English people”, and “Ashkenazi Jews” (114) (see table S7 for the top 10 pages).

Talk pages can reflect content that users have wanted to include, but did not meet the thresholds for inclusion because of, e.g., low quality sourcing (see Supplementary Text—Wikipedia editing policies and guidelines). One example is hereditarian writing on intelligence and genetics, which is largely concentrated in the talk pages of “Black people” (52 discussions), “White people” (20 discussions), and “Ashkenazi Jews” (13 discussions). We analyze these pages as case studies.

### Case studies: Hereditarianism, race, and genetics in the corpus

While our corpus pages did not contain hereditarian writing as of the cut-off date, we identified historical instances in the pages “Ashkenazi Jews” and “Black people”. The page “White people” did not contain hereditarian content, but we did identify that the page “Genetic history of Europe” was initially written as a “Genetic History” section of “White people”.

“Ashkenazi Jews”

The page “Ashkenazi Jews” was the first of these three pages in the corpus to have a genetics keyword added. On March 9, 2004 (revision 2696783) a user linked to the Chicago Center for Jewish Genetic Disorders. In the next revision, text was added referring readers to the Wikipedia page “Race and intelligence”. In June 2005, there was a “Genetic Traits” section with a subsection titled “IQ”. “Genetic Traits” was renamed “Population genetics” on July 28, 2005 (revision 19761650). Text about IQ was moved to different areas of the page, often citing the 2006 paper “Natural history of Ashkenazi intelligence” by Cochran, Hardy, & Harpending (hereafter CHH) (*21*). The last explicit reference to CHH was in August 28, 2023 (revision 1172681072) before it was removed as a fringe viewpoint (see Supplementary Text—Wikipedia editing policies and guidelines). However, as of our cut-off date, there was an implicit reference to the work, where a sentence about Nobel Prize winners cited a 2006 article by Steven Pinker, which in turn relied on CHH (*22*).

As of the cut-off date for our analyses, the page contained 71,121 characters of plain text. The two largest sections were “History” (26,102 characters; 36.7% of page text) and “Genetics”, which contained seven subsections of its own: four about genetic origins, one about East Asian ancestry, one about medical genetics, and one about the Khazar origin hypothesis (table S11). There are also two standalone Wikipedia pages linked within the genetics section: “Genetic studies of Jews” and “Medical genetics of Jews”. None of these mentioned CHH or intelligence. While the genetics section is the last one in the page, our clickstream analysis shows it has a higher-than-expected number of link clicks for its position, suggesting readers are interested in this topic specifically (see figure S13, table S12 and Methods—Case studies)

“Black people”

The first revision in “Black people” to contain a genetics keyword was from July 10, 2005 (revision 18548755). The keyword appeared in the second sentence of the page (as part of the markup linking to the page “dominant gene”):

> “Since black featues[*sic*] are largely [[Dominant gene|dominant]], just one black ancestor is often enough for someone to be regarded as black, especially in traditionally non-black cultures, such as the [[Western World|West]].”

This text was removed the following day.

“IQ” first appeared on October 8, 2005 (revision 25145533) in a critique of IQ scores; the text was expanded to suggest that racial differences in IQ had a biological basis on October 26 (revision 26507345), and was removed on November 11 (revision 28007596). Hereditarian writer J. Phillipe Rushton was first cited on October 4, 2006 (revision 79539121) with a link to his website, which detailed his beliefs on race, genetics, and intelligence and contained excerpts to his book *Race, Evolution, and Behavior* (*23*); it was provided as a citation in the section titled “Definitions”, which was later renamed “Biological definitions”. Rushton was cited as late as May 22, 2007 (revision 132573700), regarding racial relations with Arabs. Another hereditarian writer, Arthur Jensen, was first cited on October 6, 2006 (revision 79896369), also on the definition of “Black people”, and was last cited March 30, 2007 (revision 119051297).

A section titled “Genetic Averaging” was introduced May 8, 2006 (revision 52225223) and removed May 15, 2006. A subsection titled “Admixture” was added to a section titled “Who is Black today?” on September 13, 2006, describing how Black individuals are defined. The name of this section changed over time (e.g. to “Intermediates” and “People of Black heritage”) though it continued to cite genetics research (e.g. revision 80267867 on October 8, 2006 cited an mtDNA study from Brazil (*24*) and a clustering study that included Ethiopian individuals (*25*)). The final genetics section to appear was titled “Genetic study” and was removed on June 3, 2007 (revision 135457540). As of the cut-off date, the page was organized into sections based on continental geography, and contained 35 genetics keywords; no genetics-related links appeared in the clickstream data in December 2025.

### “White people”

While we did not identify any hereditarian text in the page’s history, we did identify genetics keywords and sections. The earliest revision for the page “White people” is from January 7, 2003; the first genetics keyword appeared on November 30, 2003 (revision 1853629), and the first genetics section appeared September 30, 2005 (revision 24413995) and was titled “Black-white admixture in the Americas”, citing a United Press International piece about genetic research done by researchers at Penn State University (*26*). On April 16, 2006 (revision 48643798), a section titled “Genetic History” was created, and on August 20, 2006, it was spun off into the standalone page “Genetic history of Europe”. Various genetics sections were added and removed over the years, e.g. sections titled “Genetic Traits”, “Race and Genetics”, and “Physiology and genetics” and covered topics such as haplogroups, the genetics of skin colour, and human genetic variation. The last genetics section we identified was removed on May 13, 2013 (revision 554879919).

As of the analysis cut-off date, there were 51 genetics keywords. The only genetics-related link with enough clicks to appear in clickstream data was the page “race and genetics”, which was linked in the final paragraph of the introduction (link text bolded and underlined):

> “Contemporary anthropologists and other scientists, while recognizing the reality of **<u>biological variation between different human populations</u>**, regard the concept of a unified and distinguishable White race as a social construct with no scientific basis.”

Though it was the 19th link in the introduction it was the sixth most-clicked linked in the introduction (figure S14), and the 11th-most clicked link on the page.

### AI and human genetics research in the information ecosystem

We investigated the role of AI and online encyclopedias in the information ecosystem. Wikipedia is known to be part of the training corpus of chatbots, though the exact extent is not clear (*19*). We investigated whether chatbots also mention genetics when queried about nationality, as well as whether they refer to Wikipedia in their responses.

Further, we compared Wikipedia pages about nationalities, as well as our case studies, to Grokipedia. In contrast to the decentralized contribution structure of Wikipedia, in which users write content independently, Grokipedia is centralized and written by Grok, a large language model.

### Wikipedia frequently used as a source on nationalities by large language models

We investigated whether popular LLMs use Wikipedia as a source of information, whether the LLM responses include genetics keywords, and whether the LLM responses directed users to Wikipedia. We prompted three popular LLMs (ChatGPT, Gemini, and Claude) with “Tell me the history of [demonym]” for each of the 137 demonyms that had Wikipedia pages. We found that ChatGPT mentioned Wikipedia in 82.5% of responses, Gemini in 94.9%, and Claude in 100%; ChatGPT contained genetics keywords in 31.4% of responses, Gemini in 9.5%, and Claude in 18.2% (see Methods—Large language model queries).

In the second query, we prompted “Tell me about the genetic history of [demonym]. List your sources as URLs” for ChatGPT and Claude, and “Tell me about the genetic history of [demonym]” with a follow-up prompt “List every single URL you used during your search for this query at the bottom of your response” for Gemini (see Methods—Large language model queries). Wikipedia was linked in 71 (51.8%) responses by ChatGPT, 104 (75.9%) responses by Claude, and 114 (83.2%) responses by Gemini.

### Nationalities more likely to have genetics content on Grokipedia than Wikipedia

We identified 130 demonym pages that existed both in our corpus and on Grokipedia (see Methods— Grokipedia). In these pages, genetics keywords were more likely to occur on Grokipedia than Wikipedia, with 120 (92.3%) Grokipedia pages having genetics keywords compared to 85 (65.3%) on Wikipedia. Using binary indicators of genetics keywords on a page, this resulted in an odds ratio of 8.81 (95% CI: 1.64–89.22; *p* < 0.01). Genetics sections were also more likely to appear in a Grokipedia page, with 93 (71.5%) having genetics sections compared to 68 (52.3%) on Wikipedia, resulting in an odds ratio of 25.61 (95% CI: 7.16–141.10; *p* < 0.0001). Figure 5 visualizes the differences between demonym pages on the two projects in keyword counts and genetics section lengths.

**Figure 5:**
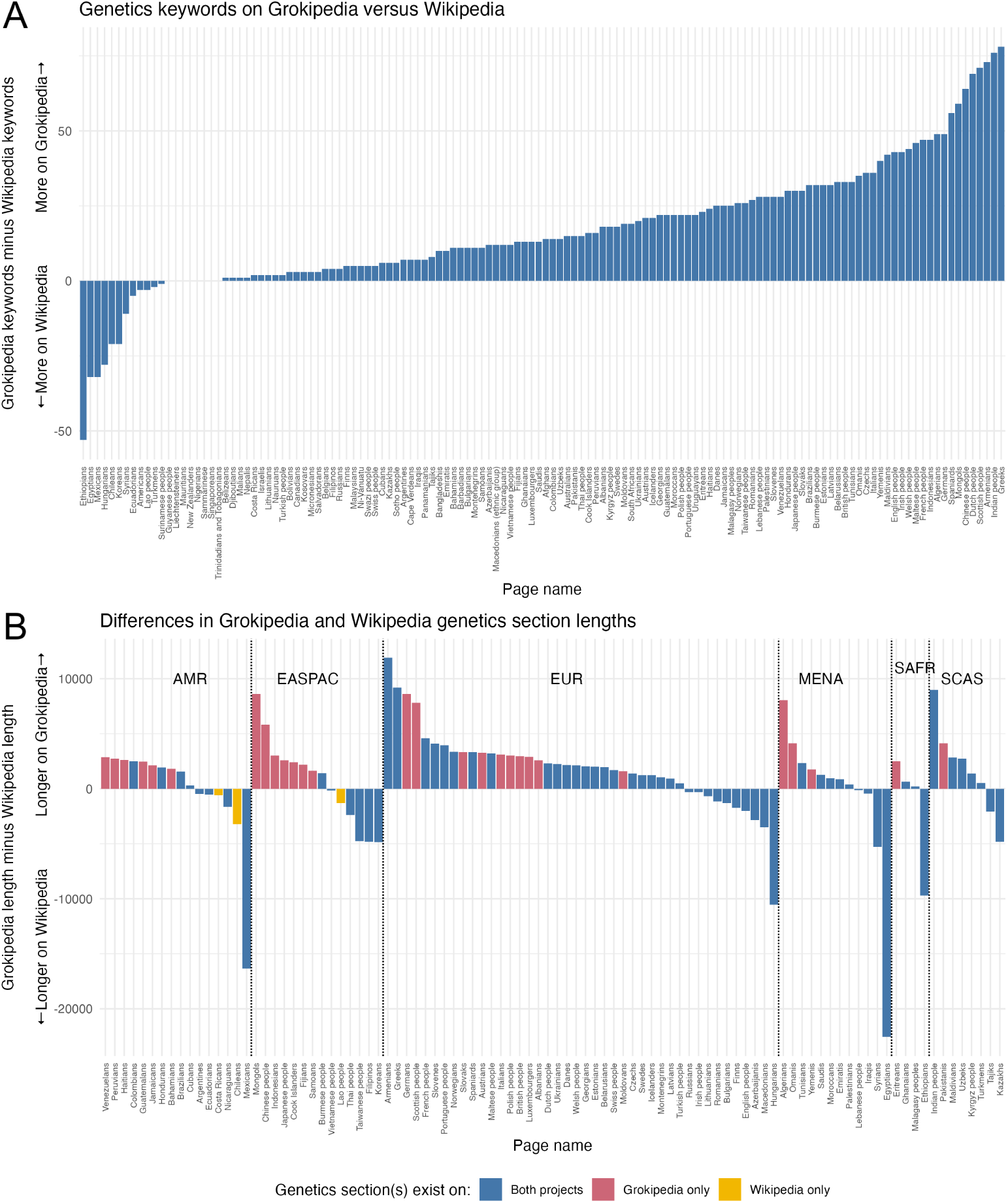
Grokipedia pages about nationalities are more likely to have genetics sections and have more genetics keywords. These figures compare demonym pages that exist on both websites. **(A)** The number of genetics keywords on a Grokipedia page minus the corresponding number on the Wikipedia page. **(B)** Differences in plain text length (number of characters) of Grokipedia sections and Wikipedia sections, arranged geographically (AMR: Americans and Caribbean; EASPAC: East Asia and Pacific; EUR: Europe; MENA: Middle East/North Africa; SAF: Southern Africa; SCAS: South/Central Asia). Grokipedia sections are more uniform in size compared to Wikipedia, but there are more demonym pages with genetics sections on Grokipedia than Wikipedia. Blue bars are pages that have genetics sections on both websites; red only on Grokipedia; marigold only on Wikipedia.

### Hereditarianism, race, and genetics on Grokipedia

We found that the Grokipedia pages “Ashkenazi Jews”, “Black people”, and “White people” all contained hereditarian text, and the pages “Black people” and “White people” defined the racial groups explicitly by genetics.

The Grokipedia pages “White people” and “Black people” contained more genetics keywords than their Wikipedia counterparts, at a respective 247 (compared to 51 on Wikipedia) and 201 (compared to 35 on Wikipedia). While neither Wikipedia page contained a genetics section as of the analysis cut-off date, the Grokipedia page “Black people” had a section titled “Genetic and Biological Foundations”, which contained 8.5% of the page’s text, and “White people” had a section titled “Genetic and Biological Characteristics”, which contained 7.8% of the page’s text. Both Grokipedia pages defined the racial groups with genetics in their first sentences, with “Black people” defined as “forming a distinct continental cluster in STRUCTURE analyses of global genomic data” (n.b. the supporting citation for this sentence is misrepresented; see Supplementary analysis—Grokipedia) and “White people” defined as “humans whose primary genetic ancestry derives from indigenous European populations”. The Grokipedia page “Ashkenazi Jews” had 183 genetics keywords, and 21.5% of its text was contained in a section (compared to 272 and 31.2% on Wikipedia); the genetics section was titled “Genetic Origins and Characteristics”.

“Ashkenazi Jews” contained multiple sections asserting that the group has higher-than-average IQ resulting from natural selection, including a section titled “Cognitive Abilities and Intelligence”, and cited CHH and hereditarian writer Richard Lynn. The page “Black people” contained multiple sections outlining negative social outcomes and asserted they were due to genetics (e.g. that high crime rates were caused by a variant in MAOA gene, which it calls the “warrior gene”) and cited Lynn, Jensen, and Rushton; it did not assert a genetic basis for any positive outcomes, such as for its “Achievements and Contributions” section. The page “White people” states that differences in polygenic scores of educational attainment and cognitive performance between Europeans and Africans are a result of “causal genetic contributions”, citing a blog, an unrelated BBC interview, and an unrelated research paper (see Supplementary text—Case study comparisons to Grokipedia for details).

We identified distinguishing words in the Grokipedia pages using log-odds ratios of word frequencies. Hereditarian words were in the 25 highest-ranking distinguishing words on the pages “Ashkenazi Jews” and “Black people”, and all three pages had genetics terms in their 25 highest-ranking words (see tables S13, S14, S15 for all terms, and Supplementary text—Case study comparisons to Grokipedia for details). On the Grokipedia page “Ashkenazi Jews”, the highest-ranking word was “IQ”, and the words “selection”, “cognitive”, “bottlenecks”, and “intelligence” all appeared in the top 25. In the page “Black people”, the word “IQ” was ranked ninth, and the top 25 contained the words “variants” and “analyses”, referring to (e.g.) “principal component analyses”.

## DISCUSSION

Using novel methods and datasets, we provide the first large-scale analysis of how human genetics research has been synthesized in the public-facing information ecosystem over the last 25 years. We find that in Wikipedia pages about ethnicity, nationality, and race, genetics terminology has become more common over time (figure 1), and that human genetics research is frequently placed into dedicated sections in these pages (figure S3). The proportion of pages incorporating human genetic research and genetics sections has increased as well, and the most popular pages—and pages about nationalities, in particular—are more likely to have genetics terminology and genetics sections (figure S6).

We show that one of the most common approaches is presenting human genetics research as a genetics section within a page on ethnicity, nationality or race; these sections have a mean size of 11% of a page’s total plain text, approximately the same size as sections dedicated to language, or about one-third of the length of a typical “History” section (figure S5). Rather than relying on a handful of sources or scientific publications, we found that 69.9% of citations in genetics sections appeared in only one page, with only four citations appearing in ten or more pages (see Methods—Citation analysis). The journals cited were also varied, with the most frequently cited being *The American Journal of Human Genetics* (table S5). Rather than relying heavily on a small set of sources or outlets, genetics sections on Wikipedia appear to reflect an earnest attempt to synthesize relevant sources of human genetics research.

Chatbots are growing as a source of information, making them an important new component of the information ecosystem. In our chatbot analysis, we prompted three popular models using “tell me the history of [demonym]” and found that, depending on the chatbot, Wikipedia was referenced in 82.5% to 100% of responses, and 9.5% to 31.4% of responses contained genetics keywords. When prompting the chatbots to “tell me about the genetic history of [demonym]”, Wikipedia was linked in 51.8% to 83.2% of responses.

Our analysis found that on pages about nationalities, genetics content is significantly more present on Grokipedia than Wikipedia. In the 130 pages that existed on both websites, genetics keywords appeared in 92.3% of Grokipedia pages (versus 65.3% on Wikipedia) and genetics sections in 71.5% of Grokipedia (versus 52.3%). In the case study pages “Black people” and “White people”, Grokipedia defines them explicitly in terms of genetics in the first sentence, and in the pages “Ashkenazi Jews”, “Black people”, and “White people”, it asserts that genes are causal factors underlying group-level measures such as IQ and crime rates.

### Limitations and future work

The pages in our corpus were those that were categorized as by Wikipedia users as ethnic groups and are not a comprehensive set (see Methods—Data collection); the corpus may contain false positives, e.g. extinct historical populations. We limited our study to the English-language Wikipedia, which has known systemic biases (*27, 28*). Our methodology is flexible and can easily be extended to include other categories of Wikipedia pages and other language editions of Wikipedia. Other language editions are an avenue for future research: the page “Spaniards” featured a genetics section in the Spanish- and French-language Wikipedia (*29, 30*) (Spanish-language revision 72575304, French-language revision 233898883), and the page “Russians” featured a genetics section in the Russian-language Wikipedia (*31*) (Russian-language revision 152703854).

We identified pages that were related to this research but deemed out of scope (see Supplementary Text—Related pages and categories). These included categories dedicated to genetics by ethnicity (21 pages), genetics by country (15 pages), Jewish genetics (9 pages), Y-DNA haplogroups (119 pages), mtDNA haplogroups (55 pages), 11 pages about origin hypotheses, and 29 pages on “genetic history” or “genetic studies” of regions or ethnicities.

### Social and ethical implications: Priming the public for genetic essentialism

When human genetics research is repackaged under discrete article topics such as those we observe in our analysis, there is a risk that readers can be primed to think of ethnicity, nationality, or race as genetically stable, uniform, and immutable—a genetic essentialist definition of human groups (*14*). Wikipedia’s discrete structure gives both each topic a separate page and each section a prominent place as a navigable link in a table of contents. We estimate that the 419 pages with genetics sections received over 90 million views in 2025, potentially reinforcing genetic essentialism to readers.

Genetics methods (e.g. clustering (*13, 32*)) and products (e.g. ancestry testing (*33, 34*)) are vulnerable to increasing genetic essentialist beliefs. Genetic researchers have sounded the alarm on the dangers of essentialism and typological thinking within the field (*4*), and it is understood that genetic essentialism is associated with many negative outcomes including racism and prejudice (*35*); it is further associated with opposition to policies aimed at racial equality (*14*) and with extremist ideologies (*36*). Human genetics research has been stripped of context and remixed in manifestos of perpetrators of racially-motivated massacres, part of a growing “curriculum of violence” (*37*).

Our work identifies a trend: genetics sections are becoming more common (especially among the most-viewed pages, such as nationalities) and taking up substantial portions of a page’s text, perhaps because there is contemporaneous research offering new insights into population genetics (as opposed to historical facts that may seem to change more slowly or be of less interest to Wikipedia contributors). Extrapolating from this, the Wikipedia page of every nationality or ethnic group could soon have a genetics section, likely with more negative consequences than positive ones (*5, 10, 38*).

The Wikipedia community has debated genetics sections at length, with at least ten discussions between 2009 and 2018 (*39*). A 2016 discussion on whether to remove them entirely cited concerns about genetic essentialism, politically motivated and decontextualized writing, outsized attention being given to genetics over (e.g.) language or history, and users’ lack of technical knowledge of population genetics. Ultimately there was no consensus to remove them, and the sections remained (see Supplementary Text—Wikipedia editing policies and guidelines). The Wikipedia community does, however, respond to scientific guidance. In a contrasting discussion from 2020 (*40*), the community debated whether hereditarian writings about race, genetics, and IQ were considered reliable sources. Citing scientific books and editorials, and statements from scientific societies, they concluded that the scientific consensus was that a genetic link between race and intelligence was a fringe theory, and hereditarian writing should not be used for citations.

While guidance from population geneticists for scientists/academics exists—e.g. on interpreting mixed-membership clustering (*41*) or the relationship between genetics and race and ethnicity (*42*)—there are few such resources that are intended for and widely communicated to those outside the field. This asymmetry is present in other fields, e.g. climate science (*43, 44*), and is detrimental to understanding research. Work by (*45*) has shown genetic essentialism is learned and that it, along with racism, can be reduced through education; given the massive reach of Wikipedia, there is room to be proactive and provide public guidance on how to synthesize and interpret human genetics research.

It is uncertain how Wikipedia’s text and structure influence chatbot responses, but Wikipedia’s free licensing and size make it a useful training corpus (*19*); our analysis found that Wikipedia was referenced in the majority of chatbot responses, suggesting it carries important weight. Looking forward, human genetics research could be remixed and repackaged using AI on an unprecedented scale, and with little transparency. Such remixing has been found in far-right communities and the manifestos of mass shooters (*5, 7*). A project like Grokipedia—which, according to xAI CEO Elon Musk, was created to “purge out the propaganda” on Wikipedia (*46*) and has been found to reflect his views (*47*)—represents a new version of this; many Grokipedia pages have been found to be similar to their Wikipedia counterparts, with divergent pages reflecting rightward ideological bias, particularly in pages about religion and history (*48, 49*). Our analyses demonstrate its explicit genetic essentialism, and while Wikipedia’s traffic dwarfs that of Grokipedia (2.1 billion versus 1.3 million monthly views in 2026 (*50*)), there is now a threat of automatically generating racist text under a mantle of scientific credibility.

In 2024, Panofsky et al. (*7*) argued that the way human genetics research is written, it has not been “weaponized”, but rather is akin to leaving a loaded weapon unattended. To fully realize the potential of human genetics research requires recognizing how it can be misused and misconstrued. Our study provides insights into how, over 25 years, human genetics research has been repackaged for public consumption in a good faith effort by the Wikipedia community. Our work underscores that the way human genetics research—and by extension any area of applied science—is communicated matters; that scientists’ words have weight throughout the information ecosystem; and, crucially, shows that those words echo and how far they can reach.

## Acknowledgments

We thank Jedidiah Carlson, Doc Edge, Ariella Gladstein, Simon Gravel, Arbel Harpak, Brandon Ogbunu, and Noah Rosenberg for their feedback.

## Funding

A.D.P. acknowledges support from the Natural Sciences and Engineering Research Council of Canada (NSERC) PDF-599527-2025. S.R. acknowledges support from the National Institutes of Health (NIH) Grant R35 GM139628 (S.R.). A.K. is supported by the National Science Foundation GRFP 2024366082 and as a trainee by NIH/NIGMS T32 GM149433, the Predoctoral Training Program in Biological Data Science at Brown University (A.K.).

## Author contributions

Writing—original draft: A.D.P. Writing—review and editing: A.D.P. and S.R. Conceptualization: A.D.P. and S.R. Investigation: A.D.P., A.K., and S.C.D. Methodology: A.D.P. Resources: S.R. Data curation: A.D.P., A.K., and S.C.D. Formal analysis: A.D.P. Software: A.D.P. and A.K. Visualization: A.D.P.

## Competing interests

We have no competing interests to declare.

## Data, code and materials availability

All scripts used to collect and analyze data are available at https://github.com/diazale/wikigenetics. All Wikipedia data are publicly available via the MediaWiki API (https://www.mediawiki.org/wiki/API:Action_API). Wikipedia data are subject to the Creative Commons Attribution-ShareAlike 4.0 International License (“CC BY-SA 4.0”). Grokipedia content is subject to the xAI Community License Agreement (https://huggingface.co/xai-org/grok-2/blob/main/LICENSE). Grokipedia content may also be subject to CC BY-SA 4.0. Grokipedia and LLM data used are available on Zenodo with DOI 10.5281/zenodo.21537405.

## Supplementary materials

Materials and Methods

Supplementary Text

Figs. S1 to S17

Tables S1 to S21

References (41–68)

## Supplementary Materials

## Materials and Methods

Wikipedia pages are written using a custom markup language that is rendered to generate the website. We analyzed the corpus by downloading the markup for every page and scanning it for strings related to genetics (e.g. “genetic”, “DNA”, etc.), which we call “genetics keywords” throughout this work. In addition to the written text that is visible to users, these keywords may appear as parts of citations (e.g. titles of publications or books, journal names, or in URLs), in markup that is invisible to users but part of the page (e.g. a link to a Wikipedia page about a haplogroup), or in file names (e.g. a file titled “admixture plot.jpeg”). Wikipedia pages can be subdivided into sections and sub-sections, which are delineated in the markup (e.g. “==History==”); for analyses of sections of pages, we use these delineations. To identify sections specifically about genetics, which we call “genetics sections” throughout this work, we searched for keywords within section titles. For an example of Wikipedia markup, see figure S15.

### Data collection

Wikipedia is an online encyclopedia that anyone can edit, with all published material made available under a Creative Commons Attribution-ShareAlike 4.0 International License (CC BY-SA 4.0). All Wikipedia data for this project were collected or derived from the English language Wikipedia, which at the time of writing contained approximately 7.1 million encyclopedia articles (which we call “pages”). The cut-off date for data was December 31, 2025 (inclusive). In this supplement we provide a list of all pages that were analyzed. However, as Wikipedia is dynamic, some pages may have been renamed, merged, or deleted after the cut-off date. Consequently a small portion of pages used in our analyses may no longer be publicly available as of the date of publication of this research. Some revisions were suppressed by Wikipedia administrators for violating website policies on copyright, malicious content, targeting individuals, or gross vulgarity. The revisions are visible in the revision histories, but their contents are not accessible. All page data were collected January 4–9, 2026, with the following exceptions: the data for pages for “Black people”, “Brown (racial classification)”, and “Native American identity in the United States” were downloaded on January 27, 2026; the data for all pages outlined in the supplementary section “Related pages and categories” were downloaded on January 27, 2026; and the data for the page “Demographics of Barbados” were collected February 3, 2026.

Wikipedia pages can be moved, which is equivalent to renaming them (e.g. the page “Spanish people” was created in 2004 and moved to “Spaniards” in 2015). In general, revision histories are fully preserved during these moves. Each page analyzed in this research is defined by its revision history, i.e. when discussing the page “Spaniards” we are discussing the full revision history of the page both under its current name (“Spaniards”) and its previous name (“Spanish people”).

All data were collected using the publicly-accessible MediaWiki API, with command line and Python scripts provided at https://github.com/diazale/wikigenetics. In addition to Wikipedia, the Wikimedia Foundation has several other projects. We used the MediaWiki Action API to access Wikipedia page information; the MediaWiki Analytics API to access pageview information; Wikidata, a sister project containing knowledge graphs, which we use to aid in data collection; and Wikimedia clickstream data, which we use to measure how frequently links in Wikipedia pages are clicked.

### Corpus of Wikipedia pages analyzed

We selected Wikipedia pages for analysis if they met at least one of two criteria: (1) the page focuses on a country-level demonym (e.g. pages entitled “Americans”, “Canadians”, “Russians”, etc.); or (2) the page belongs to a relevant Wikipedia category, which we outline below. Wikipedia pages are placed into categories by users—either manually or with semi-automated tools—by adding specific categorization code to a page (e.g. “[[Category:Ethnic groups in Canada]]”; see figure S16); these categories are intended to provide a list of pages related to a broader topic. Categories are visible on a web browser at the bottom of a page. A page may belong to multiple categories with no upper limit, or no categories at all, and categories are not mutually exclusive. Categories can also have sub-categories. For example, the category “Ethnic groups in Canada” is a first-level sub-category of “Ethnic groups by country” and itself contains the sub-category “Ethnic groups in Canada by city”. We refer to the level of sub-categorization as “depth”. Categories only list pages whose most recent revision contains the categorization code, i.e. category membership is a snapshot of whenever it is accessed—it is not tracked over time, and one cannot identify pages that were formerly in a category by examining the category alone.

For our analyses of demonyms, we use the demonyms listed in the Wikipedia pages for their respective countries. For our list of countries for demonyms, we used current states or observer states of the United Nations (except the Vatican City), as well as associated states, extant former member states (*53–55*), and the four constituent countries of the United Kingdom, for a total of 202 states. These are imperfect proxies as some demonyms may cover multiple countries (e.g. “Koreans”, “Irish people”). Categorization and sub-categorization are not standardized across Wikipedia, and in some cases we manually identified sub-categories that we included for this study. Examples include: Ethnic groups officially recognized by China (which contained 61 pages including “Han Chinese”); Multiracial ethnic groups in the United States (which contained 17 pages including the page “Hispanic and Latino Americans”); Multiracial ethnic groups in insular areas of the United States (which contained the page “Puerto Ricans”); and African nomads (which contained 35 pages including “Khoisan”). A full list of categories with the number of pages used in this study is available in table S16. Since a page may belong to multiple categories, the total number of pages analyzed will be smaller than the sum of pages in each category. We also downloaded demographics pages. These were pages that focused on country-level demographics (e.g. the page entitled “Demographics of Canada”). These were retrieved from the category “Demographics by country” and compared to the UN list.

### Wikipedia page text

All Wikipedia pages can be edited. Each edit of a page is a “revision” and has an associated integer ID that is unique across the English-language Wikipedia. Each revision is connected as a linked sequence, allowing for comparisons between any two versions of the same page. The contents of these revisions can be retrieved via a browser, or through the Wikimedia API by using the revision ID or the page name. In addition, each revision carries metadata such as the date and time of the edit and summaries of the edit. Any revision can be accessed using the URL format:
https://en.wikipedia.org/wiki/index.php?oldid=REVISION_ID
or
https://en.wikipedia.org/wiki/Special:PermanentLink/REVISION_ID

Wikipedia text is written in a specialized markup language sometimes called “wikitext”; we refer to it as “markup” throughout this work. All user-generated content on the website is rendered from markup; it contains all text, links, citations, tables, figures, etc. The markup of every revision of a Wikipedia page is available for retrieval, with rare exceptions (e.g. copyright violations, malicious contributions, or gross vulgarity). The process we used for this study was to retrieve all pages within each category and sub-category and remove any that were not relevant (e.g. pages about ancient peoples, census results, specific individuals, etc.). The total number of removed pages was 557. For every remaining page, we retrieved every revision ID in its history and its metadata, and for every revision ID, we retrieved the markup.

There were 6,541 corpus pages and 2,864,806 corpus revisions spanning from March 28, 2001 to December 31, 2025. Additionally, we analyzed a complementary set of 197 pages titled “Demographics of X”, where “X” is a country. These pages held 185,617 revisions from January 22, 2001 to December 31, 2025. In total, our main analysis dataset consisted of 6,738 pages and 3,050,422 revisions. Wikipedia pages are manually delimited into sections in the markup. We consider markup written before sections have been defined to be the “Introduction”. It is possible for a Wikipedia page to have no sections defined; in this case, the entire page is considered the “Introduction section”.

Section lengths are calculated using the character counts of plain text. A small number of pages feature template text, in which text written in one Wikipedia page is re-used in another by referencing a variable name; this underestimates the number of keywords and the lengths of sections because the markup only contains a variable name.

### Wikipedia talk pages

Every Wikipedia page has an associated talk page, where users can discuss the contents of the page, for example to suggest improvements, identify errors, etc. It can be reached with the prefix “Talk:” concatenated with the page’s name (e.g. “Talk:Canadians”). Talk pages are edited in the same manner as regular pages and their revision histories and contents can be accessed as with regular pages. Sections in talk pages are delineated with markup as in regular pages, with each section heading indicating a topic of discussion. Wikipedia volunteers also regularly clean talk pages by, for example, removing vulgarity, off-topic discussion, or non-sequiturs.

Unlike regular pages—where text is regularly revised, removed, modified, or split off into another page—comments left on talk pages are preserved; to prevent talk pages from growing too large, old comments are moved to archive pages. Archive pages are subpages of talk pages (accessed with a forward slash), and by convention are a sequentially numbered series of pages titled “Archive”, e.g. the first talk page archive for “Canadians” can be found at “Talk:Canadians/Archive 1”. We used the Wikimedia API to identify all subpages of talk pages for every Wikipedia page in our study, and collected the revision histories and contents for every subpage beginning with “Archive” for associated talk pages in our study.

When analyzing the contents of talk pages, we combined all archived text and all text on the talk page available as of the latest revision before our cut-off date. We identified instances of duplicated text (e.g. archived text may have been accidentally copied to multiple archives of the same talk page), keeping only the first appearance. We searched for genetics keywords within every section of the combined talk page contents, including section titles. Introduction sections in talk pages were excluded as these almost always consist of navigation aids and internal documentation rather than user-generated discussions.

### Clickstream data

Wikipedia clickstream data is available as far back as November 2017 at https://dumps.wikimedia.org/other/clickstream/. It provides monthly aggregated data for the number of visits to a page, as well as the source of the visit. The data are provided in the format *{*referrer, target, type, count*}*. For websites outside of Wikipedia, the referrer is listed as “other-empty” and the type is “external”; for links between Wikipedia pages, the referrer is the source page and the target is the destination page; the type is “link”. These data are aggregated by month, so if there are multiple revisions of the referring page in a month, it is not possible to know which revision was used. For this reason, we only analyzed one month in the case studies. If the same link appears multiple times in a Wikipedia page, it is not possible to know which one was clicked. In our analyses, we assign clicks to the first appearance of a link, meaning we underestimate the number of clicks for links that appear further down the page. Clickstream data is not available for links clicked fewer than 10 times in a month.

We used clickstream data to measure readers’ interest in the genetics of case study populations. Readers are most likely to click links in the opening section of a page, and text near the top of a page is more likely to be read than text at the bottom of a page (*56*). Following this logic, links that exist further down in a page but receive higher-than-expected clicks suggest that the section is being read more often than expected. Many Wikipedia pages have redirects, which account for typos, alternate spellings or names, etc. (e.g. “African-Americans” redirects to “African Americans”). Clickstream data for a page includes the visits from redirects. For our clickstream analysis for case studies, we used the following process:

1. Extract all outgoing links (“target” links) from the case study page, as well as their click counts, giving a (target, count) tuple
2. Use the Wikipedia API to identify all redirects for each target link, giving a (redirect, target) tuple
3. Identify all links in the case study page (the “wikilinks”), giving a (wikilink, redirect) tuple
4. Perform a join on these tuples, giving a (wikilink, redirect, target, count) tuple

### Pageview data

Pageviews are available as far back as July 2015 and are available for each page using the Wikipedia Analytics API at:

https://doc.wikimedia.org/generated-data-platform/aqs/analytics-api/.

We used the following parameters in our API call:

1. Project: en.wikipedia.org
2. Access: all-access
3. Agent: user
4. Granularity: daily
5. Start: 2025010100
6. End: 2025123100

These parameters retrieve daily views for all of 2025 for a given page and include desktop, mobile app, and mobile web users, and exclude automated page access (e.g. web crawlers). Most Wikipedia pages have redirects that allow for common typos, alternate names or spellings, etc. We identified each page’s redirects via the Wikipedia API, and calculated a page’s total views as the sum of the views to the page itself as well as all of the views of the redirects.

### Text analysis

To identify text related to genetics, we searched for substrings within the markup of the Wikipedia pages we extracted for our analyses. We term the substrings “genetics keywords”. The list of substrings is:

- admix
- autosom
- chromosom
- the regex pattern ‘\bgenes?\b’
- genetic
- genom
- genotyp
- haplo
- mitochon
- mt-DNA
- mtDNA
- (mt-DNA)|
- (mtDNA)|
- principal component
- PCA plot
- x chromosom
- x-chromosom
- y chromosom
- y-DNA
- y-chromosom
- yDNA
- (yDNA)|

We excluded specific strings that regularly triggered false positives: “genetic mod” and “genetically mod”, which referred to agriculture or scientific activity; and “dnaindia”, the URL for Daily News and Analysis (DNA), an Indian news program. Keywords are case insensitive, except for “DNA”. For figures that show counts of keywords, substrings contained within other strings (e.g. “DNA” in “Y-DNA”) were adjusted in downstream statistics to ensure they were not double-counted. The keywords (mtDNA)| and (Y-DNA)| indicate markup used to link to a page about haplogroups, e.g. the wiki markup [[Haplogroup N (mtDNA)|N]] generates the text “N” with an underlying link to the page entitled “Haplogroup N (mtDNA)”.

Markup was parsed using the *wikitextparser* (*57*), *mwparserfromhell* (*58*), and *wikiciteparser* (*59*) Python libraries. To calculate the length of plain text, we first parsed the text to remove citations, navigation templates, tables, and underlying links, leaving only readable prose. We then calculated the number of characters in the remaining text, stripping any trailing or leading spaces.

Inline references in Wikipedia pages are generated using markup code or tags. These are rendered as footnotes or endnotes in the page. When analyzing pages and talk pages, we consider inline references to be part of the section in which the citation appears.

We identified genetics sections as those that contained genetics keywords in the section title. To identify history sections, we searched for the regex pattern (?i:histor) in the section title. For language sections, we used the pattern(?i:language)|(?i:linguis).

For demonym and demographics pages we analyzed, we categorized the contexts of genetics keywords. When a keyword was identified, we derived a context based on the title of the section that contained the text with the keyword. We assigned contexts by searching for substrings in the section title, as listed in table S1. Keywords can appear in multiple contexts, e.g. a section may be titled “Genetic origins”, which would have the contexts “Biology” and “Origins”. For temporal analyses of keywords in pages and talk pages, we use daily counts of keywords. Since pages often have multiple revisions in a single day, we select the final revision of a day (according to Coordinated Universal Time; UTC) as the canonical version for tabulating that day’s counts.

To identify figures related to genetics, we searched for keywords using a regular expression pattern within markup identified as media (e.g. [[File:Filename|Caption]]). This would identify figures that are genetics-based (e.g. PCA plots, admixture plots) or figures that cite genetics (e.g. distribution maps, haplogroup maps).

### Text topic analysis

To analyze topics in text, we identified paragraphs of plain text that had genetics keywords in their underlying markup, such as in citations or links. For example, the following markup (from “Italians”, revision 1329753224) contains the keywords “genom” in the title of a paper within a citation:

~~~
Italians, like most Europeans, largely descend from three distinct lineages:<ref name=“Indo-European” /> Paleolithic [[hunter-gatherer]]s, such as the [[Epigravettian]] culture, who arrived in the Italian peninsula as early as 35,000 to 40,000 years ago;
{{Cite journal |last=Posth, C., Yu, H., Ghalichi, A. |date=2023 |title=Palaeogenomics of Upper Palaeolithic to Neolithic European hunter-gatherers | journal=[[Nature (journal)|Nature]] |volume=615 |issue=2 March 2023 |pages=117{126 |bibcode=2023Natur.615..117P
|doi=10.1038/s41586-023-05726-0 |pmc=9977688 |pmid=36859578}}
[…markup truncated]
~~~

The plain text is:

> Italians, like most Europeans, largely descend from three distinct lineages: Paleolithic hunter-gatherers, such as the Epigravettian culture, who arrived in the Italian peninsula as early as 35,000 to 40,000 years ago; Neolithic Early European Farmers who migrated from Western Asia and the Middle East during the Neolithic Revolution 9,000 years ago; and Yamnaya Steppe pastoralists who expanded into Europe from the Pontic–Caspian steppe of Ukraine and southern Russia during the Indo-European migrations 5,000 years ago.

Though the plain text does not contain any genetics keywords, the underlying markup does, and this paragraph is added to the topic analysis. Plain text excludes templates, which are frequently used for navigation (e.g. “infobox” templates that are common at the beginning of Wikipedia pages), as well as figures. Tables are also usually stripped out, however, manually formatted tables may sometimes bypass automated filters.

For Wikipedia page text, we used a snapshot of the latest revision of every year (i.e. the last revision whose UTC date-time fell before January 1 of the next year). For Wikipedia talk page text, we used all discussions, including active and archived discussions that met the analysis cut-off date. For talk page text, we included paragraphs within a discussion that contained a genetics keywords, even if the paragraph itself did not contain a keyword in its underlying markup. This is because discussions have a free-flowing structure frequently referencing previous paragraphs but without citations or links.

We determined which topics to analyze and which terms would relate to each topic through manual curation and the aid of an LLM (Claude Opus 4.8). We provided the LLM with the plain text of corpus snapshots as well as our topics of interest.

We used the following prompts to begin the analysis:

> I have a collection of documents about ethnic groups, which I have scanned for keywords related to genetics. If a paragraph contains a keyword, it is added to the corpus I am analyzing. I want to identify patterns in the types of technical terminology used (e.g. terms like cluster or haplogroup) as well as topics such as ancestry, origins, phenotypes, physical characteristics, health, or differences or similarities to other ethnic groups

> This is a sample of some paragraphs I would use. Are there terms here that are missing from the categories that could be relevant? *[sample attached]*

> Note that the motivation for this analysis is to understand how the text is written, not to derive conclusions about the ethnic groups.

> **Claude:** When you say ‘how the text is written,’ which is closest?

> **Respose:** Which technical terms/topics appear (vocabulary content)

We then iterated with the LLM with samples from different time periods in our corpus, removed terms that we flagged as false positives, and selected the 14 topics that were most relevant to our study, plus hereditarianism.

The terms for the topics on regular pages compared to talk pages were largely similar except for the following, either because they were infrequent in the data (e.g. from grammatical differences in page writing versus talk page writing) or triggered too many false positives (e.g. users on talk pages discussing each others’ IQ):

- Phenotypes, talk page only: “fair skin”, “light skin”, “skin tone”, “dark skin”, “blonde”, “olive skin”, “freckles”, “blond”
- Race, talk page only: “coloured”, “whites”, “pardo”, “pardos”, “colored”, “criollo”, “zambo”, “criollos”, “mestizos”, “castizo”, “blacks”, “casta”
- Identity & ethnicity, talk page only: “self-declared”, “declared”, “classified”, “enumerated”, “self-reported”, “identify”, “classification”
- Hereditarianism, prose only: “jensen”, “iq”
- Hereditarianism, talk only: “iq test”, “iq scores”, “iq score”, “iq gap”

We provide JSON files of our topics and their terms as supplementary files:

- categories_prose_custom_v2.json
- categories_talk_v3.json

The list of topics is:

- Population genetics methods
- Statistics and clustering
- Ancestry and origins
- Phenotypes
- Medical genetics
- Group comparison
- Subsistence mode
- Admixture
- Archaic and deep lineage
- Social structure
- Race
- Identity and ethnicity
- Language and linguistics
- Religion
- Hereditarianism

If a paragraph contained any term related to a topic, we defined it as containing that topic. We ran hierarchical clustering on the paragraphs that contained a topic. To adjust for very long pages with many paragraphs containing genetics terms, we used the proportion of paragraphs within a page that contained topics. This was because pages with many paragraphs of genetics-related content tended to cover every topic, skewing the signal. We used a subset of pages with *n* ≥ 5 paragraphs with any topics identified, resulting in a subset of 187 pages; running on the full set of pages did not change the overall conclusion (that writing tended to follow a “methods framing” or “identity framing”). We excluded hereditarianism from hierarchical clustering because it is largely not present in Wikipedia text.

We attempted topic modeling using BERTopic (*60*) using multiple sentence transformer models (all-MiniLM-L6-v2, all-mpnet-base-v2, allenai-specter) but the clusters returned were uninformative, reflecting either geographical distributions of populations or forming one large cluster around a small set of technical terms like “haplogroup”.

### Regression modeling

To model whether more popular pages (i.e. pages with more views) are more likely to have genetics keywords, we use a logistic regression model with a binomial link. The outcome variable is a boolean indicator if genetics keywords are present in the most recent revision of a page, and the explanatory variable is the log of (average daily pageviews + 1). See Tables S17 and S18 for results for the corpus and talk pages, respectively. We use this transformation because the pageviews for pages in our analysis follow an approximately log-normal distribution, and we add a constant to adjust for pages with zero views.

Our model for whether genetics keywords or genetics sections appeared was:

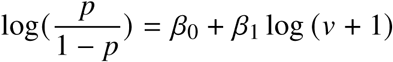

where *v* is the mean daily page views for 2025 and *p* is the probability that a page has a genetics keywords or contains a genetics section.

### Citation analysis

Wikipedia citations are primarily stored in reference (“ref”) tags, e.g. Diaz-Papkovich et al. Users may input any text in between the ref tags, e.g. a raw URL or a manually-written citation. Wikipedia provides a variety of citation templates that users may fill in to generate a properly formatted citation. We extract the details of these templates using the *wikitextparser* (*57*), *wikiciteparser* (*59*), and *mwparserfromhell* (*58*) Python libraries. We use the template name to identify the type of reference (e.g. “cite journal” is an academic journal, “cite news” is a news source, “cite web” is a website, etc.). We extracted identifiers from the templates such as URLs, DOIs, and titles and used these to count unique citations. A second, less-common type of reference uses shortened footnotes instead of ref tags. These are formatted as *{{*sfn|Author|Year*}}* in the text with full reference details elsewhere in the page. We identify these in the text using the same Python libraries.

### Case studies

For the case studies, we selected the three pages with the most talk page discussions that mentioned hereditarian keywords: “Ashkenazi Jews”, “Black people”, and “White people”. For these pages we analyzed historical revisions up until the cut-off date (December 31, 2025), talk page discussions up until the cut-off date (including discussions in the archive subpages), and clickstream data for December 2025. For the historical revisions, we used the same genetics keywords and keyword analysis as for the rest of the corpus.

For clickstream analysis, we used the most recent revision of a page in December 2025. Clickstream data is aggregated by month and links can be moved between revisions, meaning clickstream analysis will be approximate at best. Clickstream data does not indicate which link within a page was clicked. We assign all clicks to the first time a link appears in a page. This means that estimates of clicks for sections are biased upwards for sections that appear earlier in a page and biased downwards for sections that appear later in a page. We also removed navigation templates from this analysis (example in figure S17).

Previous research has shown that the relative position of a link influences how often it is clicked, with links that appear earlier in a page more likely to be clicked than those later on (*61*). Despite being the last section of prose in the page, the “Genetics” section received 2,150 (4.0%) of 53,352 clicks that month (figure S13)

### Supplementary data from Wikidata

Wikidata is a sister project of Wikipedia, available at https://www.wikidata.org. It is a knowledge graph that contains user-generated metadata about Wikimedia Foundation projects, including Wikipedia pages. To help filter pages from the category scraping, we queried Wikidata for a list of all Wikipedia pages that were about ethnic groups, excluding historical ethnic groups. The query was:

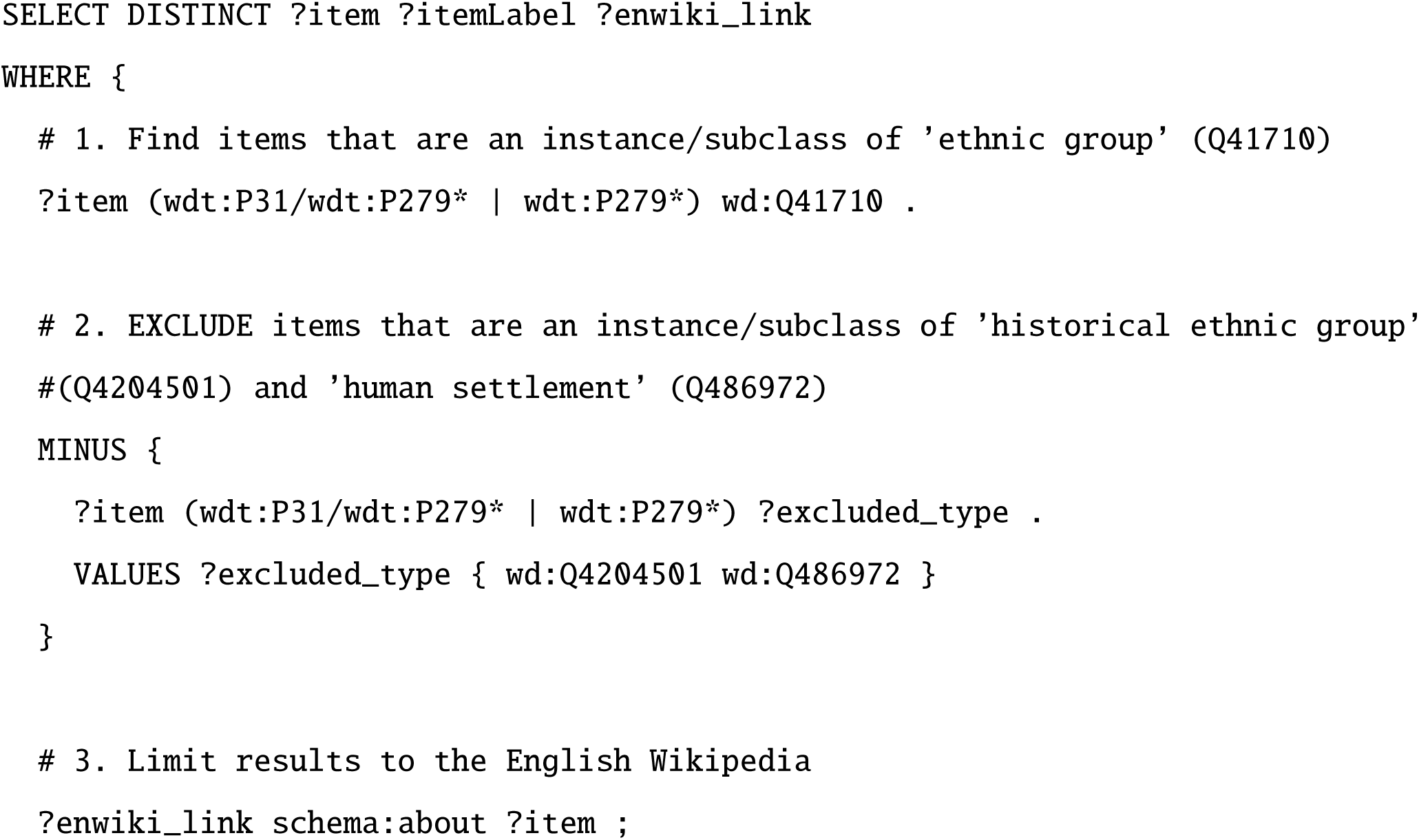
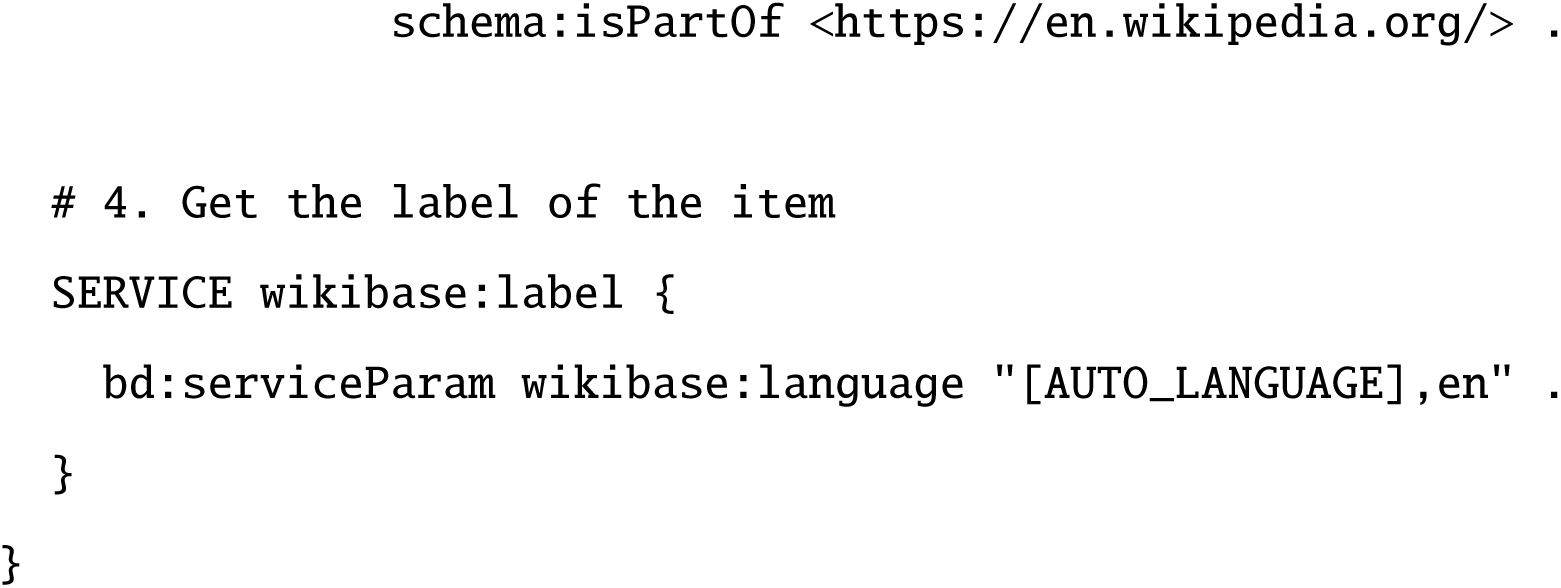

This query returned 10,979 pages from the English Wikipedia. When scraping certain categories, we compared page names to this list: if they were present, they were downloaded; otherwise, they were skipped. This was to exclude Wikipedia pages about geographical locations, individual people, historical events, etc. As Wikidata is user-generated, the query result is not a comprehensive list of Wikipedia pages about ethnic groups, but rather a collection of Wikipedia pages that users had marked on Wikidata as being an instance of an “ethnic group”. For these reasons, we chose to use Wikidata as a supplementary data source rather than a sample frame.

### Large language model queries

We queried three publicly available large language models (LLMs) — ChatGPT (GPT 5.3; OpenAI), Gemini (3 Fast; Google), and Claude (Sonnet 4.6; Anthropic) — using the free-tier versions of each platform available for access from April 3, 2026, to April 16, 2026. For each of the 137 nationalities with a Wikipedia page in our corpus, we submitted two sequential prompts within the same conversation thread to each LLM studied.

The first prompt requested the LLM describe the history of a demonym: “Tell me about the history of [demonym]. List your sources as URLs.” For Gemini, the prompt was modified to: “Tell me about the history of [demonym]. List every single URL you used during your search for this query at the bottom of your response.” The revision of this prompt was necessary to standardize Gemini’s source citation outputs. Specifically, Gemini has a source panel in the form of a collapsible sidebar that appears alongside responses where Gemini has performed a live Google Search. The Gemini 3 Fast model only performed a live web search and thus included sources in the output if it determined the query to be sufficiently complex or time-sensitive. We found our modified prompt produced more complete and consistent source reporting. We validated all three prompts in a trial study on a smaller sample of demonyms to confirm that response lengths were comparable across LLMs.

The second prompt asked the LLMs to describe the genetic history of the demonym with the same modified sourcing request across LLMs. Specifically, “Tell me about the genetic history of [demonym]. List your sources as URLs.” was used for ChatGPT and Claude, and “Tell me about the history of [demonym]. List every single URL you used during your search for this query at the bottom of your response.” was used for Gemini. This second prompt was submitted following the first prompt and LLM response for each demonym. The use of a follow-up prompt was motivated by our observation that ChatGPT frequently suggested follow-up prompts relating to genetics when responding to the first prompt’s history queries. Prior to analysis, responses from ChatGPT were inspected for duplicate source sections, as the platform inconsistently produced both a list of URLs and an inline-cited reference list within the same response. Responses were then analyzed using a Python script available at https://github.com/diazale/wikigenetics. For each response, we recorded the number of times Wikipedia URLs and genetics keywords appeared in the cited sources and output body, respectively. The genetics keywords studied were identical to those defined for the Wikipedia corpus analysis. We provide the responses and our data as supplementary zip files.

### Grokipedia

We downloaded the raw HTML from each Grokipedia page using cURL and parsed the text using a custom script available at https://github.com/diazale/wikigenetics. All pages were downloaded between March 24 and April 4, 2026. Unlike Wikipedia, Grokipedia does not allow edits by users but may implement edit suggestions. While Grokipedia pages have a tab to show their edit history, these are not consistently available and frequently display “Failed to load edits.” We have made the HTML and parsed text for our Grokipedia page downloads available at 10.5281/zenodo.21537405. Of the 137 demonyms that had pages on Wikipedia, 130 also had pages on Grokipedia. The seven demonyms that had pages on Wikipedia but not Grokipedia were: Antiguans and Barbudans, Croatians, Dominicans, Somalians, Serbs, Qataris, and Paraguayans. There did not appear to be a systematic pattern in these missing pages.

When calculating text length, we defined it as character length (including spaces) after references have been removed. This is consistent with the text length used in our Wikipedia analyses. To detect genetics keywords, we used the same keyword patterns as in the Wikipedia analyses. To compare full-page text between Grokipedia and Wikipedia, we used the term frequency-inverse document frequency (TF-IDF) measure using cosine similarity as our distance measure. To vectorize our text, we use the TfidfVectorizer function in the Python library *scikit-learn*, computing

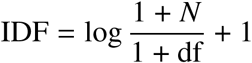

and

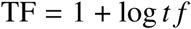

TF is the relative frequency of a term within a document and IDF is the smooth inverse document frequency. The final TF-IDF score is given by the cosine similarity of the L2-normalized TF × IDF vectors.

To rank the top terms, we use the same tokens (words) as in the TF-IDF score and calculate a log-odds of the two pages:

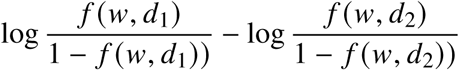

where *f* (*w*, *d_i_*) is the frequency of the word in document *i*. A high positive value indicates a word is more prevalent in document 1, and a high negative score indicates a word is more prevalent in document 2.

## Supplementary Text

### Case study comparisons to Grokipedia

Grokipedia’s citation have been found to be of lower quality than on Wikipedia (*47*). We identified many instances of hallucinated or misrepresented citations, or of low quality citations. The page “Black people” cites racialist blogs, StackExchange, subreddits, encyclopedia.com, and hereditarian writers (e.g. Harpending, Jensen, Lynn, Cochran) which are generally not considered reliable sources by the Wikipedia community. Grokipedia text also misinterprets academic sources. For example, the opening sentence of “Black people” cites Rosenberg et al. (*62*) in defining “Black people” as a cluster in STRUCTURE, but this is never mentioned in the Rosenberg paper.

For each case study page, we compared the Wikipedia and Grokipedia versions using TF-IDF and a log-odds ranking of terms (see Methods—Grokipedia).

For “Ashkenazi Jews”, the keywords “genetic” and “admixture” appear in the introduction. The page had a total of 183 genetics keywords, with genetics sections comprising an adjusted 21.5% of page text. The top term on Grokipedia was “IQ”, reflecting a substantial amount of text written about higher-than-average IQ scores and a putative genetic basis; the top term on Wikipedia was “people”. Numerous top-ranked terms on Grokipedia were related to genetics (e.g. “selection”, “bottlenecks”, “analyses”) or to intelligence measures.

For “White people”, the keywords “genetic” and “admixture” appear in the first sentence. There were 247 genetics keywords, and genetics sections are 7.8% of all text. The top term on Grokipedia was “steppe”, referring to ancient DNA and haplogroup studies; the top term on Wikipedia was “nineteenth”. Many genetics-related terms on Grokipedia related to ancient DNA studies (e.g. “hunter gatherer”, “migrations”, “neolithic”, “bronze age”).

For “Black people”, the keywords “genetic” and “genomic” appeared in the first sentence. The top term on Grokipedia was “rates”, reflecting a strong focus on negative socioeconomic outcomes (e.g. “crime rates”, “homicide rates”). The top term on Wikipedia was “aboriginal”, reflecting a section written about Indigenous Australians, who are not mentioned in Grokipedia. Genetics and genetics-adjacent terms strongly present in Grokipedia include “analyses”, “variants”, “structure”, and “samples”.

### Related pages and categories

A number of pages and categories were related to the project but deemed out of scope and not included in the corpus. These include the category “Modern human genetic history” (36 pages plus 10 subcategories), “Genetics by ethnicity” (21 pages), “Jewish genetics” (9 pages), “Genetics by country” (15 pages), and “Human haplogroups” (which includes 55 mtDNA haplogroup pages and 119 Y-DNA haplogroup pages). We additionally identified 29 pages focusing on the genetics or genetic histories of populations (“genetic history” or “genetic studies” pages), which collectively received over 1.175 million views in 2025, and at least 11 “origin hypotheses” pages for extant groups, nine of which contain genetics sections and collectively received over 530,000 views in 2025.

### Wikipedia editing policies and guidelines

Wikipedia is written by volunteers working in good faith to create a free encyclopedia, collaboratively writing comprehensive articles on topics for which reliable sources exist. The site has many policies and guidelines, the most salient of which are: pages should have a neutral point of view, giving due weight to majority and significant minority viewpoints and not giving undue weight to fringe views (*63*); all material should be verifiable based on published reliable sources (*64*); original research is forbidden, as material must be directly and explicitly supported by cited reliable sources with no original synthesis (*65*); and reliable sources should have a reputation for fact-checking and accuracy, with reviews given more weight than primary research for scientific publications (*66*). The standards for “due weight” and “reliable sources” have evolved over time. For example, in a centralized discussion from April 2020, the community came to a consensus that the idea that IQ differences across racial groups were rooted in genetic differences represented a fringe scientific view (*40*).

Consensus is the fundamental method of decision-making on Wikipedia (*67*). Users may propose changes to pages, policies, or guidelines, and then discuss it amongst themselves or request feedback from the broader community. The Wikipedia community itself has had multiple discussions about whether and how to include population genetics research in pages about ethnic groups, including at least 10 community discussions between 2009 and 2018 (*39*). In one discussion from 2018, users from a project dedicated to ethnic groups discussed whether such pages should have genetics sections (*68*). In favour of removing the sections, editors stated concerns over eugenics and that genetics do not and should not define ethnic groups; that genetics sections draw an outsized amount of energy and attention compared to, e.g., language or history; that most Wikipedia editors lack the technical expertise to interpret genetic studies; and that many users who write about genetics are politically motivated. In opposition to removing the sections, editors stated that genetics studies are peer-reviewed scientific content and reliable sources; that people are interested in genetic studies and should be pointed to useful sources; that pages could be rewritten to better represent scientific viewpoints and to counteract concerns over racism; and that academic sources state there is a relation between population genetics, history, ethnicity, and identity. The discussion was ultimately closed with no consensus.

### Examples of topics in plain text

All excerpts are taken from the most recent publicly-available revision as of December 31, 2025. Scoring high in this context means the the page had one of the highest counts of terms related to a topic. This may be the consequence of repeating words (e.g. “haplogroup”). The hierarchical clustering was done on binary indicators for each topic for each paragraph; multiple mentions of the same term within the same paragraph did not impact the clustering.

**“Kadazan-Dusun”** (revision 1324423170). Scored high on “population genetics methods”. Samples paragraph:

> Maternal or Matrilineal Studies Using mitochondrial DNA (mtDNA), is a test used to explore genetic ancestry from the mother using mtDNA that is obtained from outside of a nucleus cell that isn’t contaminated by the presence of Y-chromosome. According to a study published in 2014, by Kee Boon Pin on 150 volunteers from the Kadazandusun people all over the Sabah region, the Kadazandusun people belongs to 9 mtDNA Haplogroups (subjected to the numbers and types of samples involved in the study), with Haplogroup M being the highest frequency, where it represents (60/150 40%) of all maternal lineages. Followed by Haplogroup R (26/150 17.33%), Haplogroup E (22/150 14.67%), Haplogroup B (20/150 13.33%), Haplogroup D (9/150 6%), Haplogroup JT (6/150 4%), Haplogroup N (4/150 2.67%), Haplogroup F (2/150 1.33) and Haplogroup HV (1/150 0.67%)

**“Greeks”** (revision 1329183413). Scored high on “statistics & clustering”.

> Genetic studies using multiple autosomal, Y-DNA, and mtDNA markers, show that Greeks share similar backgrounds as the rest of the Europeans and especially Southern Europeans (Italians and Balkan populations such as Albanians, Slavic Macedonians and Romanians). A study in 2008 showed that Greeks are genetically closest to Italians and Romanians and another 2008 study showed that they are close to Italians, Albanians, Romanians and southern Balkan Slavs such as Slavic Macedonians and Bulgarians. A 2003 study showed that Greeks cluster with other South European (mainly Italians) and North-European populations and are close to the Basques, and FST distances showed that they group with other European and Mediterranean populations, especially with Italians (−0.0001) and Tuscans (0.0005). A study in 2008 showed that Greek regional samples from the mainland cluster with those from the Balkans, principally Albanians while Cretan Greeks cluster with the central Mediterranean and Eastern Mediterranean samples.

**“Hungarians”** (revision 1329183413). Scored high on “ancestry & origins”.

> Modern Hungarians stand out as linguistically isolated in Europe, despite their genetic similarity to the surrounding populations. The population of the Carpathian Basin has the common European gene-pool which formed in the Bronze Age through the admixture of three sources: Western Hunter-Gatherers, who were the first Homo sapiens appearing in Paleolithic Europe, Neolithic farmers originating from Anatolia, and Yamnaya steppe migrants that arrived in the late Neolithic to early Bronze Age. This common European gene pool in the Carpathian Basin, has been overlaid by migration waves originating from the east since the Iron Age. According to genetic studies, the Carpathian Basin was continuously inhabited from at least the Bronze Age.

**“Lumad”** (revision 1323517735). Scored high on “phenotypes”.

> They are traditionally hunter-gatherers and consume a wide variety of wild plants, herbs, insects, and animals from tropical rainforests. The Mamanwa are categorized as having the “negrito” phenotype: dark skin, kinky hair, and short stature. The origins of this phenotype (found in the Agta, Ati, and Aeta tribes in the Philippines) are a continued topic of debate, with recent evidence suggesting that the phenotype convergently evolved in several areas of southeast Asia.

**“Japanese Americans”** (revision 1329911939). Scored high on “medical genetics”.

> Studies have looked into the risk factors that are more prone to Japanese Americans, specifically in hundreds of family generations of Nisei (The generation of people born in North America, Philippines, Latin America, Hawaii, or any country outside Japan either to at least one Issei or one non-immigrant Japanese parent) second-generation pro-bands (A person serving as the starting point for the genetic study of a family, used in medicine and psychiatry). The risk factors for genetic diseases in Japanese Americans include coronary heart disease and diabetes. One study, called the Japanese American Community Diabetes Study that started in 1994 and went through 2003, involved the pro-bands taking part to test whether the increased risk of diabetes among Japanese Americans is due to the effects of Japanese Americans having a more westernized lifestyle due to the many differences between the United States of America and Japan. One of the main goals of the study was to create an archive of DNA samples which could be used to identify which diseases are more susceptible in Japanese Americans.

**“Malaysian Malays”** (revision 1329528430). Scored high on “group comparison”.

> Within the Malay Peninsula itself, the Malays are differentiated genetically into distinct clusters between the northern part of the Malay Peninsula and the south. SNP analyses of five of their sub-ethnic groups show that Melayu Kelantan and Melayu Kedah (both in the northern Malay Peninsula) are closely related to each other as well as to Melayu Patani, but are distinct from Melayu Minang (western), Melayu Jawa and Melayu Bugis (both southern). The Melayu Minang, Melayu Jawa and Melayu Bugis people show close relationship with the people of Indonesia, evidence of their shared common ancestry with these people. However, Melayu Minang are closer genetically to Melayu Kelantan and Melayu Kedah than they are to Melayu Jawa. Among the Melayu Kelantan and Melayu Kedah populations, there are significant Indian components, in particular from the Telugus and Marathis. The Melayu Kedah and Melayu Kelantan also have closer genetic relationship to the two subgroups of the Orang Asli Semang, Jahai and Kensiu, than other Malay groups. Four of the Malay sub-ethnic groups in this study (the exception being Melayu Bugis, who are related to the people of Sulawesi, Indonesia) also show genetic similarity to the Proto-Malay Temuan people with possible admixture to the Jawa populations and the Wa people of Yunnan, China.

**“African Pygmies”** (revision 1314287229). Scored high on “subsistence mode”.

> The mitochondrial DNA and Y-Chromosome haplogroups found in the ancient Shum Laka foragers were Sub-Saharan African haplogroups. Two earlier Shum Laka foragers were of haplogroup L0a2a1 – broadly distributed throughout modern African populations – and two later Shum Laka foragers were of haplogroup L1c2a1b – distributed among both modern West and Central African agriculturalists and hunter-gatherers. One earlier Shum Laka forager was of haplogroup B and one later Shum Laka forager haplogroup B2b, which, together, as macrohaplogroup B, is distributed among modern Central African hunter-gatherers (e.g., Baka, Bakola, Biaka, Bedzan).

**“Puerto Ricans”** (revision 1329310619). Scored high on “admixture and mixing”.

**“Indigenous Australians”** (revision 1327604145). Scored high on “archaic and deep lineage”.

> Genetic studies have suggested that Aboriginal Australians largely descended from an Eastern Eurasian population wave during the Initial Upper Paleolithic, and are most closely related to other Oceanians, such as Melanesians. The Aboriginal Australians also show affinity to Ancient Ancestral South Indians, the Andamanese people, as well as to East Asian peoples. Phylogenetic data suggests that an early initial eastern non-African (ENA) or East-Eurasian meta-population trifurcated, and gave rise to Australasians (Oceanians), the Ancient Ancestral South Indians, Andamanese and the East/Southeast Asian lineage including the ancestors of Native Americans, although Papuans may have also received some geneflow from an earlier group (xOOA) as well, around 2%, next to additional archaic admixture in the Sahul region.

**“Aari people”** (revision 1260272339). Scored high on “social structure”.

> The Ari peoples of Ethiopia comprise different occupational groups and their society is socially divided and stratified according to each Aari individual’s respective occupation. The lower castes of the society is composed of potters, tanners and blacksmiths and collectively named as mana in the Aari language. Blacksmiths (faka mana)i who also do woodworking are marginalized and occupy an inferior position to tanners and potters (tila mana). Kantsa is the name given to the agriculturalist caste which holds a privileged position in the society. Intermarriage between mana and katsa is forbidden and considered as taboo according to Ari customs. The occupational segregation and caste-based endogamy practiced among the Ari have been revealed by advances in archaeogenetics to be one the oldest continuous caste systems in existence. After the introduction of Christianity the social division between Christian Aari belonging to differing castes have reported to become less important. More of the societies make agriculture their livelihood, and most of them practice mixed farming.

**“Mokshas”** (revision 1330183261). Scored high on “race classification”.

> K.Yu. Mark distinguishes the Sub-Ural and North Pontic type among the Mokshans, and among the Erzyans — the Sura type, close to the Atlanto-Baltic anthropological type. Anthropologist Tatyana Ivanovna Alekseeva argued that in the Mokshans, compared to the Erzyans, the features of Southern Europeans are more noticeably manifested, and she attributes the Erzyans more to the circle of Northern Europeans. V.E. Deryabin noted that the Moksha people have an Eastern European base, modified by a Pontic anthropological component in combination with a slight Uraloid admixture. According to the publication of the Russian Academy of Sciences (2000) edited by Aleksandr Zubov, the Erzyans belong to the White Sea-Baltic version of the Caucasian race, which is represented, in addition to the Erzyans, by the majority of the Baltic Finnish-speaking peoples and part of the Komi-Zyryans. The Mokshas belong to the Ural race, within which the Mokshas are classified as the Sub-Ural subtype. The anthropological difference between the Erzyans and Mokshas, who are basically Caucasian race and subethnic groups of one of the most anthropologically homogeneous peoples, lies, in particular, in the fact that the Atlantic and North Pontic types are to some extent superimposed on the White Sea-Baltic basis of the Mordovians. The first type is represented predominantly among the Erzyans, the second — among the Mokshans, although both types are present in both categories of the population. Anthropologically, Moksha was formed as a result of the mixing of various types (White Sea, Pontic, East Baltic) of the Caucasian race.

**“Indigenous peoples of Mexico”** (revision 1329359626). Scored high on “identity & ethnicity”.

> The Indigenous groups within what is now Mexico are genetically distinct from each other. The genetic differences between geographically separated Indigenous groups (e.g., between Indigenous people living in the Yucatán Peninsula compared to Indigenous people living in western Mexico) can be as large as the genetic differences seen between a European and an East Asian person.

**“Turkic peoples”** (revision 1330294826). Scored high on “language & linguistics”.

> Many vastly differing ethnic groups have throughout history become part of the Turkic peoples through language shift, acculturation, conquest, intermixing, adoption, and religious conversion. Nevertheless, Turkic peoples share, to varying degrees, non-linguistic characteristics like cultural traits, ancestry from a common gene pool, and historical experiences. Some of the most notable modern Turkic ethnic groups include the Altai people, Azerbaijanis, Chuvash people, Gagauz people, Kazakhs, Kyrgyz people, Turkmens, Turkish people, Tuvans, Uyghurs, Uzbeks, and Yakuts.

**“Paranakan Chinese”** (revision 1328864737). Scored high on “religion”.

> Many Peranakan in Java, Indonesia are descendants of non-Muslim Chinese men who married abangan Javanese Muslim women. Most of the Chinese men did not convert to Islam since their Javanese wives did not ask them to, but a minority of Javanese women asked them to convert so a Chinese Muslim community made out of converts appeared among the Javanese. In the late half of the 19th century, Javanese Muslims became more adherent to Islamic rules due to going on hajj and more Arabs arriving in Java, ordering circumcision for converts. The Batavian Muslims in the 19th century completely absorbed the converted Chinese Muslims who originally had their own separate kapitan and community in the late 18th century. The remaining commoner non-Muslim Chinese Peranakans descended from Chinese men and Javanese Muslim women generally stopped marrying Javanese and the elite Peranakans stopped marrying Javanese completely and instead started only marrying fellow Chinese Peranakans in the 19th century, as they realized they might get absorbed by the Muslims. DNA tests done on Chinese Peranakan in Singapore showed that those Peranakan who are mixed with Malays are mostly of paternal Han Chinese descent and of maternal Malay descent. Peranakans in Malaysia and Singapore formed when non-Muslim Chinese men were able to marry Malay Muslim women a long time ago without converting to Islam. This is no longer the case in modern times where anyone who marries Malay women is required to convert to Islam.

---

**Table S1:** Substrings used to assign a context to each genetics keyword. To determine the context in which a genetics keyword appeared in a demonym page, we identified the section of the page that contained the keyword and assigned one or more contexts by searching for the substrings below within the section title. Keywords can appear in multiple contexts, e.g. a section titled “Genetic origins” is assigned both “Biology” and “Origins”.

| Context | Substrings |
| --- | --- |
| Biology | dna, genet, genom, haplo, autosom, structure, biolog, pheno, physical |
| Origins | origin, ancestr, ethnogen, arriv, identi, lineage |
| Culture | culture, language, history, period |
| Reference | refer, citation, cited, further, see also, source, publication, external |
| Race, ethnicity, nationality, citizenship | ethnic, race, racial, nationality, citizenship |
| Introduction section | Any substrings found before a section has been delineated |

**Figure S1:**
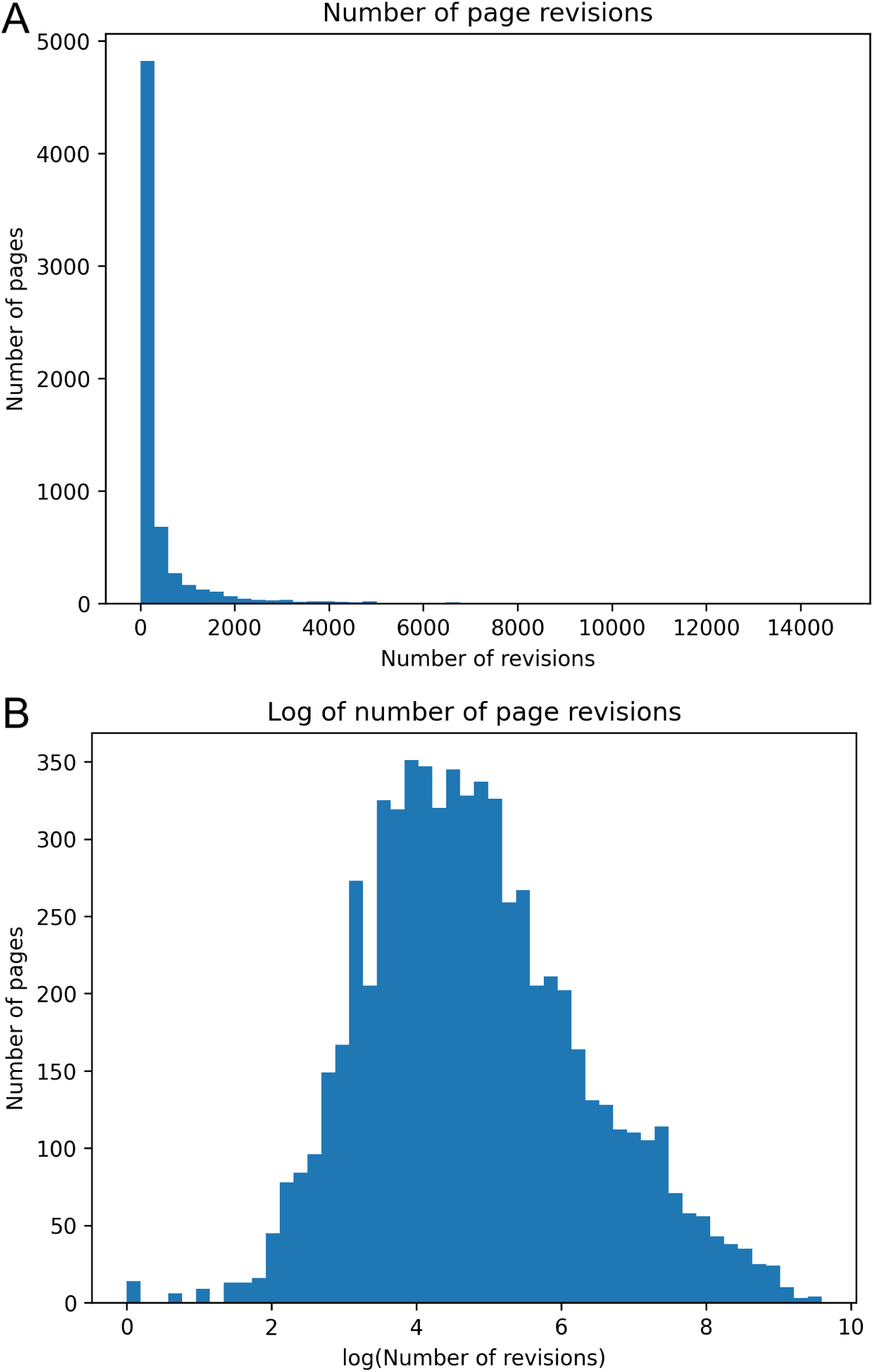
The distribution of edits to pages is highly skewed. **(A)** Histogram of the number of edits to every page in the corpus. **(B)** Histogram of the log of the number of edits to every page in the corpus. The vast majority of pages receive relatively few edits (median = 107).

**Figure S2:**
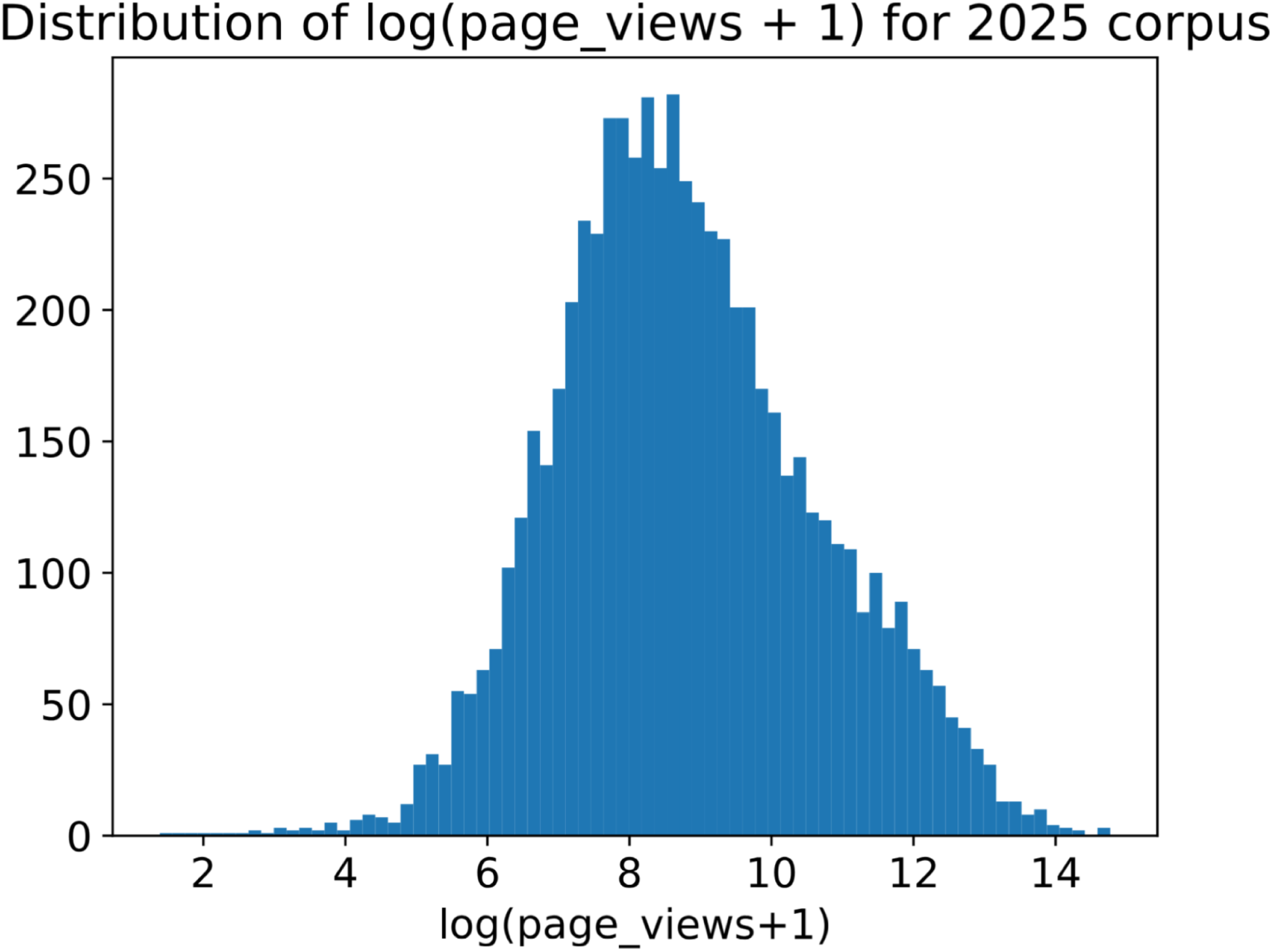
Views of Wikipedia pages in the corpus are approximately log-normally distributed. The distribution of the log of (total views + 1) from January 1, 2025 to December 31, 2025 for all pages in the analysis corpus. Views are users only and exclude automated agents. The mean value is 8.800 and the standard deviation is 1.829.

**Figure S3:**
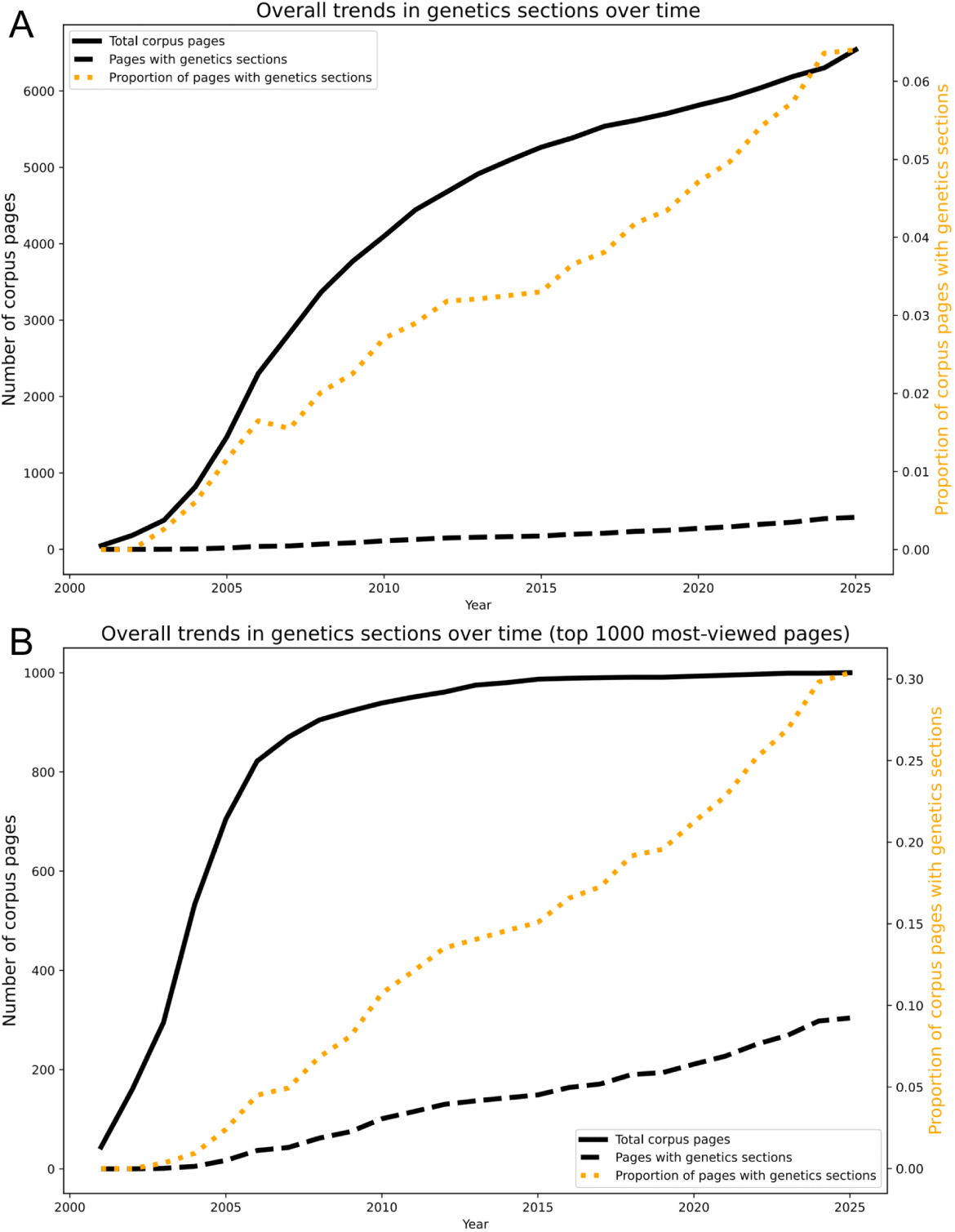
The number of pages with genetics sections increased steadily over time, especially among the most popular pages. A time series of the number of corpus pages with genetics sections. The solid line indicates the number of corpus pages created at that time. The dashed line indicates the number of corpus pages with genetics sections. **(A)** All corpus pages. **(B)** The top-1000 most viewed corpus pages.

**Figure S4:**
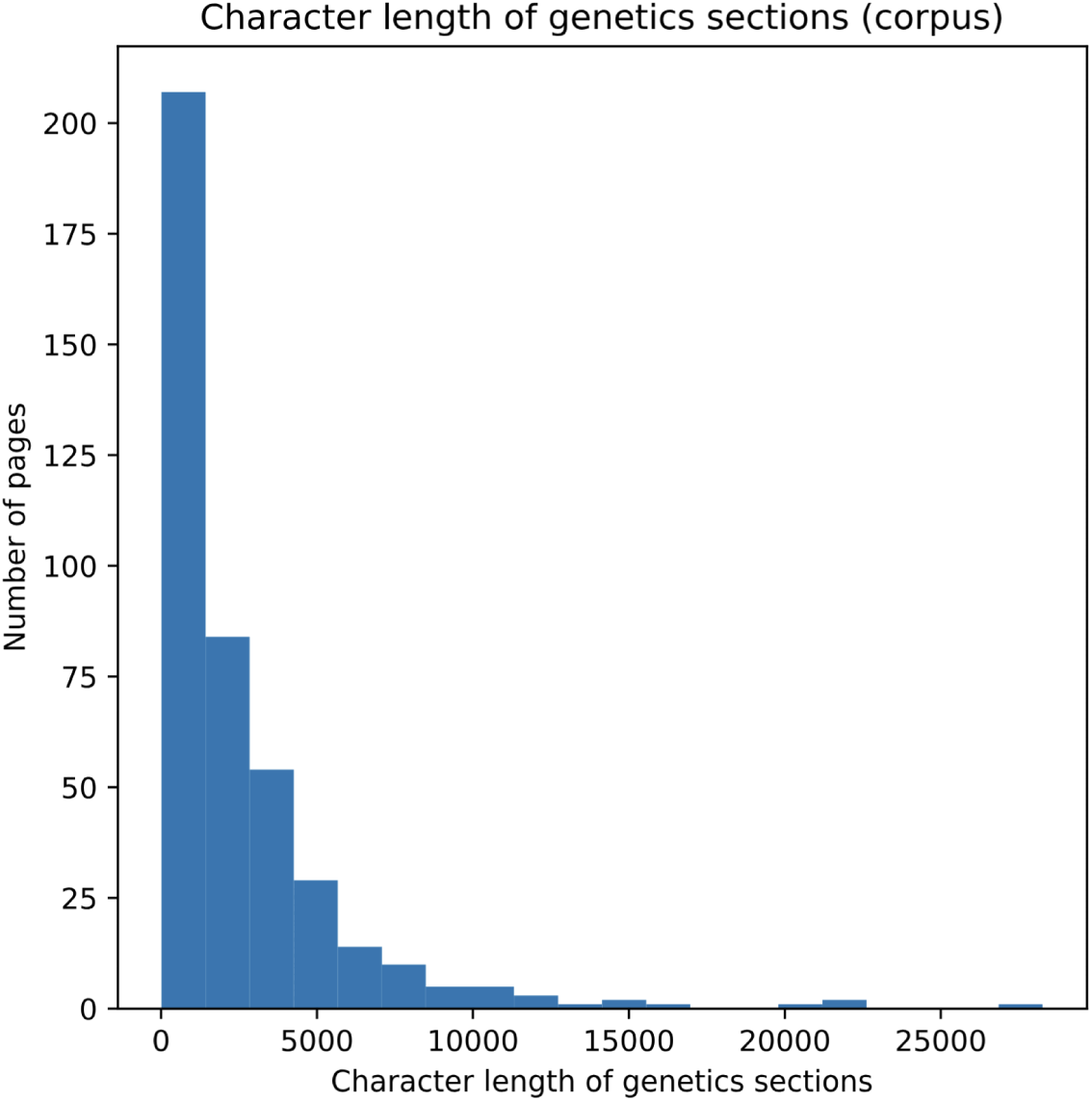
Genetics sections have a skewed distribution of lengths. The average genetics section has a length of 2,588 characters and takes up 11.1% of the page’s text. Length is calculated as the number of characters, including spaces, in plain text.

**Figure S5:**
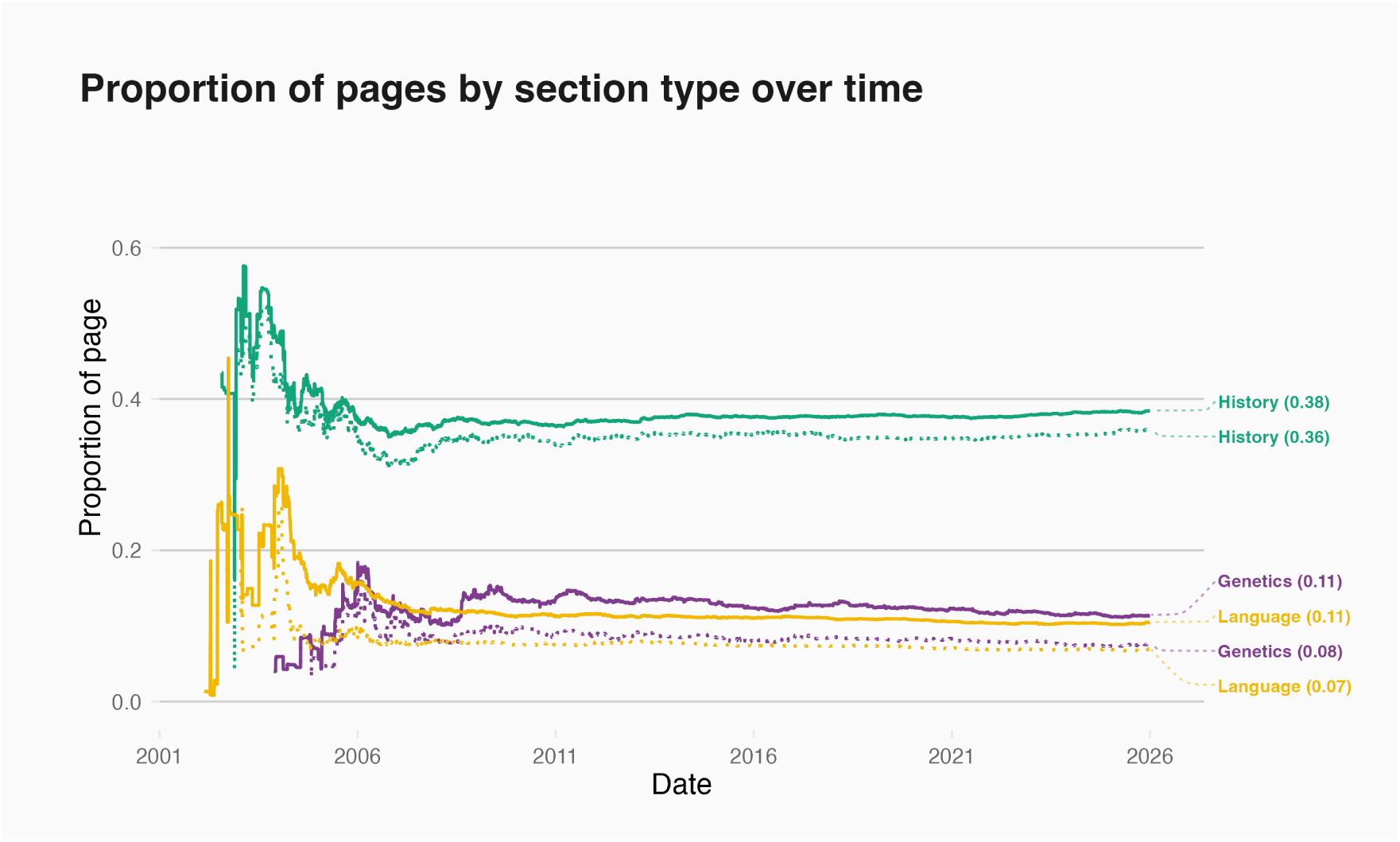
Genetics sections in our corpus pages take up a mean of 11% of a page’s text. Comparing the proportion of a page’s plain text taken up by different types of common sections, genetics sections are comparable to language sections in size, but smaller than history sections. While these proportions vary widely between pages, the average sizes have stabilized over time. Solid lines indicate the mean, dashed lines indicate the median.

**Figure S6:**
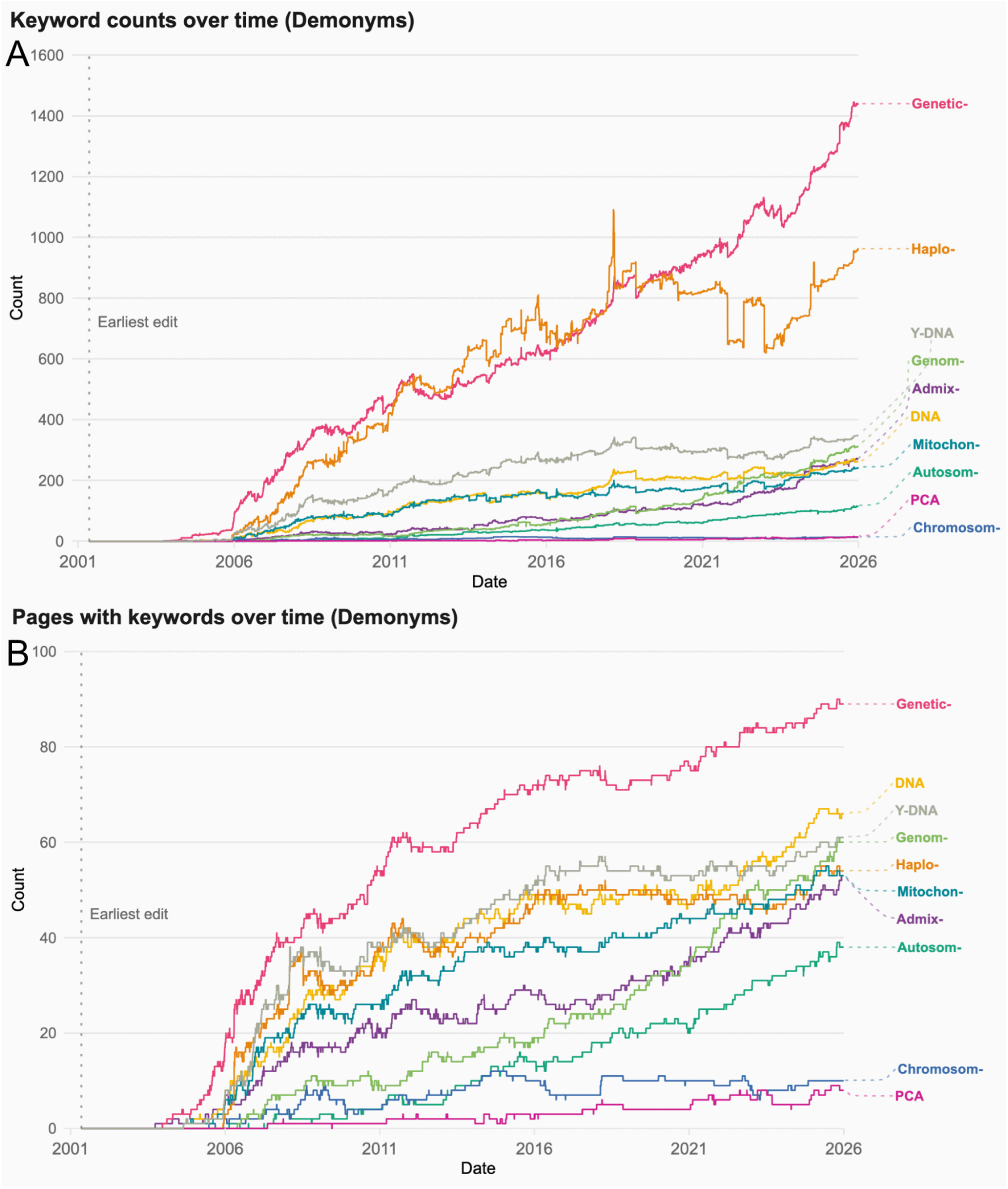
Genetics keywords in Wikipedia pages about demonyms have increased steadily over time. **(A)** The number of keywords used across all 137 demonym pages. **(B)** The number of demonym pages in which each genetics keyword appears.

**Figure S7:**
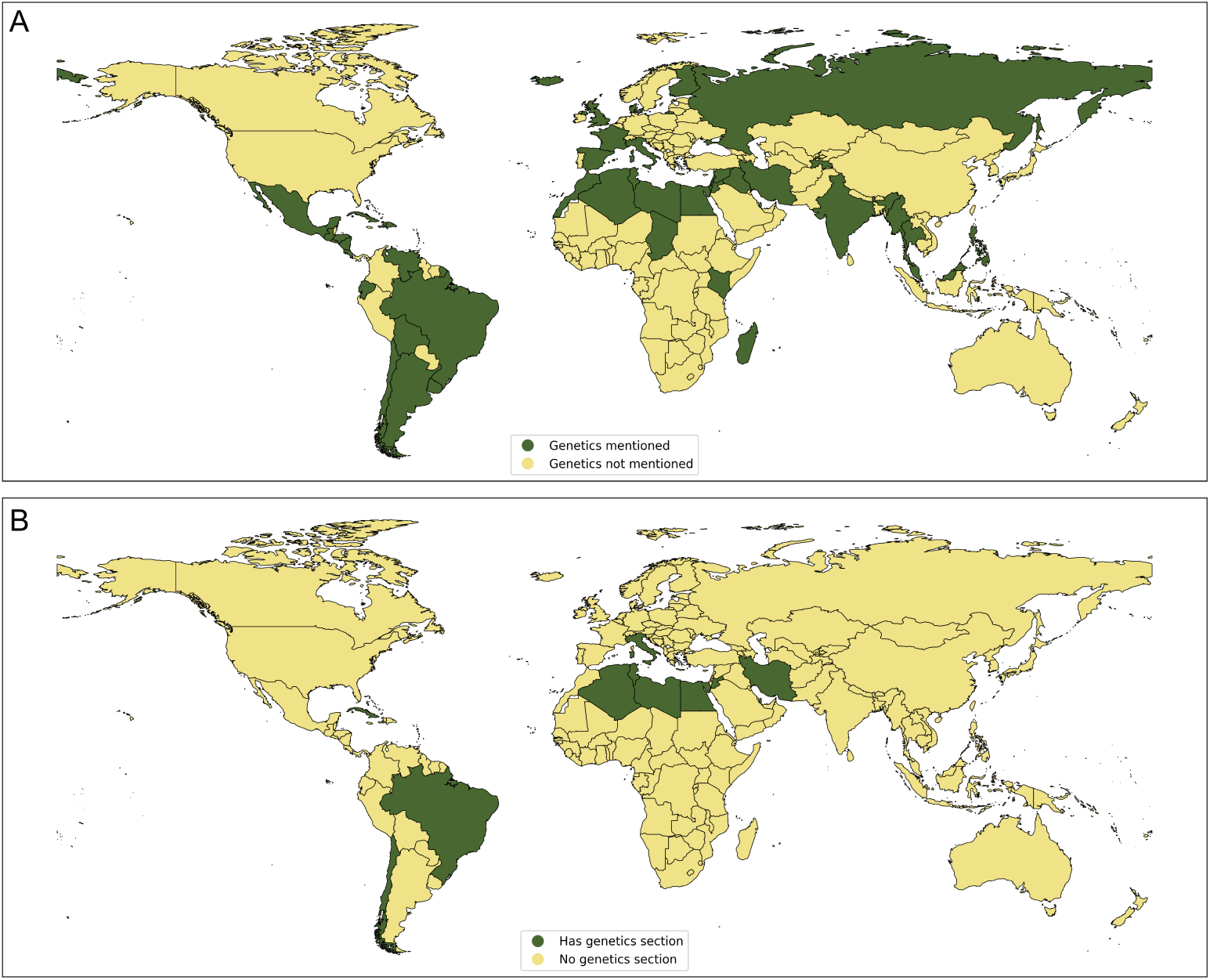
Map of presence of genetics keywords and sections among demographics pages. There are 197 pages about the demographics of countries, whose titles follow the format “Demographics of [COUNTRY]” (e.g. “Demographics of Kenya”). (**A**) 55 (27.9%) pages had genetics keywords. Countries in green have a demographics page with genetics keywords; countries in yellow do not. (**B**) 13 (6.6%) pages had genetics sections. Countries in green have a demographics page with a genetics section; countries in yellow do not.

**Figure S8:**
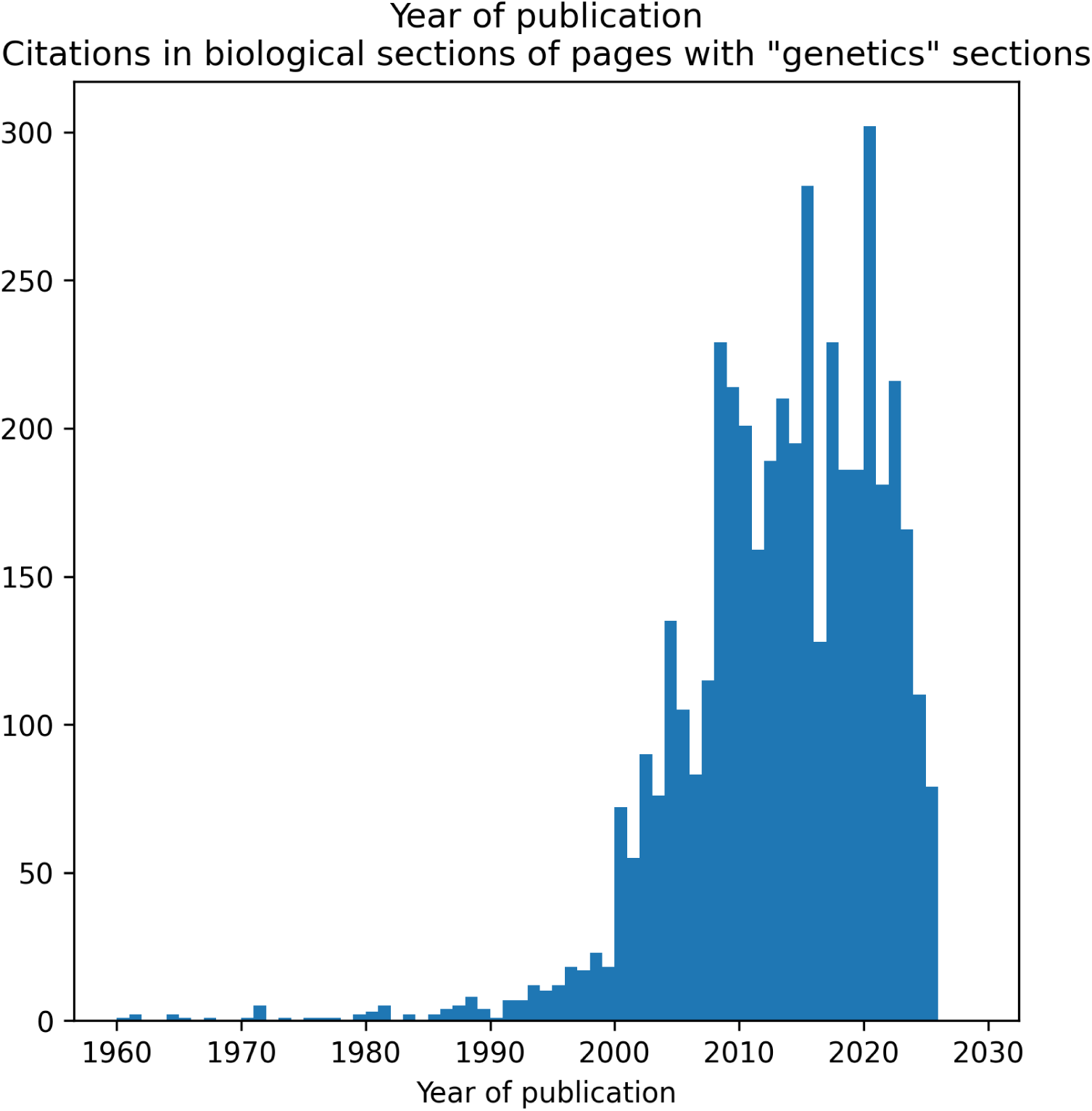
The vast majority of citations have been published since the release of the first draft of the human genome. Citations within genetics sections of pages in the corpus are relatively recent, with the vast majority (3,245; 91.7%) having been published on or after 2003.

**Figure S9:**
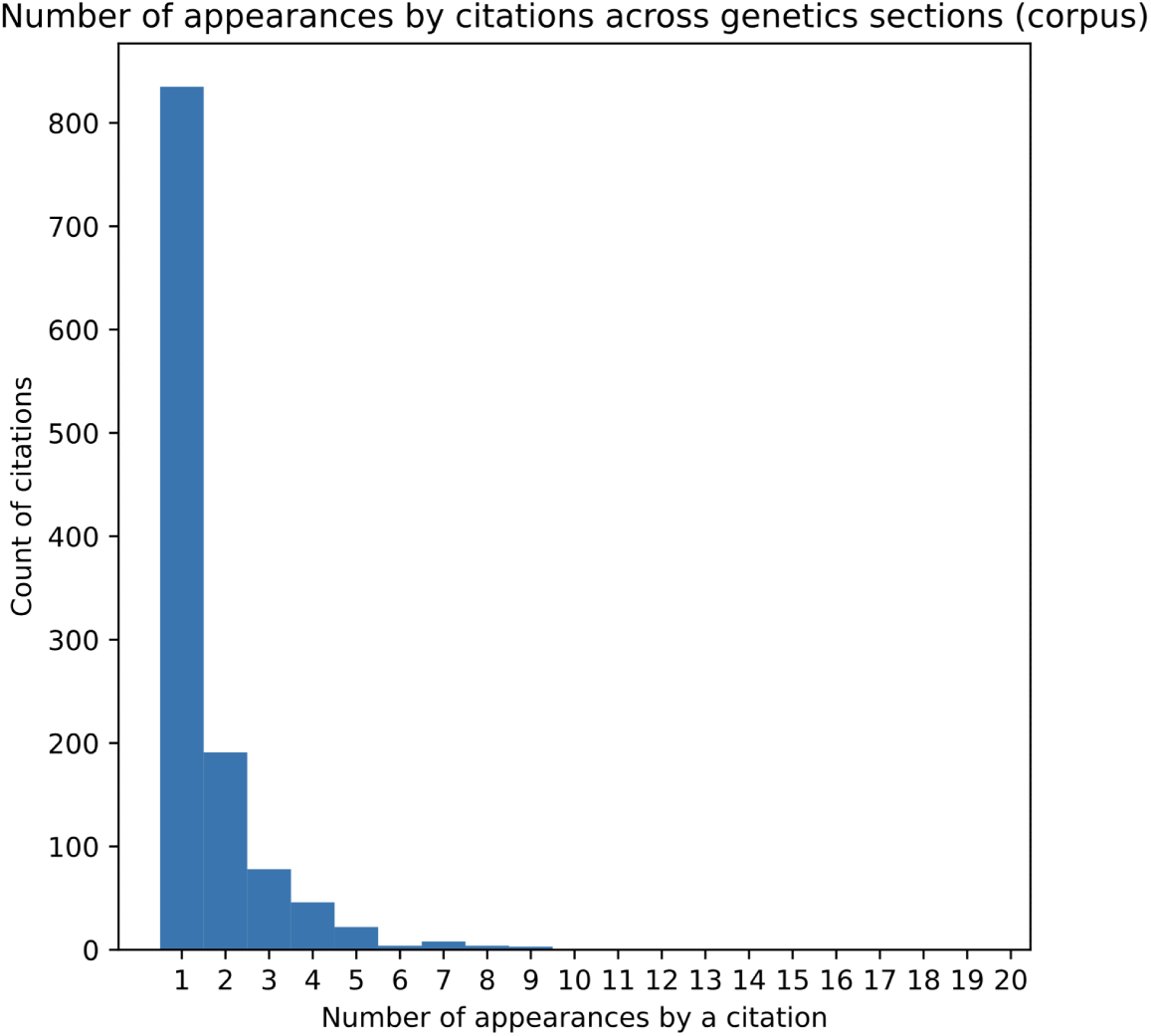
The vast majority of citations within genetics sections are only used in one page. We identified all citations used in genetics sections in pages within the corpus (419 of 6,541 corpus pages had a genetics section). From these citations, we extracted 1,942 digital object identifiers (DOIs), of which 1,195 were unique. The vast majority (835; 69.9%) were only cited in one page (see also table S5).

**Figure S10:**
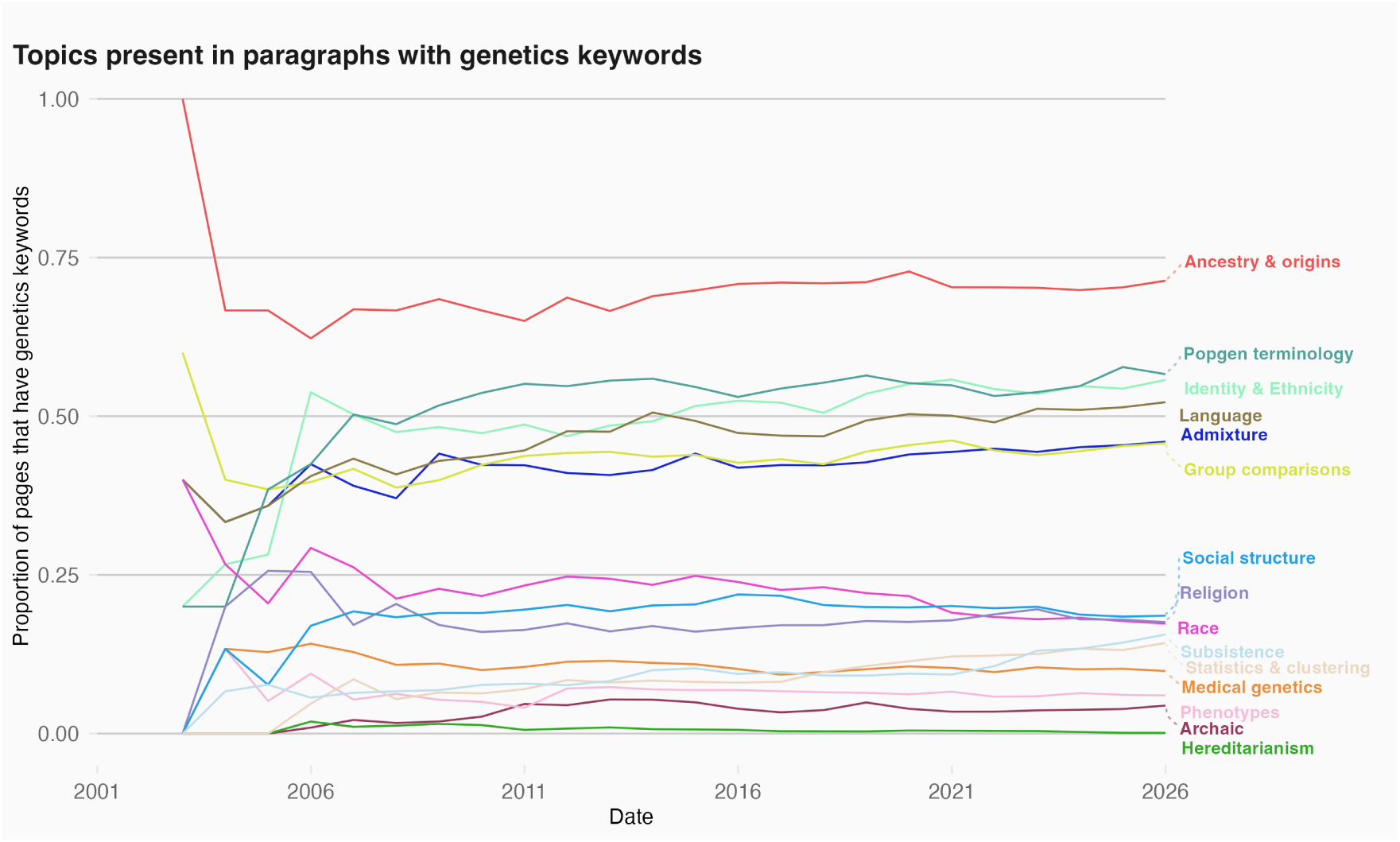
Topics in the corpus text over time. As of December 31, 2025, the most common topics in prose that uses genetics keywords are, in order: ancestry & origins; population genetics methods; identity & ethnicity; language; admixture, group comparisons; social structure; religion; race; subsistence mode; statistics & clustering; medical genetics; phenotypes; archaic ancestry; and hereditarianism. These values represent the proportion of corpus pages that have genetics keywords as of December 31 of each year.

**Figure S11:**
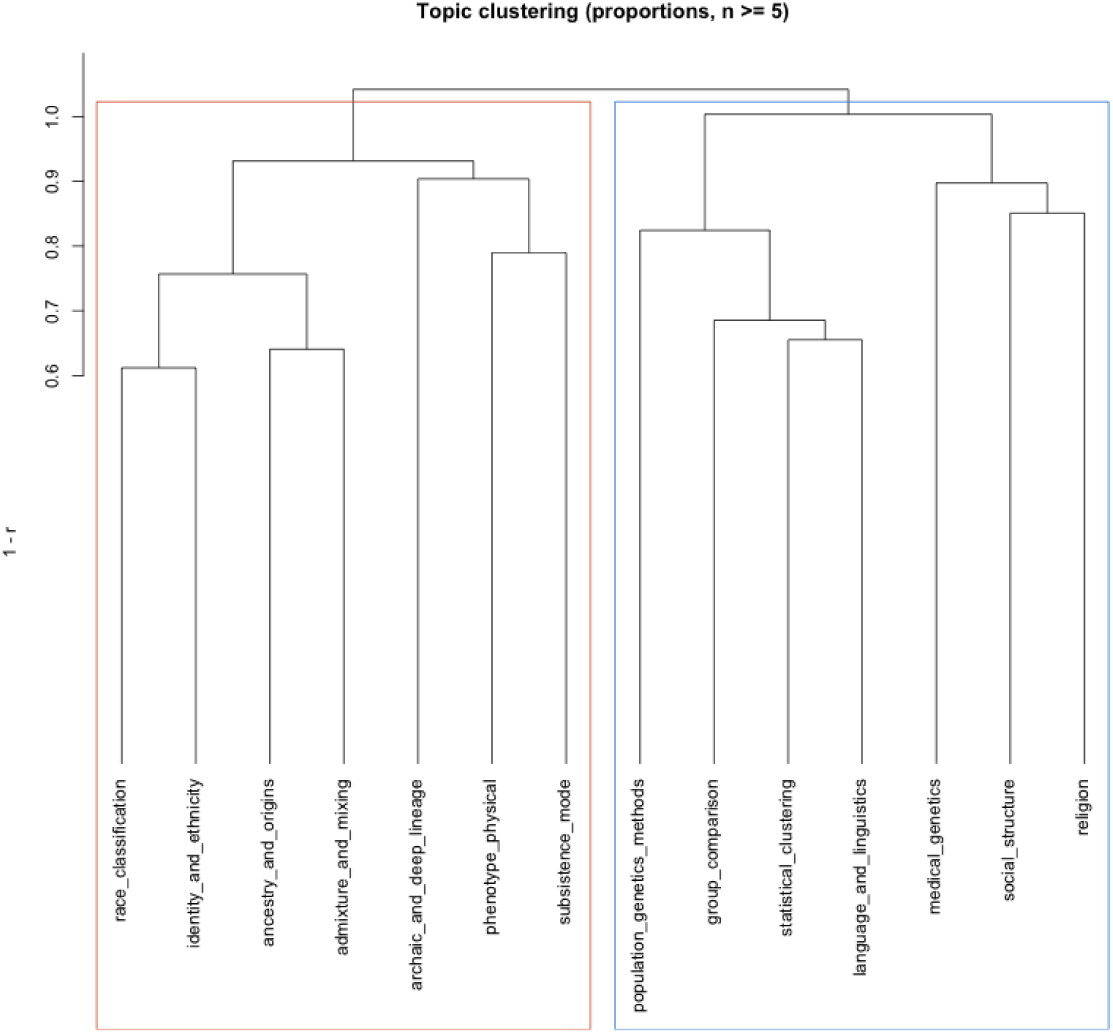
Hierarchical clustering of corpus text topics. We identified every page in the corpus that had at least five paragraphs of plain text that had a genetics term (187 pages) and carried out hierarchical clustering on topics within those paragraphs (see Methods—Text topic analysis for full details). We identified two broad clusters: those related to the topics of “race”, “identity & ethnicity”, “ancestry & origins”, “admixture”; and those related to “population genetics methods”, “group comparison”, “statistics & clustering”, “language & linguistics”.

**Figure S12:**
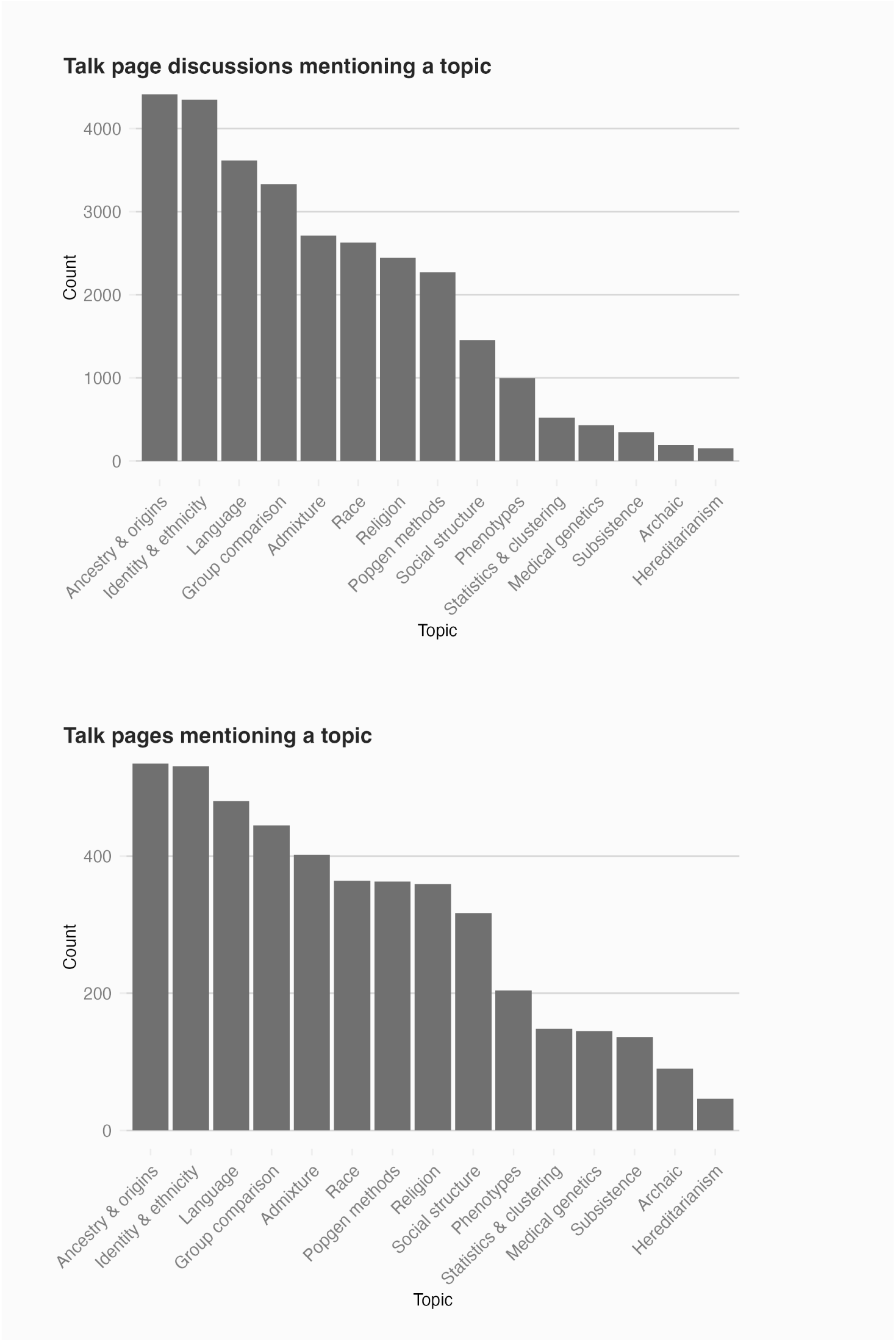
Topics analyzed in talk pages across corpus pages.

**Figure S13:**
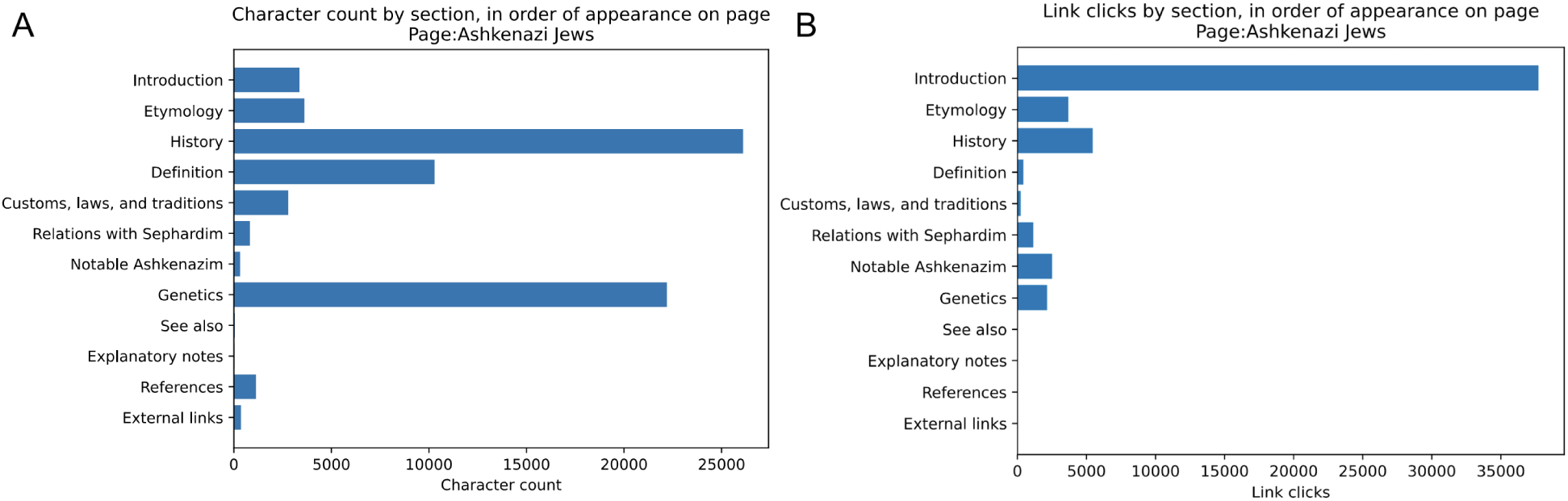
The “Genetics” section of the page “Ashkenazi Jews” receives an outsized amount of attention when measured by both section length and links clicked. **(A)** The character count of each section of the page “Ashkenazi Jews”, with the sections listed from top to bottom. The two longest sections are “History” (26,102 characters; 36.7% of total page text) and “Genetics” (22,201; 31.2%). **(B)** The estimated number of times an internal link within a section was clicked for the first time links appear in a page. The most-clicked links are usually at the top of a page. Despite being at the bottom of the page, the “Genetics” section received at least 2,150 clicks (7.6% of all clicks within the page), suggesting it is being read more often than expected.

**Figure S14:**
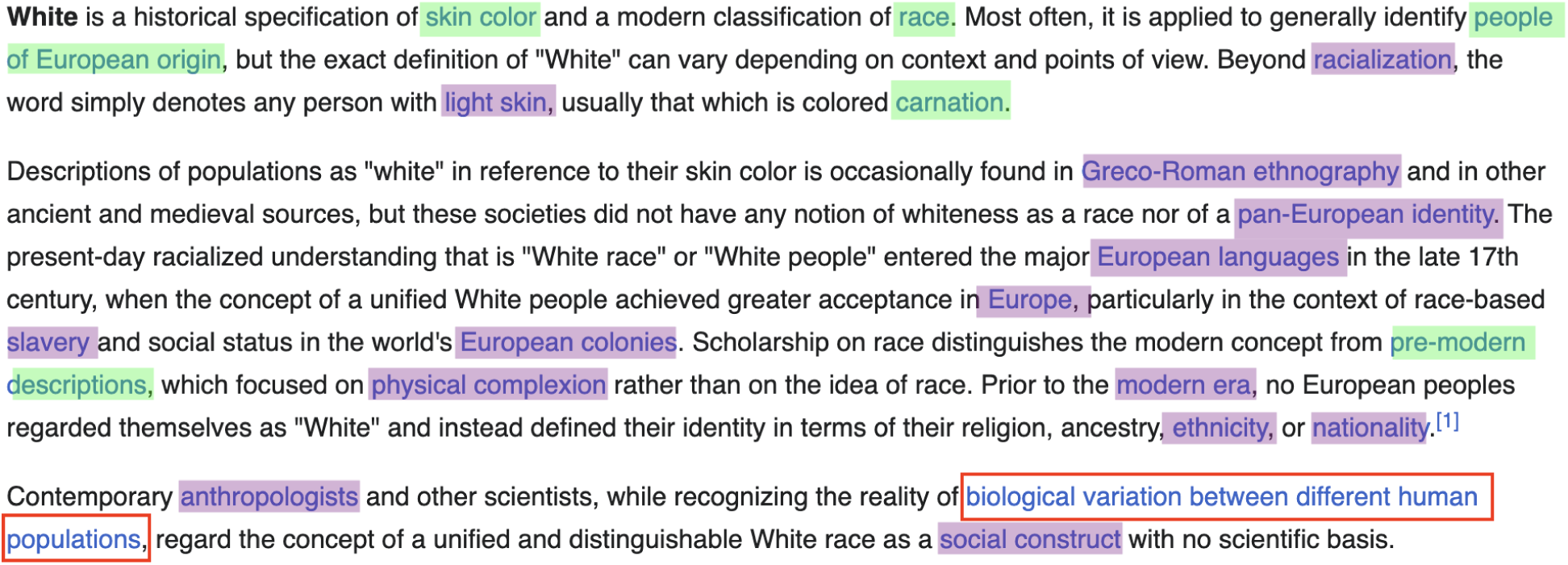
A screenshot of the introduction section of the page “White people” from December 2025. Only one link (“Race and genetics”, highlighted in red box) related to genetics appeared in the page “White people”. Links highlighted in green have more clicks than “Race and genetics”, while those highlighted in purple have fewer.

**Figure S15:**
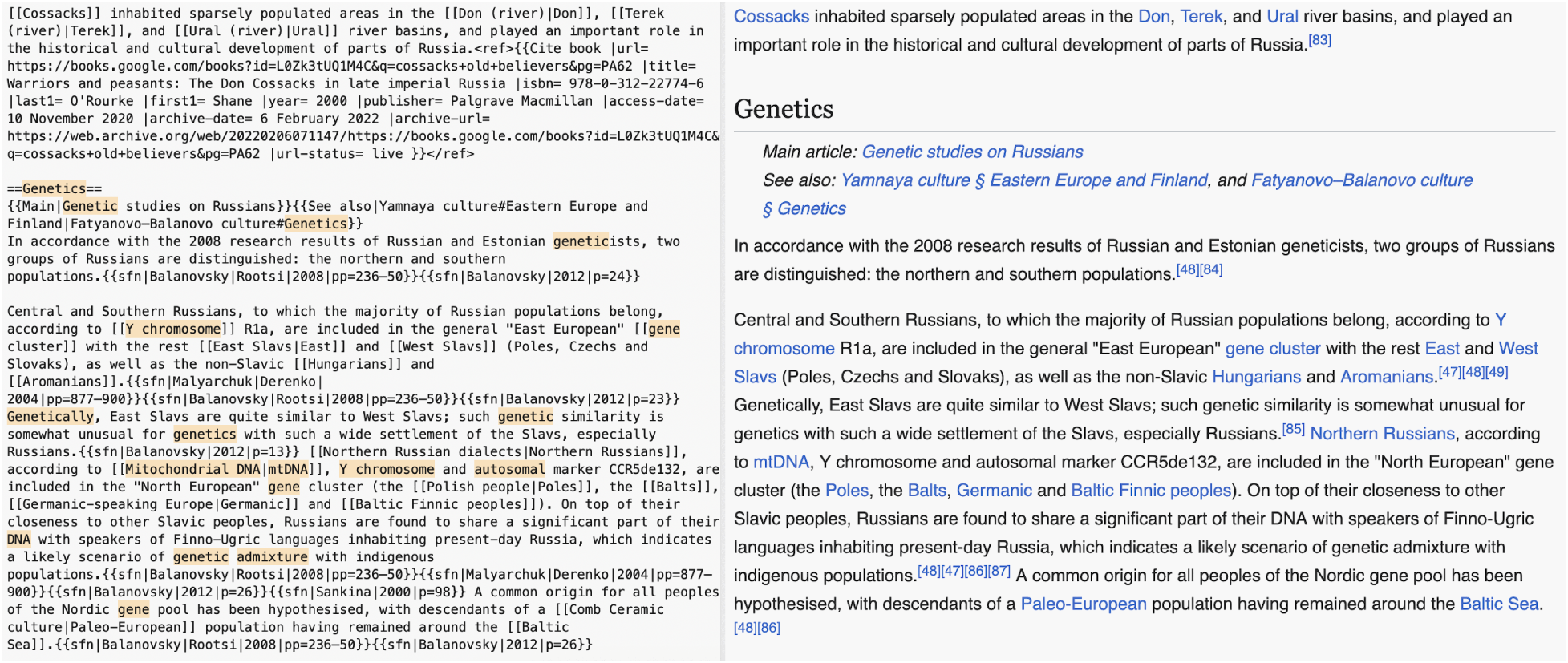
An example of Wikipedia markup. (Left) A sample of Wikipedia markup from the page “Russians”. Genetics keywords have been highlighted. The markup code ==Genetics== delineates a section header in the page. (Right) How the text is rendered in a web browser.

**Figure S16:**
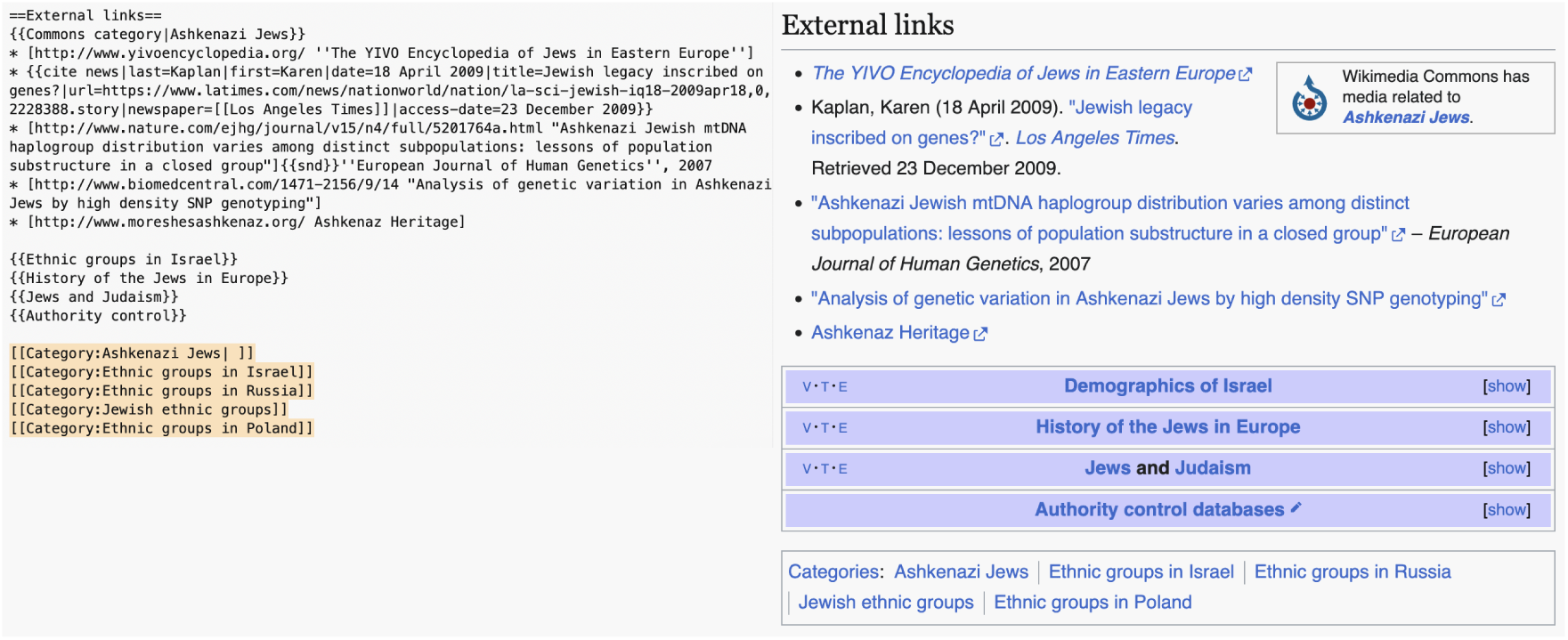
Adding specialized markup code to a Wikipedia page places that page into a category. (Left) Markup code from the “External links” section of the page “Ashkenazi Jews”. At the bottom, code used to add categories is highlighted. This code is of the format [[Category:NAME]], where “NAME” is the name of the category. This page belongs to five categories: “Ashkenazi Jews”, “Ethnic groups in Israel”, “Ethnic groups in Russia”, “Jewish ethnic groups”, and “Ethnic groups in Poland”. (Right) The section generated from the markup text, as viewed in a browser; the categories are visible at the bottom of the page.

**Figure S17:**
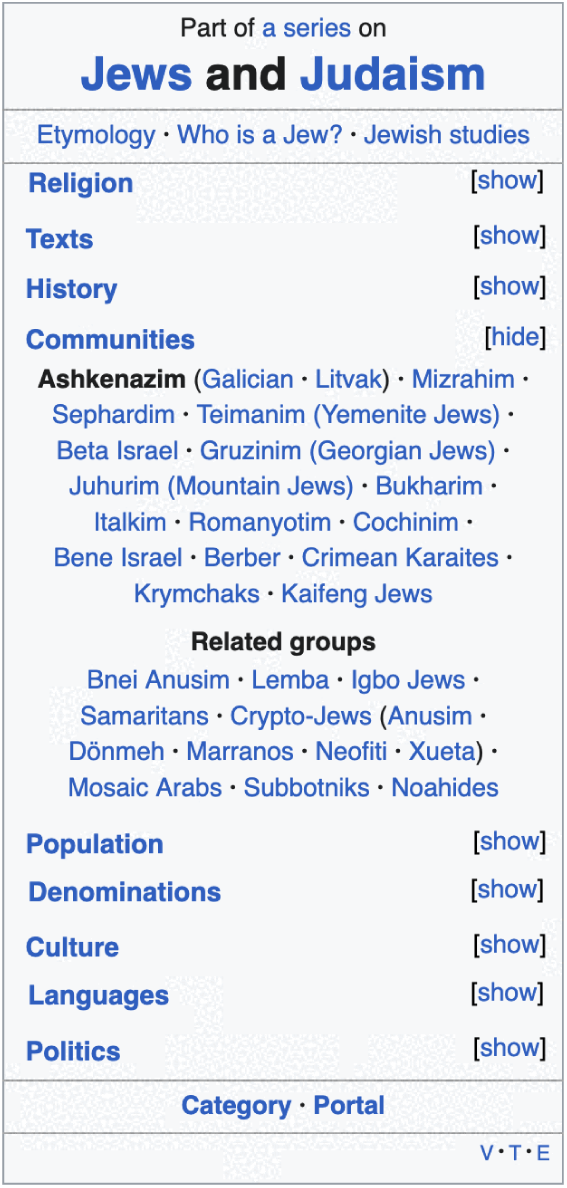
Navigation boxes are used to provide links to related topics. This navigation box (titled “Jews and Judaism sidebar”) is present in the “Ashkenazi Jews” page, with the default view presented.

**Table S2:** On the 137 Wikipedia pages about nationalities, genetics research is most often used in a biological context. Of 137 nationalities that had Wikipedia pages, 93 (67.8%) had genetics keywords. To determine the context in which a genetic keyword appears, we identified the sections of a Wikipedia page that contained genetics keywords and then assigned a context based on the section’s title (outlined in the supplementary material). The most common context in which genetics keywords appeared on 137 demonym pages was in a section of the page dedicated to biology or biological studies (e.g. a “Genetics” or “Genetic studies”) section. The next most common were the origin or formation of a group (e.g. “Ethnogenesis” or “Origins”) or its culture (e.g. “History” or “Language”). Keywords could be used in multiple contexts, e.g. a “History” section could have a “Genetic origins” sub-section, which would count towards both “Culture” and “Biology”.

| Context | Pages |
| --- | --- |
| Biology | 72 (52.6%) |
| Origins | 37 (27.0%) |
| Culture | 33 (24.1%) |
| Introduction | 26 (19.0%) |
| Reference | 23 (16.8%) |
| Race, ethnicity, nationality, and citizenship | 20 (14.6%) |
| Other | 6 (4.4%) |

**Table S3:** Demonym pages with genetics figures.

| Page name |
| --- |
| Dominicans |
| Maltese_people |
| Palestinians |
| Filipinos |
| Dutch_people |
| Mexicans |
| Macedonians_(ethnic_group) |
| Estonians |
| Ukrainians |
| Czechs |
| Kazakhs |
| Greeks |
| Finns |
| Kyrgyz_people |
| Bulgarians |
| Romanians |
| Lithuanians |
| Italians |
| Mongols |
| Tajiks |
| Ethiopians |
| Micronesians |
| Egyptians |
| Hungarians |
| Portuguese_people |

**Table S4:** The vast majority of citations within genetics sections are only used in one page. This data corresponds to figure S9. We extracted 1,195 unique DOIs from citations within genetics sections; the table lists, for each number of pages a DOI appears in, the number of DOIs and their proportion of the total.

| Number of pages | DOIs | Proportion of total |
| --- | --- | --- |
| 1 | 835 | 0.6987 |
| 2 | 191 | 0.1598 |
| 3 | 78 | 0.0653 |
| 4 | 46 | 0.0385 |
| 5 | 22 | 0.0184 |
| 6 | 4 | 0.0033 |
| 7 | 8 | 0.0067 |
| 8 | 4 | 0.0033 |
| 9 | 3 | 0.0025 |
| 10 | 1 | 0.0008 |
| 13 | 1 | 0.0008 |
| 17 | 1 | 0.0008 |
| 18 | 1 | 0.0008 |

**Table S5:** All references in genetics sections in the corpus appearing in at least ten pages.

| Citation | Count |
| --- | --- |
| Kushniarevich et al. (2015). Genetic Heritage of the Balto-Slavic Speaking Populations: A Synthesis of Autosomal, Mitochondrial and Y-Chromosomal Data. *PLOS ONE*. | 18 |
| Tambets et al. (2018). Genes reveal traces of common recent demographic history for most of the Uralic-speaking populations. *Genome Biology*. | 17 |
| Tishkoff et al. (2009). The Genetic Structure and History of Africans and African Americans. *Science*. | 13 |
| Hassan et al. (2008). Y-chromosome variation among Sudanese: Restricted gene flow, concordance with language, geography, and history. *American Journal of Physical Anthropology*. | 10 |

**Table S6:** Most cited journals in genetics sections in the corpus.

| Journal | Citations |
| --- | --- |
| The American Journal of Human Genetics | 76 |
| European Journal of Human Genetics | 62 |
| American Journal of Physical Anthropology | 56 |
| Scientific Reports | 53 |
| PLoS ONE | 44 |
| Nature | 41 |
| Molecular Biology and Evolution | 37 |
| Annals of Human Genetics | 35 |
| Proceedings of the National Academy of Sciences | 34 |
| Human Immunology | 32 |
| Science | 32 |
| Nature Communications | 28 |
| Journal of Human Genetics | 27 |
| Current Biology | 22 |
| Forensic Science International: Genetics | 21 |
| Human Genetics | 21 |
| PLOS ONE | 20 |
| PLoS Genetics | 19 |
| Human Biology | 16 |
| Annals of Human Biology | 16 |
| Genes | 14 |
| BMC Evolutionary Biology | 14 |
| American Journal of Human Biology | 13 |
| Frontiers in Genetics | 13 |
| Cell | 12 |

**Table S7:** The top ten Wikipedia pages ranked by number of talk page discussions with genetics keywords present. Every Wikipedia page has an associated “talk page”, split into sections corresponding to topics of discussion. Users may use these talk pages to discuss page contents. We extracted the section from every discussion of every talk page and identified any discussions that contained genetics keywords. This table lists the ten pages with the highest number of discussions that contai<u>ned a genetics keyword.</u>

| Page title | Discussions with<br>genetics keywords | Total<br>discussions | Percent with<br>genetics keywords |
| --- | --- | --- | --- |
| Black people | 272 | 919 | 29.6 |
| White people | 265 | 881 | 30.1 |
| Palestinians | 172 | 601 | 28.6 |
| Ashkenazi Jews | 138 | 411 | 33.6 |
| English people | 130 | 330 | 30.4 |
| Spaniards | 113 | 252 | 44.8 |
| Jews | 102 | 894 | 11.4 |
| Turkish people | 94 | 372 | 25.3 |
| African Americans | 89 | 750 | 11.9 |
| Kurds | 85 | 572 | 14.9 |

**Table S8:** Number of plain text paragraphs with genetics keywords in their markup, as of December 31 of each year (f<u>rom corpus page text).</u>

| Year | Paragraphs with keywords | Paragraphs without keywords |
| --- | --- | --- |
| 2002 | 6 | 396 |
| 2003 | 17 | 759 |
| 2004 | 65 | 2097 |
| 2005 | 270 | 6920 |
| 2006 | 467 | 13631 |
| 2007 | 582 | 19970 |
| 2008 | 736 | 22745 |
| 2009 | 861 | 30623 |
| 2010 | 971 | 33890 |
| 2011 | 1126 | 37270 |
| 2012 | 1281 | 40019 |
| 2013 | 1430 | 34979 |
| 2014 | 1483 | 38056 |
| 2015 | 1634 | 40781 |
| 2016 | 1755 | 43632 |
| 2017 | 1903 | 47191 |
| 2018 | 1985 | 49399 |
| 2019 | 2125 | 51792 |
| 2020 | 2288 | 57571 |
| 2021 | 2372 | 63054 |
| 2022 | 2531 | 66986 |
| 2023 | 2655 | 72303 |
| 2024 | 2889 | 78795 |
| 2025 | 3100 | 83253 |
| Total | 34532 | 936112 |

**Table S9:** Count and proportion of corpus discussions with genetics keywords mentioning a topic. Counts of the number of discussions from corpus talk pages that contain genetics keywords as well as keywords related to a given topic (see Methods—Text analysis and Methods—Text topic analysis for full details). The proportion given is the proportion of discussions with genetics keywords and also any topic keyword (5,806).

| Topic | Discussions | Proportion |
| --- | --- | --- |
| Ancestry & origins | 4415 | 0.76042 |
| Identity & ethnicity | 4327 | 0.74526 |
| Language | 3616 | 0.62280 |
| Group comparison | 3290 | 0.56665 |
| Admixture | 2713 | 0.46727 |
| Race | 2630 | 0.45297 |
| Religion | 2445 | 0.42111 |
| Popgen methods | 2270 | 0.39097 |
| Social structure | 2147 | 0.36978 |
| Phenotypes | 1000 | 0.17223 |
| Statistics & clustering | 521 | 0.08973 |
| Medical genetics | 433 | 0.07457 |
| Subsistence | 346 | 0.05959 |
| Archaic | 195 | 0.03358 |
| Hereditarianism | 154 | 0.02652 |

**Table S10:** Count and proportion of corpus talk pages with genetics keywords mentioning a topic. Counts of the number of corpus talk pages that have at least one discussion with a genetics keyword as well as keywords related to a given topic (see Methods—Text analysis and Methods— Text topic analysis for full details). The proportion given is the proportion of corpus talk pages with genetics keywords and also any topic keyword (649).

| Topic | Pages | Proportion |
| --- | --- | --- |
| Ancestry & origins | 535 | 0.8243451 |
| Identity & ethnicity | 530 | 0.8166410 |
| Language | 480 | 0.7395994 |
| Group comparison | 439 | 0.6764253 |
| Admixture | 402 | 0.6194145 |
| Social structure | 385 | 0.5932203 |
| Race | 364 | 0.5608629 |
| Popgen methods | 363 | 0.5593220 |
| Religion | 359 | 0.5531587 |
| Phenotypes | 204 | 0.3143297 |
| Statistics & clustering | 148 | 0.2280431 |
| Medical genetics | 145 | 0.2234206 |
| Subsistence | 136 | 0.2095532 |
| Archaic | 90 | 0.1386749 |
| Hereditarianism | 46 | 0.0708783 |

**Table S11:** Section lengths and proportions of the “Ashkenazi Jews” Wikipedia page. Section length measures the number of characters (including spaces) of plain text; citations do not count towards <u>length.</u>

| Level | Section | Length | % Length |
| --- | --- | --- | --- |
| 0 | Introduction | 3,345 | 4.70 |
| 2 | Etymology | 3,611 | 5.08 |
| 2 | History | 26,102 | 36.70 |
| 3 | Early Jewish communities in Europe | 2,054 | 2.89 |
| 3 | Jewish–Roman Wars | 2,826 | 3.97 |
| 3 | Late antiquity | 4,718 | 6.63 |
| 3 | Early Middle Ages | 1,405 | 1.98 |
| 3 | High and Late Middle Ages migrations | 6,593 | 9.27 |
| 3 | Modern history | 7,704 | 10.83 |
| 4 | The Holocaust | 1,589 | 2.23 |
| 4 | Israel | 2,186 | 3.07 |
| 4 | United States of America | 638 | 0.90 |
| 2 | Definition | 10,291 | 14.47 |
| 3 | By religion | 3,241 | 4.56 |
| 3 | By culture | 4,757 | 6.69 |
| 3 | By ethnicity | 788 | 1.11 |
| 3 | Eastern vs. Western/Central Ashkenazim | 1,475 | 2.07 |
| 2 | Customs, laws, and traditions | 2,804 | 3.94 |
| 3 | Ashkenazic liturgy | 520 | 0.73 |
| 2 | Relations with Sephardim | 877 | 1.23 |
| 2 | Notable Ashkenazim | 321 | 0.45 |
| 2 | Genetics | 22,201 | 31.22 |
| 3 | Genetic origins | 18,217 | 25.61 |
| 4 | Male lineages: Y-chromosomal DNA | 2,332 | 3.28 |
| 4 | Female lineages: Mitochondrial DNA | 3,069 | 4.32 |
| 4 | Association and linkage studies (autosomal DNA) | 9,488 | 13.34 |
| 3 | The Khazar hypothesis | 1,850 | 2.60 |
| 3 | East Asian ancestry | 337 | 0.47 |
| 3 | Medical genetics | 1,759 | 2.47 |
| 2 | See also | 40 | 0.06 |
| 2 | Explanatory notes | 24 | 0.03 |
| 2 | References | 1,137 | 1.60 |
| 3 | Citations | 18 | 0.03 |
| 3 | Sources | 22 | 0.03 |
| 3 | References for “Who is an Ashkenazi Jew?” | 61 | 0.01 |
| 3 | Other references | 1,020 | 1.43 |
| 2 | External links | 368 | 0.52 |

**Table S12:** Sections of the page “Ashkenazi Jews”, with position, character count, and estimated internal link clicks (December 2025 data). Section length measures the number of characters (including spaces) of plain text; references do not count towards length. Out clicks counts an estimated number of clicks on links within a section (see Methods—Case studies).

| Index | Section title | Length | Out clicks |
| --- | --- | --- | --- |
| 0 | Introduction | 3364 | 37720 |
| 1 | Etymology | 3611 | 4166 |
| 2 | History | 26102 | 14081 |
| 3 | Definition | 10291 | 12134 |
| 4 | Customs, laws, and traditions | 2788 | 239 |
| 5 | Relations with Sephardim | 820 | 781 |
| 6 | Notable Ashkenazim | 321 | 2511 |
| 7 | Genetics | 22201 | 5863 |
| 8 | See also | 40 | 0 |
| 9 | Explanatory notes | 24 | 0 |
| 10 | References | 1137 | 0 |
| 11 | External links | 368 | 0 |

**Table S13:** Top terms by log odds ratio for Grokipedia and Wikipedia (Page:“Ashkenazi Jews”).

| Rank | Site | Term | Log odds | Site | Term | Log odds |
| --- | --- | --- | --- | --- | --- | --- |
| 1 | Grokipedia | iq | 3.785 | Wikipedia | people | 3.287 |
| 2 | Grokipedia | amid | 3.614 | Wikipedia | number | 3.287 |
| 3 | Grokipedia | selection | 3.407 | Wikipedia | time | 3.196 |
| 4 | Grokipedia | verbal | 3.148 | Wikipedia | ashkenazic | 3.095 |
| 5 | Grokipedia | overrepresentation | 3.070 | Wikipedia | different | 3.095 |
| 6 | Grokipedia | cognitive | 3.070 | Wikipedia | result | 2.859 |
| 7 | Grokipedia | bottlenecks | 3.070 | Wikipedia | authors | 2.815 |
| 8 | Grokipedia | exceeding | 2.987 | Wikipedia | italian | 2.715 |
| 9 | Grokipedia | analyses | 2.896 | Wikipedia | parties | 2.715 |
| 10 | Grokipedia | restrictions | 2.896 | Wikipedia | french | 2.715 |
| 11 | Grokipedia | evidenced | 2.896 | Wikipedia | druze | 2.715 |
| 12 | Grokipedia | approximately | 2.832 | Wikipedia | present | 2.548 |
| 13 | Grokipedia | effective | 2.796 | Wikipedia | called | 2.548 |
| 14 | Grokipedia | pattern | 2.796 | Wikipedia | syrian | 2.548 |
| 15 | Grokipedia | allegations | 2.684 | Wikipedia | area | 2.548 |
| 16 | Grokipedia | subsequent | 2.684 | Wikipedia | lebanese | 2.548 |
| 17 | Grokipedia | moneylending | 2.684 | Wikipedia | revolts | 2.548 |
| 18 | Grokipedia | points | 2.684 | Wikipedia | yiddishkeit | 2.548 |
| 19 | Grokipedia | fostering | 2.684 | Wikipedia | persecution | 2.548 |
| 20 | Grokipedia | intelligence | 2.684 | Wikipedia | favor | 2.548 |
| 21 | Grokipedia | norms | 2.684 | Wikipedia | style | 2.548 |
| 22 | Grokipedia | empirical | 2.684 | Wikipedia | settle | 2.548 |
| 23 | Grokipedia | shift | 2.684 | Wikipedia | according | 2.548 |
| 24 | Grokipedia | driven | 2.684 | Wikipedia | sense | 2.548 |
| 25 | Grokipedia | ongoing | 2.684 | Wikipedia | particular | 2.548 |

**Table S14:** Top terms by log odds ratio for Grokipedia and Wikipedia (Page: “Black people”).

| Rank | Site | Term | Log odds | Site | Term | Log odds |
| --- | --- | --- | --- | --- | --- | --- |
| 1 | Grokipedia | rates | 4.147 | Wikipedia | aboriginal | 3.781 |
| 2 | Grokipedia | sub | 3.459 | Wikipedia | government | 3.709 |
| 3 | Grokipedia | despite | 3.352 | Wikipedia | arabs | 3.632 |
| 4 | Grokipedia | lower | 3.254 | Wikipedia | blackness | 3.457 |
| 5 | Grokipedia | analyses | 3.201 | Wikipedia | person | 3.457 |
| 6 | Grokipedia | crime | 3.145 | Wikipedia | israel | 3.357 |
| 7 | Grokipedia | amid | 3.145 | Wikipedia | press | 3.357 |
| 8 | Grokipedia | reflecting | 3.086 | Wikipedia | considered | 3.357 |
| 9 | Grokipedia | iq | 3.024 | Wikipedia | citizens | 3.121 |
| 10 | Grokipedia | exhibit | 3.024 | Wikipedia | various | 3.121 |
| 11 | Grokipedia | disparities | 3.024 | Wikipedia | coloured | 3.121 |
| 12 | Grokipedia | environmental | 2.957 | Wikipedia | brought | 3.121 |
| 13 | Grokipedia | variants | 2.957 | Wikipedia | shipped | 2.977 |
| 14 | Grokipedia | parent | 2.885 | Wikipedia | drop | 2.977 |
| 15 | Grokipedia | diverse | 2.808 | Wikipedia | persons | 2.977 |
| 16 | Grokipedia | pre | 2.808 | Wikipedia | end | 2.977 |
| 17 | Grokipedia | empirical | 2.808 | Wikipedia | accepted | 2.977 |
| 18 | Grokipedia | elevated | 2.808 | Wikipedia | peninsula | 2.977 |
| 19 | Grokipedia | educational | 2.725 | Wikipedia | said | 2.977 |
| 20 | Grokipedia | structure | 2.725 | Wikipedia | different | 2.977 |
| 21 | Grokipedia | scores | 2.725 | Wikipedia | definition | 2.977 |
| 22 | Grokipedia | exceeding | 2.634 | Wikipedia | activists | 2.977 |
| 23 | Grokipedia | versus | 2.634 | Wikipedia | charlotte | 2.810 |
| 24 | Grokipedia | samples | 2.634 | Wikipedia | especially | 2.810 |
| 25 | Grokipedia | remain | 2.634 | Wikipedia | mainly | 2.810 |

**Table S15:** Top terms by log odds ratio for Grokipedia and Wikipedia (Page: “White people”).

| Rank | Site | Term | Log odds | Site | Term | Log odds |
| --- | --- | --- | --- | --- | --- | --- |
| 1 | Grokipedia | steppe | 3.496 | Wikipedia | nineteenth | 3.825 |
| 2 | Grokipedia | amid | 3.449 | Wikipedia | percent | 3.766 |
| 3 | Grokipedia | hunter | 3.346 | Wikipedia | colombia | 3.636 |
| 4 | Grokipedia | migrations | 3.290 | Wikipedia | twentieth | 3.488 |
| 5 | Grokipedia | neolithic | 3.290 | Wikipedia | area | 3.313 |
| 6 | Grokipedia | global | 3.231 | Wikipedia | according | 3.215 |
| 7 | Grokipedia | empirical | 3.231 | Wikipedia | chilean | 3.213 |
| 8 | Grokipedia | analyses | 3.102 | Wikipedia | different | 3.213 |
| 9 | Grokipedia | bronze | 3.030 | Wikipedia | described | 3.102 |
| 10 | Grokipedia | clusters | 3.030 | Wikipedia | seventeenth | 3.102 |
| 11 | Grokipedia | ago | 2.953 | Wikipedia | blumenbach | 3.102 |
| 12 | Grokipedia | driven | 2.953 | Wikipedia | yellow | 3.102 |
| 13 | Grokipedia | enabling | 2.953 | Wikipedia | community | 3.102 |
| 14 | Grokipedia | low | 2.870 | Wikipedia | colombian | 2.976 |
| 15 | Grokipedia | broader | 2.870 | Wikipedia | cuban | 2.976 |
| 16 | Grokipedia | continental | 2.870 | Wikipedia | question | 2.976 |
| 17 | Grokipedia | shifts | 2.870 | Wikipedia | december | 2.833 |
| 18 | Grokipedia | selection | 2.870 | Wikipedia | chileans | 2.833 |
| 19 | Grokipedia | emphasizing | 2.779 | Wikipedia | island | 2.833 |
| 20 | Grokipedia | labor | 2.779 | Wikipedia | mexicans | 2.833 |
| 21 | Grokipedia | urban | 2.779 | Wikipedia | result | 2.833 |
| 22 | Grokipedia | iron | 2.779 | Wikipedia | retrieved | 2.833 |
| 23 | Grokipedia | limited | 2.779 | Wikipedia | suriname | 2.833 |
| 24 | Grokipedia | frequencies | 2.779 | Wikipedia | use | 2.833 |
| 25 | Grokipedia | continuity | 2.779 | Wikipedia | september | 2.833 |

**Table S16:**
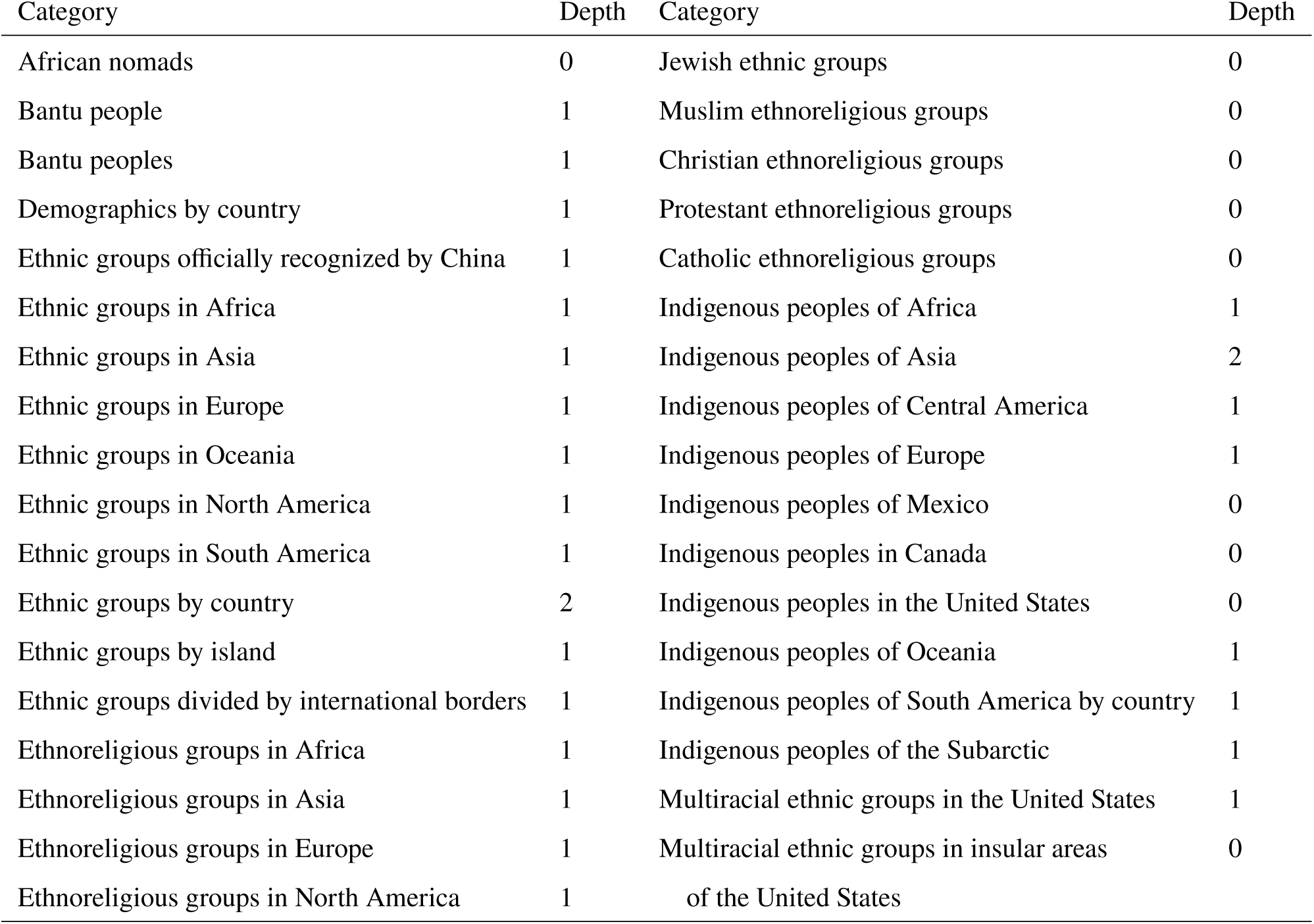
Categories used for data collection. The Wikipedia categories (and sub-categories, scraped to the noted depth) from which corpus pages were drawn. Since a page may belong to multiple categories, the total number of pages analyzed is smaller than the sum across categories.

**Table S17:** Logit Regression Results—Corpus pages.

| Model Information |  |  |  |  |  |  |
| --- | --- | --- | --- | --- | --- | --- |
| Dependent Variable | keyword_binary |  |  |  |  |  |
| No. Observations | 6538 |  |  |  |  |  |
| Model | Logit |  |  |  |  |  |
| Df Residuals | 6536 |  |  |  |  |  |
| Method | MLE |  |  |  |  |  |
| Df Model | 1 |  |  |  |  |  |
| Pseudo $R^2$ | 0.2951 | | | | | |
| Log-Likelihood | −1934.9 |  |  |  |  |  |
| Converged | True |  |  |  |  |  |
| LL-Null | −2745.0 |  |  |  |  |  |
| Covariance Type | Nonrobust |  |  |  |  |  |
| LLR $p$-value | 0.000 | | | | | |

| | Coef. | Std. Err. | $z$ | $P > \vert z\vert $ | [0.025 | 0.975] |
| --- | --- | --- | --- | --- | --- | --- |
| Intercept | −5.3842 | 0.133 | −40.514 | 0.000 | −5.645 | −5.124 |
| log_daily_views_plus_one | 0.9493 | 0.029 | 32.901 | 0.000 | 0.893 | 1.006 |

**Table S18:** Logit Regression Results—Talk pages.

| Model Information |  |  |  |  |  |  |
| --- | --- | --- | --- | --- | --- | --- |
| Dependent Variable | keyword_binary |  |  |  |  |  |
| No. Observations | 6538 |  |  |  |  |  |
| Model | Logit |  |  |  |  |  |
| Df Residuals | 6536 |  |  |  |  |  |
| Method | MLE |  |  |  |  |  |
| Df Model | 1 |  |  |  |  |  |
| Pseudo $R^2$ | 0.4571 | | | | | |
| Log-Likelihood | −1148.1 |  |  |  |  |  |
| Converged | True |  |  |  |  |  |
| LL-Null | −2114.8 |  |  |  |  |  |
| Covariance Type | Nonrobust |  |  |  |  |  |
| LLR $p$-value | 0.000 | | | | | |

| | Coef. | Std. Err. | $z$ | $P > \vert z\vert $ | [0.025 | 0.975] |
| --- | --- | --- | --- | --- | --- | --- |
| Intercept | −8.2410 | 0.242 | −34.112 | 0.000 | −8.714 | −7.767 |
| log_daily_views_plus_one | 1.4010 | 0.046 | 30.151 | 0.000 | 1.310 | 1.492 |

**Table S19:** The number of pages in the corpus created every year, with a cumulative count.

| Year | Pages created | Cumulative | Year | Pages created | Cumulative |
| --- | --- | --- | --- | --- | --- |
| 2001 | 47 | 47 | 2014 | 179 | 5097 |
| 2002 | 137 | 184 | 2015 | 169 | 5266 |
| 2003 | 197 | 381 | 2016 | 122 | 5388 |
| 2004 | 439 | 820 | 2017 | 151 | 5539 |
| 2005 | 653 | 1473 | 2018 | 78 | 5617 |
| 2006 | 830 | 2303 | 2019 | 89 | 5706 |
| 2007 | 530 | 2833 | 2020 | 109 | 5815 |
| 2008 | 535 | 3368 | 2021 | 99 | 5914 |
| 2009 | 403 | 3771 | 2022 | 132 | 6046 |
| 2010 | 330 | 4101 | 2023 | 144 | 6190 |
| 2011 | 345 | 4446 | 2024 | 114 | 6304 |
| 2012 | 235 | 4681 | 2025 | 237 | 6541 |
| 2013 | 237 | 4918 |  |  |  |

**Table S20:** Topics in paragraphs of text that contain genetics keywords, as of December 31, 2025. Topics are identified using a secondary set of keywords. This analysis was run on plain text where the underlying markup contained genetics keywords (see Methods—Text topic analysis for details).

| Topic | Pages | Proportion (corpus) | Proportion (pages w/ keywords) |
| --- | --- | --- | --- |
| Ancestry & origins | 630 | 0.0963 | 0.7135 |
| Popgen terminology | 500 | 0.0765 | 0.5663 |
| Identity & Ethnicity | 492 | 0.0752 | 0.5572 |
| Language | 461 | 0.0705 | 0.5221 |
| Admixture | 406 | 0.0621 | 0.4598 |
| Group comparisons | 404 | 0.0618 | 0.4575 |
| Social structure | 164 | 0.0251 | 0.1857 |
| Religion | 155 | 0.0237 | 0.1755 |
| Race | 153 | 0.0234 | 0.1733 |
| Subsistence | 138 | 0.0211 | 0.1563 |
| Statistics & clustering | 126 | 0.0193 | 0.1427 |
| Medical genetics | 87 | 0.0133 | 0.0985 |
| Phenotypes | 53 | 0.0081 | 0.0600 |
| Archaic | 39 | 0.0060 | 0.0442 |
| Hereditarianism | 1 | 0.0002 | 0.0011 |

**Table S21:** Changes in topics of corpus text over 20 years. Comparing the proportion of corpus pages that have genetics keywords in paragraphs discussing different topics. We compare snapshots of corpus pages from December 31, 2025 to December 31, 2005. The largest absolute increases in proportion were in text using terms from population genetics methods, discussing language or linguistics, or subsistence mode. The largest decreases in proportion were in text using terms related to medical genetics, religion, and race.

| Topic | Proportion (2005) | Proportion (2025) | Difference |
| --- | --- | --- | --- |
| Popgen terminology | 0.4245 | 0.5663 | 0.1417 |
| Language | 0.4057 | 0.5221 | 0.1164 |
| Subsistence | 0.0566 | 0.1563 | 0.0997 |
| Statistics & clustering | 0.0472 | 0.1427 | 0.0955 |
| Ancestry & origins | 0.6226 | 0.7135 | 0.0908 |
| Group comparisons | 0.3962 | 0.4575 | 0.0613 |
| Admixture | 0.4245 | 0.4598 | 0.0353 |
| Archaic | 0.0094 | 0.0442 | 0.0347 |
| Identity & Ethnicity | 0.5377 | 0.5572 | 0.0195 |
| Social structure | 0.1698 | 0.1857 | 0.0159 |
| Hereditarianism | 0.0189 | 0.0011 | -0.0177 |
| Phenotypes | 0.0943 | 0.0600 | -0.0343 |
| Medical genetics | 0.1415 | 0.0985 | -0.0430 |
| Religion | 0.2547 | 0.1755 | -0.0792 |
| Race | 0.2925 | 0.1733 | -0.1192 |

## Notes

https://doi.org/10.5281/zenodo.21537405

https://github.com/diazale/wikigenetics

### Competing Interest Statement

The authors have declared no competing interest.

